# Pentose Sugars Encode Sequence-Dependent DNA-RNA Segregation for Programmable Engineering of Multiphase Condensates

**DOI:** 10.1101/2025.06.23.656378

**Authors:** Wei Guo, Feipeng Chen, Andrew B. Kinghorn, Xiufeng Li, Haohui Ou, Yi Pan, Rong Luo, Yuchao Wang, Kwan Kiu Lau, Tianjiao Mao, Fang Wang, Zhenyu Yang, Xiaolai Li, Ying Chen, Sihan Liu, Yage Zhang, Yang Song, Tiantian Kong, Xiangze Zeng, Ho Cheung Shum

**Affiliations:** Guangdong Key Laboratory of Biomedical Measurements and Ultrasound Imaging, School of Biomedical Engineering, Shenzhen University Medical School, Shenzhen University, Shenzhen 518060, China; Department of Mechanical Engineering, Faculty of Engineering, The University of Hong Kong, Hong Kong (SAR), Hong Kong, China; Advanced Biomedical Instrumentation Centre, Hong Kong Science Park, Shatin, New Territories, Hong Kong (SAR), Hong Kong, China; School of Biomedical Sciences, LKS Faculty of Medicine, The University of Hong Kong, Hong Kong (SAR), Hong Kong, China; College of Food Science and Technology, Nanjing Agricultural University, Nanjing, 210095, Jiangsu, China; Institute of Biomedical Engineering, College of Medicine, Southwest Jiaotong University, Chengdu, 610031, P.R.China; Department of Ophthalmology, LKS Faculty of Medicine, The University of Hong Kong, Hong Kong (SAR), Hong Kong, China; State Key Laboratory of Metal Matrix Composites, School of Material Science and Engineering, Shanghai Jiao Tong University, Shanghai 200240, China; Department of Physics, Hong Kong Baptist University, Kowloon Tong, Hong Kong, China; Department of Chemistry and Department of Biomedical Engineering, City University of Hong Kong, Hong Kong SAR, China

## Abstract

The nucleus segregates DNA and RNA with high precision to safeguard genome stability and gene regulation despite their homologous structures, yet the underlying molecular principles have long evaded both understanding and synthetic translation. Here we show that a single atomic difference between the pentose sugars, a 2’-OH in RNA versus a 2’-H in DNA, suffices to drive their phase segregation with cationic peptides. This subtle chemical distinction makes RNA bind peptides more strongly than sequence-identical DNA, generating an asymmetry that, when modulated by sequence-encoded homotypic interactions, drives the formation of core-shell multiphase condensates. Using a supervised machine-learning approach, we distill these molecular principles into a predictive design rule and harness it to program a library of oligonucleotide pairs, termed SEGREGamers, that self-assemble into droplets with coexisting DNA- and RNA-rich domains. These synthetic condensates recapitulate nuclear compartment functions, including selective molecular partitioning and enhanced RNA catalysis. Our results establish a chemically encoded platform for engineering synthetic nuclear mimics and programmable biomolecular condensates, and suggest that sugar identity may have served as an ancient physical mechanism for organizing nucleic acids.

## Introduction

In the cell nucleus, DNA and RNA are meticulously organized into distinct compartments at high spatial resolution^1, 2^, a segregation essential for genome stability and gene regulation. Recreating such precise spatial segregation between DNA and RNA in synthetic compartments has remained a fundamental challenge, yet his capability is essential for synthetic cells to enable spatial gene circuitry^3, 4^ and for gene delivery vehicles such as CRISPR co-delivery systems to prevent premature mixing of co-encapsulated DNA and RNA^5^. Biomolecular condensates formed by phase separation have emerged as a key mechanism underlying intracellular DNA and RNA compartmentalization^2, 6–8^. However, owing to their homologous chemical structures^9^, DNA and RNA tend to mix entropically^10^, and the molecular principles that drive their high-precision segregation in cells remain undeciphered and untapped for synthetic systems.

Synthetic condensates can recapitulate complex multiphase architectures that segregatively enrich distinct biopolymer species^11–13^. In principle, multiphase condensates can be encoded through molecular complexity, such as sequence heterogeneity^14, 15^ and competitive interactions^16–21^. For nucleic acids, phase separation is typically modulated through three known handles: electrostatic complexation via the phosphate backbone^17, 22, 23^, base pairing^24–28^ and its associated higher-order structures^29–31^. The pentose sugar, however, has been largely overlooked. Polymer physics predicts that phase behavior is exquisitely sensitive to subtle molecular differences^32, 33^, raising the possibility that the minimal chemical distinction between DNA and RNA, a single 2’-OH group in ribose versus a 2’-H in deoxyribose, could itself encode their spatial segregation and thereby serve as a new, orthogonal thermodynamic handle for engineering synthetic multiphase compartments. This possibility, however, has remained largely unexplored for nearly a century since Phoebus Levene first distinguished ribose from deoxyribose^34^.

Here we demonstrate that this single atomic difference suffices to drive sequence-dependent phase segregation of DNA and RNA when mixed with cationic peptides. Using systematic experiments, theory, and all-atom molecular dynamics simulations, we show that RNA binds peptides more strongly than its sequence-identical DNA counterpart. This asymmetry, tunable by specific sequence features, governs the balance between homotypic and heterotypic interactions, dictating whether the system assembles into single-phase or core-shell multiphase condensates with DNA-rich shells and RNA-rich cores. By integrating a data-driven, machine-learning-augmented approach^35, 36^, we distill the sequence features that control multiphase assembly into a predictive design rule. This rule enables the *de novo* design of a library of DNA and RNA oligonucleotide pairs, termed SEGREGamers, which spontaneously form multiphase droplets with programmable core-shell morphologies and material properties, spanning an estimated 10^5^ to 10^9^ distinct variants. These synthetic condensates recapitulate key nuclear compartment functions, including selective small-molecule partitioning, RNA aptamer folding, and enhanced DNAzyme-mediated RNA cleavage. Our findings establish that the long-overlooked chemical dissimilarity between DNA and RNA actively encodes phase behavior, providing a physical basis for nucleic acid compartmentalization^6^. By linking sugar identity to sequence-encoded design rules for multiphase architectures, our work establishes a new paradigm for condensate engineering^37^, opening avenues for the formulation of next-generation gene-editing delivery vehicles^38^, synthetic protocell design^39^, and cell-free biomanufacturing platforms^40^.

## Results and Discussion

### Multiphase droplets can be formed by sequence-identical DNA and RNA oligonucleotides

Nuclear transcriptional condensates enrich both DNA and RNA, recruiting proteins for biogenesis^2, 6^ (**Fig. 1A**). In addition, some DNA-binding transcription factors have been reported to exhibit affinity for RNA^41^, while certain RNA-binding proteins can also interact with DNA counterparts^42, 43^. These observations suggest the presence of competitive DNA-and RNA-protein interactions (**Fig. 1B** and **Supplementary Fig. 1**), even when DNA and RNA have identical sequences (**Fig. 1C**). To explore whether these cooperative interactions could facilitate DNA-RNA segregation, we employed DNA and RNA oligonucleotides in combination with poly-L-lysine (PLL) as model systems. We selected the RNA oligonucleotide [UUAGAA]_4_ (TERRA_M2), derived from the telomeric repeat-containing RNA (TERRA, [UUAGGG]_n_), which contributes to telomeric heterochromatin formation. The two successive G-to-A mutations inhibit non-canonical G-quadruplex structures and promote droplet formation when complexed with PLL^29^. An intriguing observation emerged when studying oligonucleotide-PLL complex coacervation. Both [UUAGAA]_4_ and its DNA counterpart, [TTAGAA]_4_, formed droplets with PLL (**Fig. 1D**). Remarkably, when mixing [UUAGAA]_4_, [TTAGAA]_4_, and PLL together, core-shell multiphase droplets formed, with DNA localizing to the shell and RNA enriched in the core (**Fig. 1D**).

**Fig. 1.**
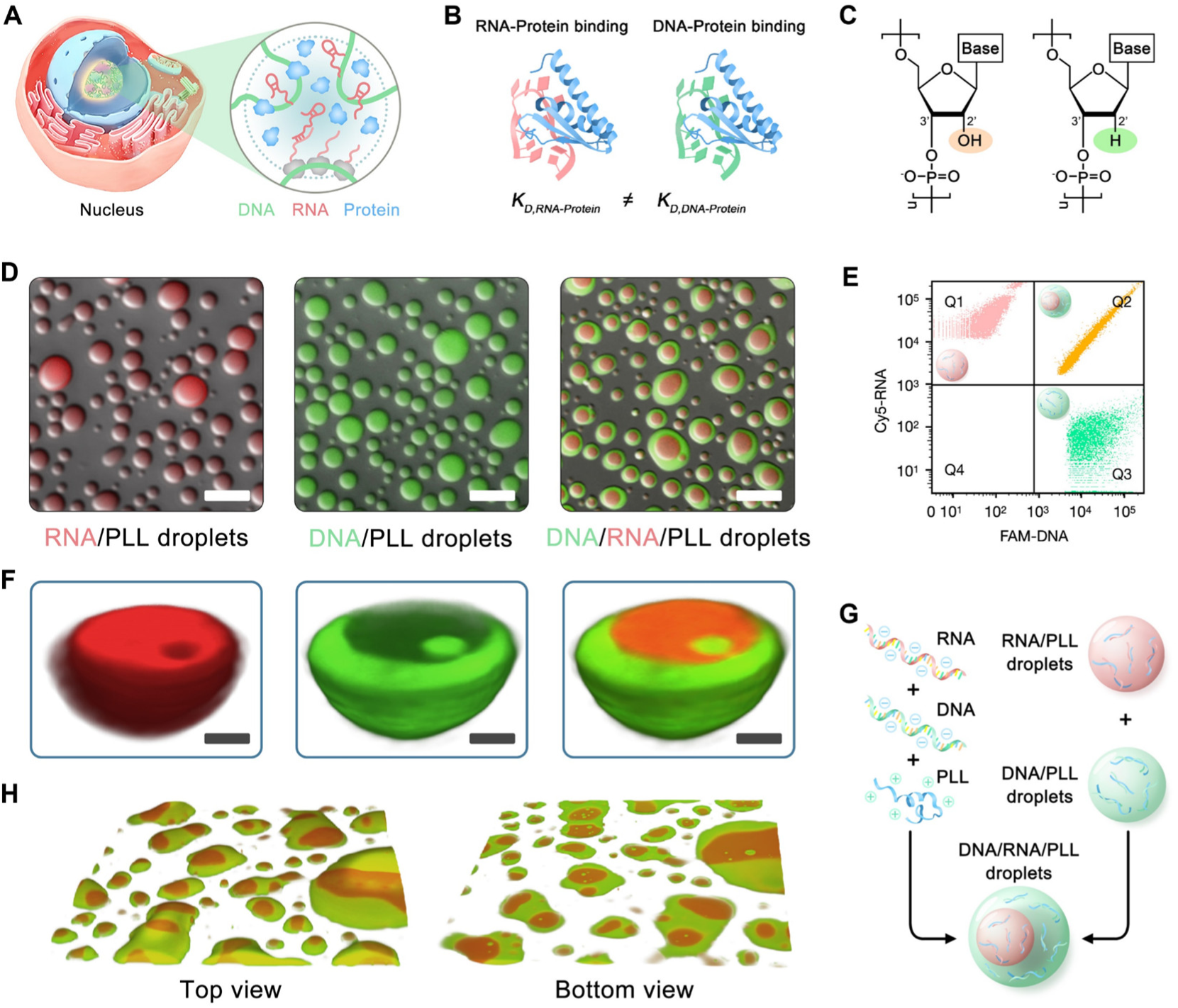
Single-stranded DNA and RNA oligonucleotides with the identical sequence form multiphase droplets. (**A**) Schematic of nuclear compartments within nucleus. (**B**) Competitive binding of proteins between DNA and RNA, where *K* is the association constant. (**C**) Molecular structure of RNA (red) and DNA (green). (**D**) Droplets formed by RNA-PLL, DNA-PLL and DNA-RNA-PLL phase separation. Droplets were prepared at the charge concentration of [e+] = [e-] = 5 mM in 500 mM KCl buffer. FAM-DNA and Cy5-RNA were used at 2 μM. For DNA-RNA-PLL phase separation, the [DNA]: [RNA] ratio was 1:1. Scale bar, 10 μm. (**E**) FACS fluorescence-gated analysis of the three types of droplets in (C). (**F**) Stacked confocal microscopy images of a multiphase DNA/RNA/PLL droplet. Scale bar, 2 μm. (**G**) Two equivalent pathways to form multiphase DNA/RNA/PLL droplets. (**H**) Glass wetting of multiphase DNA/RNA/PLL droplets. An observation box of 88.3 × 88.3 × 3.8 μm^3^ is shown.

Quantitative fluorescence-activated cell sorting (FACS) analysis revealed that both fluorescently-labelled DNA and RNA were sequestered into droplets (**Fig. 1E**). Within multiphase droplets, DNA and RNA separated, forming a DNA-rich phase and an RNA-rich phase (**Fig. 1F** and **Supplementary Fig. 2**, A and B), suggesting segregation between DNA and RNA within droplets. This observed phase-specific partitioning of fluorescently labeled components is a hallmark of multiphase organization within biomolecular condensates^23^. For all experiments unless otherwise specified, condensates were formed by combining a pre-mixed solution of DNA and RNA with an addition of PLL solution. To determine if the final condensate architecture was influenced by kinetic trapping, we systematically varied the order of component addition. Reassuringly, all mixing pathways-including the addition of PLL to pre-formed DNA-PLL or RNA-PLL condensates (**Fig. 1G** and **Supplementary Fig. 2C** and **2D**)-consistently yielded the core-shell structure. This robustness indicates that the multiphase organization represents the thermodynamic minimum for the system, suggesting limited miscibility between DNA-PLL droplets and RNA-PLL droplets^14^. Importantly, mixing DNA, RNA, and PLL triggers the rapid assembly of multiphase condensates within minutes, and the resultant core-shell architectures persist stably for several hours (**Supplementary Fig. 2E**, **2F** and **Supplementary Video 1**). Additionally, these core-shell structured droplets displayed liquid-like properties, demonstrated by their ability to wet a glass slide (**Fig. 1H** and **Supplementary Fig. 3A**) and fuse with one another (**Supplementary Fig. 3B**). The liquid-like properties of both shell and core phases were further confirmed by fluorescence recovery after photobleaching (FRAP) measurements (**Supplementary Fig. 3C**).

**Fig. 2.**
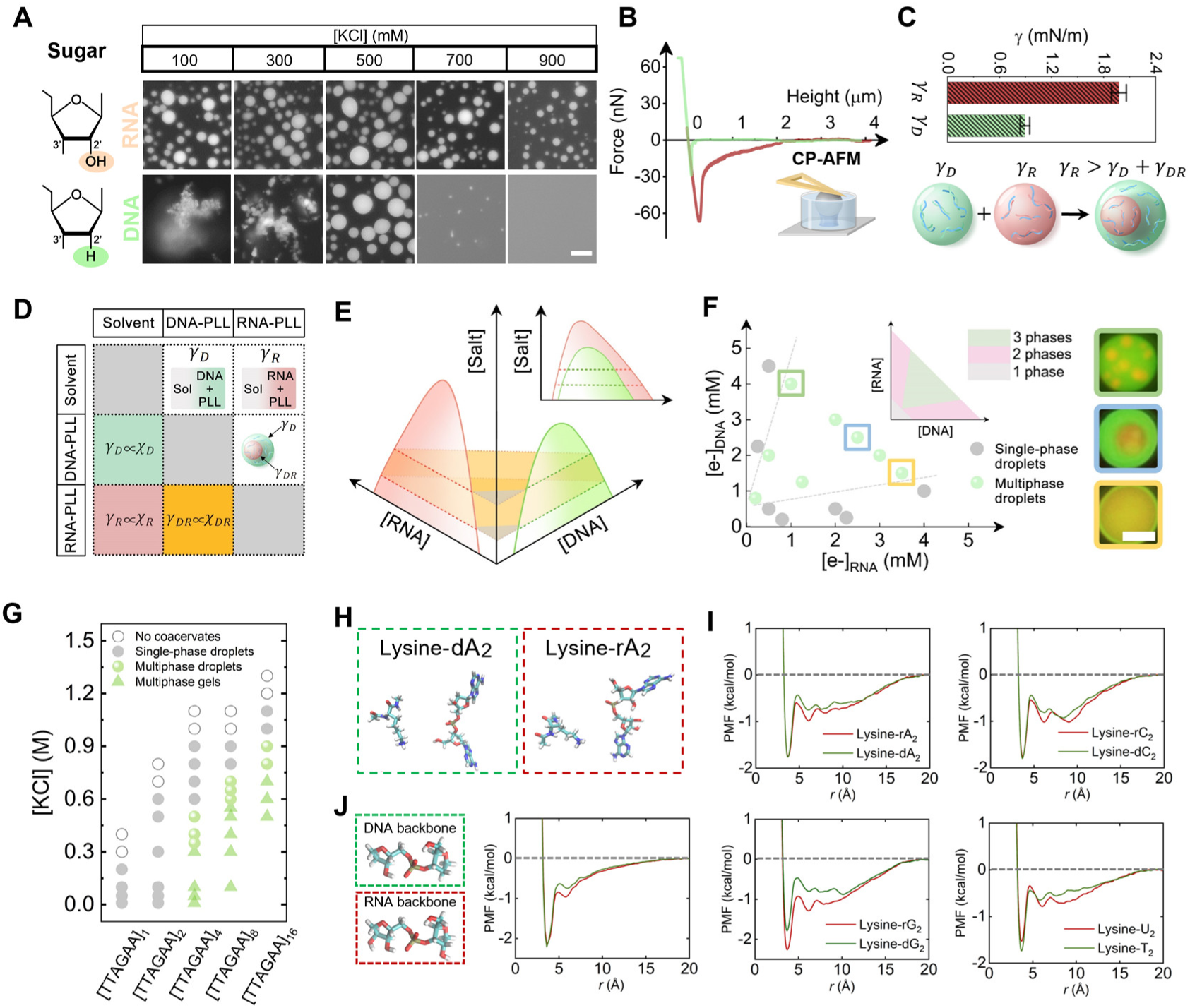
Pentose sugars encode distinct DNA-PLL and RNA-PLL interactions to drive multiphase droplet formation. (**A**) Fluorescent images of RNA-and DNA-PLL condensates at different KCl concentrations. Droplets were formed at the charge concentration of [e+] = [e-] = 5 mM. Scale bar, 10 μm. (**B**) Force-distance curves of DNA-PLL droplets (green) and RNA-PLL droplets (red) measured by colloidal probe atomic force microscopy (CP-AFM) for coacervate rheology measurement. (**C**) DNA-PLL droplets have lower interfacial tension than that of RNA-PLL droplets, enabling the engulfment of RNA droplets by DNA droplets. Error bars represent SEM (standard error of the mean) from five individual measurements. **D**) The interaction matrix for DNA/PLL, RNA/PLL, and DNA/RNA/PLL droplet formation. The correlation between interfacial tension γ and interaction strength χ, γ∝χ, indicates stronger interaction in RNA-PLL phase separation. (**E**) Three-dimensional phase diagram of DNA-PLL and RNA-PLL complex coacervation. Droplets containing both DNA and RNA formed at the light-yellow colored domain. Insert: Classical phase diagram of DNA-and RNA-PLL complex coacervation. (**F**) Phase diagram of DNA and RNA concentration-dependent multiphase droplet formation, with representative microscope images shown on the right. Scale bar, 5 μm. Insert: Phase diagram of PLL-scaffolded DNA-RNA segregation. (**G**) Chain length-dependent multiphase droplet formation across a range of salt concentrations. (**H**) All-atom molecular dynamics simulations of the interactions between lysine and dimerized deoxynucleotides or ribonucleotides. (**I**) Potential of mean force (PMF) profiles quantifying the interactions between lysine and each of the four (deoxy) nucleotide dimers. (**J**) All-atom simulations and corresponding PMF analyses of lysine interacting specifically with the sugar-phosphate backbones of deoxynucleotides and ribonucleotides. Unless otherwise noted, only DNA sequences are shown.

### Distinct DNA-and RNA-PLL interaction strength results in multicomponent phase separation involving DNA-RNA segregation

Interestingly, DNA-PLL complex coacervation formed solid-like aggregates at lower salt concentrations (100 and 300 mM KCl), while RNA-PLL complexation consistently produced liquid-like droplets (**Fig. 2A**). Gel-like aggregates can result from nucleic acid intra-/intermolecular hybridization^25, 44^, increased charge density^44^, and higher chain rigidity^45^, as confirmed via DNA hybridization in our experimental systems (**Supplementary Fig. 4**). Notably, both circular dichroism (CD) spectroscopy (**Supplementary Fig. 5**) and computational secondary structure predictions (**Supplementary Fig. 6**) indicated that neither the DNA nor the RNA sequence adopted stable secondary structures. Despite having identical sequences and similar charge densities, the formation of DNA/PLL aggregates and RNA/PLL droplets at lower salt concentrations was attributed to distinct interactions with PLL. The enhanced hybridization propensity of DNA, as well as the possible formation of non-Watson-Crick structures, both aided by the electrostatic neutralization provided by PLL, may contribute to this aggregation^44^. Importantly, even at lower salt concentrations (100 and 300 mM KCl), mixing DNA, RNA, and PLL resulted in separate DNA-rich and RNA-rich domains within aggregates (**Supplementary Fig. 7**, **A** and **B**), although both domains exhibited a lower fluorescence recovery level in FRAP compared to multiphase droplets (**Supplementary Fig. 7C**). Consequently, increasing salt concentration via evaporation triggered the solid-to-liquid phase transition of multiphase aggregates (**Supplementary Fig. 7D** and **Supplementary Video 2**). This evaporation assay, which concentrates salt and biomolecules through water loss (**Supplementary Fig. 8**), revealed distinct phase behaviors across a range of final concentrations, and consistent with non-evaporative control experiments using pre-set initial salt and biomolecule concentrations which also yielded multiphase condensates (**Supplementary Fig. 9**). Interestingly, adding water back into the evaporating sample results in re-occurrence of multiphase condensates (**Supplementary Video 3**). Taken together, our data show that oligonucleotides [UUAGAA]_4_ and [TTAGAA]_4,_ albeit with the identical sequence, confer phase immiscibility to DNA/RNA/PLL condensates at both liquid-and gel-like states.

To elucidate the mechanisms governing the formation of multiphase droplets, we characterized droplet rheology and interfacial tensions by colloidal probe atomic force microscopy^29, 46^ (CP-AFM, **Supplementary Text 1**). The force-distance curves revealed that RNA/PLL droplets exhibit greater attraction forces compared to DNA/PLL droplets (**Fig. 2B** and **Supplementary Fig. 10**). Notably, the substantial difference in interfacial tension between DNA/PLL droplets (γ*_D_* ≈ 0.89 mN/m) and RNA/PLL droplets (γ*_R_* ≈ 1.98 mN/m) (**Fig. 2C** and **Supplementary Text 1**) points to the complete engulfment of RNA/PLL droplets by DNA/PLL droplets, which is crucial for the formation of core-shell structures^47^ (**Fig. 2C**). Using an electrophoretic mobility shift assay (EMSA), we compared the binding affinities of PLL for DNA (dTERRA_M2) and RNA (rTERRA_M2). The measured dissociation constant for the RNA-PLL complex (7.4 nM) was approximately fivefold lower than that for the DNA-PLL complex (34.0 nM), indicating a stronger binding affinity for RNA, a finding consistent with previous studies (**Supplementary Fig. 11**)^48^ ^41^. Moreover, DNA/PLL droplets dissolved at 700 mM KCl, while RNA/PLL droplets remained stable even at 900 mM KCl (**Fig. 2A**). These data suggested that RNA-PLL interactions are stronger than those for DNA. Importantly, the stronger RNA-PLL interactions are also exhibited within multiphase droplets, as demonstrated by the higher salt resistance (**Supplementary Fig. 12A**) of the RNA-rich core phase, together with the more concentrated PLL distribution inside (**Supplementary Fig. 12B**). The distinct DNA-PLL and RNA-PLL interactions are also exhibited by the partitioning of FAM-DNA and Cy5-RNA into RNA-PLL and DNA-PLL droplets, respectively, with the latter display a more significant partitioning effect (**Supplementary Fig. 13**). The stronger RNA-PLL interactions predominate in the RNA-rich core phase, consistent with previous observations that polyelectrolyte pairs with stronger interactions tend to become the dominant components within the core phase^13, 14, 16^.

The multiphase droplet formation indicates a three-phase coexistence region on the DNA-RNA-PLL phase diagram. To investigate how distinct RNA-PLL and DNA-PLL interaction strengths shape their phase diagram, we considered DNA-and RNA-PLL complexes at an equal charge ratio ([e+]:[e-] = 1:1). The competing effects of solvent-polymer complex and polymer complex-polymer complex interactions are summarized in **Fig. 2D**. Since interfacial tension is directly proportional to the Flory parameter*χ*^8, 47, 49, 50^(**Supplementary Text 2**), it follows that *χ_R_* > *χ_D_* Accordingly, a higher-dimensional multicomponent phase diagram was derived^51^. DNA-PLL complexes and RNA-PLL complexes condensed into a polymer-rich phase under salt conditions where both DNA-PLL and RNA-PLL undergo phase separation (**Fig. 2E**, **Supplementary Fig. 14** and **Supplementary Text 3**). At a fixed salt concentration, the system has two-phase regions along the DNA and RNA axis, and a three-phase coexistence region in the middle^52^ (insert of **Fig. 2F**).

To validate the phase diagram, we varied DNA and RNA concentrations while maintaining the charge ratio of 1:1. We observed that multiphase droplets formed when both DNA and RNA concentrations reached a critical value, while single-phase droplets formed if either concentration was below the critical value (**Fig. 2F**, **Supplementary Fig. 15A and B**). The interface between the core and shell, along with the RNA/DNA stoichiometry-dependent core/shell size within multiphase droplets (**Fig. 2F**), suggest that DNA-PLL and RNA-PLL complexes exhibit segregative interactions, with *χ_DR_* ∝ γ*_DR_*. Such segregative interaction has been widely featured in all-aqueous two-phase systems (ATPS)^53^. Moreover, we found that, akin to ATPS, multiphase droplet formation is chain length dependent. Increasing the chain length of DNA and RNA enhanced segregative propensity between them, as indicated by the expanded salt concentration range for forming multiphase coacervates (**Fig. 2G**, **Supplementary Fig. 15C**-**F**). In contrast, no multiphase coacervates were observed for shorter-chain oligonucleotides, such as [UUAGAA]_1_ and [UUAGAA]_2_. These results demonstrate that increasing the chain length of DNA and RNA leads to more pronounced and stable phase segregation within condensates. This observation is consistent with recent studies showing that enhanced oligomerization of intrinsically disordered proteins results in increased immiscibility and multiphasic organization^12^. Together, our findings confirm that segregative interactions between DNA and RNA oligonucleotides drive multiphase droplet formation.

To test whether DNA-RNA segregation could be facilitated by other cationic polymers, we replaced PLL by other cationic polymers and found that Poly-D-lysine (PDL), Polydiallyldimethylammonium chloride (PDADMAC), Polyethyleneimine (PEI), and Poly-L-arginine (PLA) could also form multiphase droplets (**Supplementary Fig. 16A**). Notably, the salt conditions and DNA:RNA ratios varied for different cationic polymers due to the distinct charge densities and lengths of the individual polycations^16^. Additionally, we demonstrated that multiphase droplets formed when KCl was replaced by NaCl (**Supplementary Fig. 16B**). Besides long-chain polycations, we also tested R10 and K10, which were able to form multiphase condensates with DNA and RNA (**Supplementary Fig. 17**). However, when PLL was replaced with FUS (Fused in Sarcoma) or SUP-K72, no apparent multiphase condensates were observed (**Supplementary Fig. 18**), presumably due to their reduced charge density, which limits the interaction strength difference between DNA and RNA required for multiphase condensate formation. These findings suggest that DNA-RNA segregation is a universal phenomenon induced by cationic agents with high charge density, and does not rely on specific polycation-dependent molecular interactions^16^.

### Differences in the sugar ring of oligonucleotides impact their interactions with PLL

Distinct interactions of sequence-identical DNA and RNA with poly-L-lysine (PLL) are primarily attributed to differences in their sugar components. Specifically, DNA contains deoxyribose sugar, which lacks the 2’-OH group present in the ribose sugar of RNA. To further investigate the impact of sugar structures on the formation of nucleic acid-peptide condensates, we examined RNA oligonucleotides with 2’-O-methylation. These modified RNA oligonucleotides formed solid-like structures across a range of KCl concentrations from 100 to 700 mM (**Supplementary Fig. 19**), presumably due to the increased hydrophobicity^54, 55^. This observation indicates that sugar modifications significantly influence nucleic acid interactions with PLL and, consequently, the formation of condensates. Our findings are consistent with recent studies demonstrating that 2’-O-methylation alters percolation transition of RNA^56^.

To explore the impact of the deoxyribose sugar in DNA and the 2’-OH group in the ribose sugar of RNA on their interactions with PLL at the atomistic level, we utilized all-atom molecular dynamics simulations. We selected dimers of deoxyribonucleotides (e.g., dA2) and ribonucleotides (e.g., rA2) to represent DNA and RNA, respectively, and constructed a PLL monomer to interact with these nucleotide dimers (**Fig. 2H**). Each ribonucleotide dimer contains two additional 2’-OH groups compared to each deoxyribonucleotide dimer (**Fig. 2H**). Detailed simulation setups are described in the Materials and Methods section.

We assessed the potential of mean force (PMF) as a function of the intermolecular distance for different interacting sidechain pairs. Our results demonstrate that, for all four nucleobases, the PMF between lysine and ribonucleotide dimers is significantly stronger than that of deoxyribonucleotide dimers. This is consistent with experimental findings showing stronger RNA-PLL interactions compared to DNA-PLL interactions (**Fig. 2I**). Notably, the differences in PMF between lysine-ribonucleotide and lysine-deoxyribonucleotide dimers are nucleobase-dependent, with the maximum difference observed for rG2/dG2 and the minimum for U2/T2 (**Fig. 2I** and **Supplementary Fig. 20**). These findings suggest that, in addition to electrostatic interactions between the NH^3+^ of lysine and the PO^4-^ of (deoxyribo)nucleotides, lysine-(deoxyribo)nucleotide dimer interactions are also mediated by nucleobases, potentially due to hydrogen bonds, cation-π, and π-π interactions. For example, urea-induced suppression of multiphase condensate formation (**Supplementary Fig. 21**) indicates the critical role of hydrogen bonding in this process. To further investigate the differences arising from sugars, we examined interactions between lysine and the DNA/RNA backbone, excluding nucleobases to isolate sugar effects (**Fig. 2J**). Stronger interactions were confirmed between the RNA backbone and lysine compared to the DNA backbone, as indicated by a deeper PMF for lysine-RNA backbone interactions (**Fig. 2J**). Overall, these simulation results confirm that the distinct RNA-and DNA-PLL interactions are fundamentally governed by differences in their sugar components.

### Sequence-dependent assembly of DNA/RNA multiphase droplets

Our all-atom molecular simulations demonstrated that nucleotide bases modulate interactions between nucleic acids and lysine residues, with guanine exhibiting the most substantial differences in interaction strength. These findings indicate that the DNA-and RNA-PLL interactions underlying multiphase condensate formation are sequence-dependent. To elucidate the molecular determinants underlying the sequence dependence of DNA/RNA multiphase condensate formation, we investigated the role of nucleic base patterning by generating three distinct variants of TERRA_M2 (TERRA_M2_Mut1, TERRA_M2_Mut2, and TERRA_M2_Mut3). Importantly, these sequences were designed to preserve the overall nucleotide composition of TERRA_M2 while eliminating potential secondary structures. Strikingly, two of the three DNA/RNA pairs were capable of forming multiphase condensates comparable to those formed by the native TERRA_M2 sequence (**Fig. 3A**), while TERRA_M2_Mut3 formed single-phase condensates. These results indicate that the specific sequence arrangement within TERRA_M2 plays a role in DNA-RNA segregation within condensates.

**Fig. 3.**
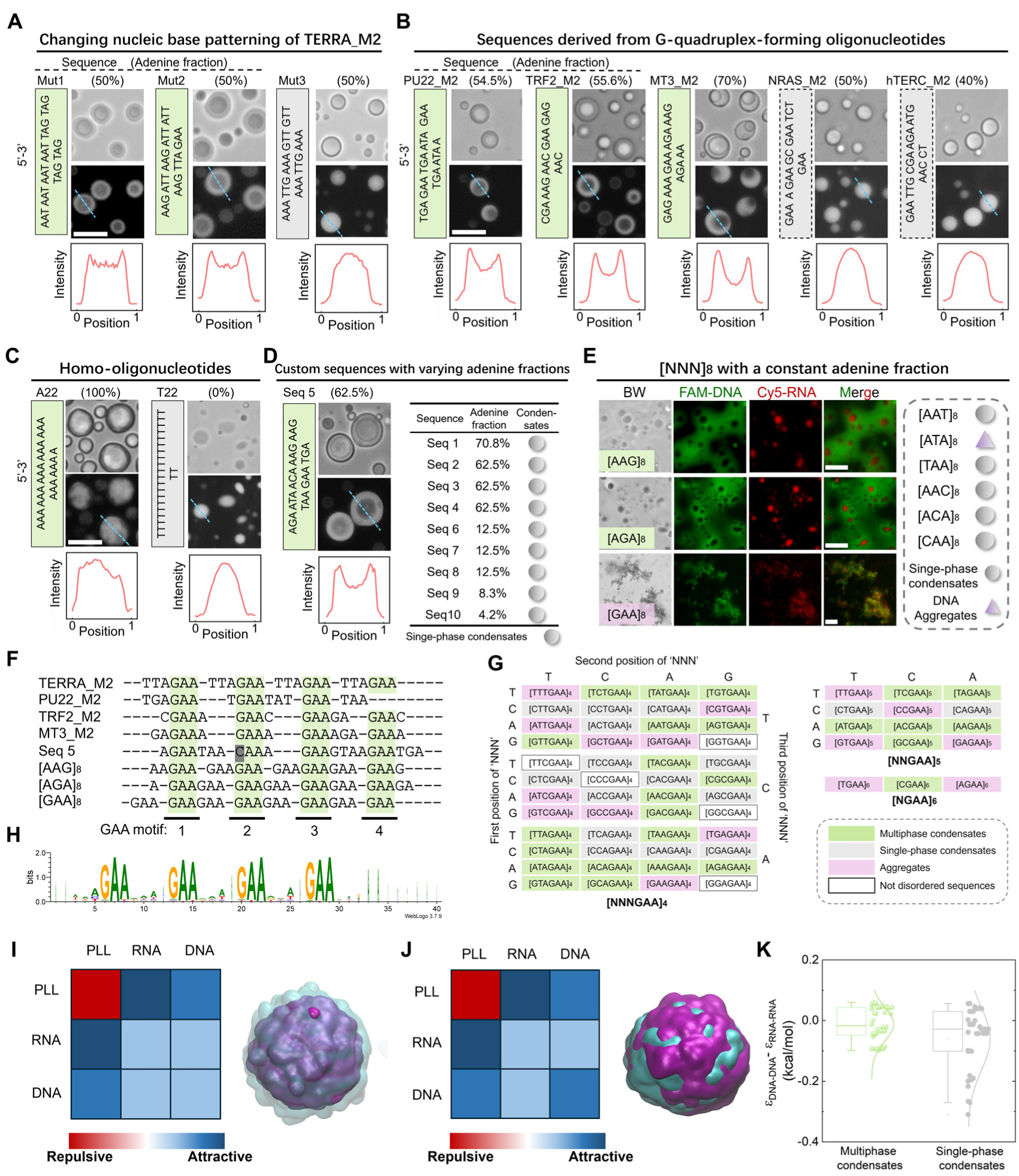
Sequence-dependent propensity of oligonucleotides for DNA/RNA/PLL multiphase condensate formation. (**A**) Varying nucleic base patterning of TERRA_M2 modulated multiphase condensate formation. Scale bar, 10 μm. (**B**) Motifs of PU22_M2, TRF2_M2 and MT3_M2 formed multiphase DNA/RNA/PLL droplets while motifs of NARS_M2 and hTERC_M2 did not. Scale bar, 10 μm. (**C**) Homo-oligonucleotides of A_22_ formed multi-phase condensates while T_22_ (U_22_) formed single-phase condensates. Scale bar, 10 μm. (**D**) A custom designed motif Seq 5, formed multiphase droplets. Scale bar, 10 μm. **(E)** For disordered trinucleotide expansions that contain two adenines in each repeating units, only those contain guanines formed DNA/RNA/PLL multiphase condensates. (**F**) Colour-coded alignment of sequences capable of forming multiphase condensates revealed the recurrent presence of multiple ‘GAA’ motifs. (**G**) GAA-rich oligonucleotides with repeating units [NNNGAA]_4_, [NNGAA]_5_ and [NGAA]_6_ exhibited sequence-dependent propensity for forming DNA/RNA/PLL multiphase condensates. (**H**) A consensus sequence motif was identified among all tested oligonucleotides that promote multiphase condensate formation. (**I**) Coarsen grained simulation of DNA/RNA/PLL multiphase condensate formation via the comparable DNA-DNA and RNA-RNA interactions. (**J**) Single-phase droplets were formed when DNA-DNA interaction strength exceeds that of RNA-RNA interaction. (**K**) Oligonucleotides capable of forming multiphase condensates exhibited comparable RNA-RNA and DNA–DNA interactions compared to those forming single-phase condensates. Unless otherwise noted, only DNA sequences are shown.

Then we explored other G-quadruplex-forming oligonucleotides with mutations similar to those in TERRA_M2, including PU22_M2, MT3_M2, TRF2_M2, NARS_M2, and hTERC_M2 (**Supplementary Table 1**). We observed multiphase droplets in PU22_M2, MT3_M2, and TRF2_M2 samples, whereas NARS_M2 and hTERC_M2 predominantly formed single-phase droplets (**Fig. 3B**). Notably, across all tested DNA/RNA oligonucleotides, RNA exhibited stronger interactions with PLL than DNA, reflected by higher critical salt concentrations required for droplet formation in both multiphase and single-phase systems (**Supplementary Table 2**). Sequence analysis revealed that TERRA_M2, PU22_M2, MT3_M2, and TRF2_M2 contained higher adenine content compared to NARS_M2 and hTERC_M2 (**Supplementary Table 1**). Moreover, complex coacervation of PLL with homo-oligonucleotides of dA_n_/rA_n_ (n=22, 40) yielded inhomogeneous droplets, a phenomenon absent for T_n_/U_n_ sequences (n=22, 40) (**Fig. 3C**, **Supplementary Fig. 22A**). In addition, N-to-A substitutions of TERRA_M2 and PU22_M2 at non-guanine sites formed multiphase droplets, while N-to-T/U substitutions did not (**Supplementary Fig. 22B**). More importantly, CD spectroscopy (**Supplementary Fig. 5**) and computational secondary structure predictions (**Supplementary Fig. 6**) indicated that none of the sequences adopted stable secondary structures, pointing to a sequence-dependent but structure-independent mechanism governing multiphase condensate formation.

These results suggest that DNA-RNA segregation within condensates is promoted by adenine and suppressed by thymine/uracil. To test whether higher adenine content specifically enhances multiphase formation, we designed ten synthetic sequences with varying adenine fractions (Seq 1 highest A fraction, Seq 10 lowest; **Supplementary Table 3**). Surprisingly, only Seq 5 formed multiphase droplets (**Fig. 3D**, **Supplementary Fig. 23**). Consistently, our CP-AFM measurements of interfacial tension reveal that, for oligonucleotides such as PU22_M2, MT3_M2, and Seq5, the RNA-PLL condensates exhibit significantly higher interfacial tension compared to their DNA-PLL counterparts. In contrast, for hTERC_M2 and Seq3, the interfacial tension values for RNA-PLL and DNA-PLL condensates are similar (**Supplementary Fig. 24**), even Seq 3 has a high adenine fraction. These results indicate that sequence-specific features beyond mere adenine content influence phase behavior.

Next, we examined sequences with triplet repeats [NNN]_8_, where two of the three nucleotides were fixed as adenine (A) to promote DNA-RNA segregation. Among the nine tested pairs, only [AAG]_8_ and [AGA]_8_ formed well-defined core-shell multiphase droplets, whereas [GAA]_8_ formed gel-like, immiscible DNA-and RNA-rich domains (**Fig. 3E**). The remaining six sequences failed to produce multiphase condensates. These observations suggest that the presence of guanine within sequences containing high adenine content is critical for multiphase assembly, consistent with our atomistic findings that guanine encodes the strongest differences in DNA-and RNA-PLL interactions (**Fig. 2I**, **Supplementary Fig. 20**).

These results confirm that although an increased adenine fraction is correlated with a higher propensity for multiphase condensate formation, it is not a definitive predictor. To further dissect the contributions of sequence composition, we systematically analyzed additional parameters, including the fractions of cytosine (C), guanine (G), and pyrimidines (U and T). However, none of these compositional metrics established a clear threshold that could reliably distinguish sequences capable of forming multiphase condensates from those that cannot (**Supplementary Fig. 25**). Collectively, these findings indicate that the ability of DNA/RNA oligonucleotides to drive multiphase condensate formation cannot be fully explained by overall sequence content alone. This suggests that other aspects of sequence information-such as the presence of specific short sequence motifs or local sequence patterns-may play a critical role.

### ‘GAA’-rich sequences exhibit a strong propensity to drive the formation of multiphase condensates

Sequence alignment of these eight multiphase-forming oligonucleotides, including TERRA_M2, PU22_M2, MT3_M2, TRF2_M2, Seq 5, [AAG]_8_, [AGA]_8_, and [GAA]_8,_ revealed they all contain multiple “GAA” motifs (**Fig. 3F**), implicating these motifs in facilitating phase separation. Supporting this, a mutant with “GAA” motifs (PU22_M1) failed to form multiphase droplets despite a higher guanine content (**Supplementary Fig. 26**). To further validate this hypothesis, we tested sequences associated with repeat expansion disorders, including [NNNGAA]_4_, [NGAA]_6_, and [NNGAA]_5_, where N is any nucleotide (A, C, G, T/U). To minimize confounding effects of secondary structure, we excluded sequences predicted to form stable G-quadruplexes or i-motifs. Among 58 sequences tested, 26 formed core-shell condensates with RNA-rich cores and DNA-rich shells, 16 formed solid-like aggregates, and 16 remained as single-phase condensates (**Fig. 3G** and **Supplementary Table 9**). These findings reinforce that GAA-rich sequences possess an intrinsic propensity to generate multiphase condensates with segregated DNA and RNA domains. Notably, a group of sequences composed exclusively of G and A residues formed multiphase condensates (**Supplementary Table 4**), indicating that the propensity for Watson-Crick base pairing is not correlated with the ability to form multiphase condensates.

Sequence alignment and comprehensive analysis of all tested oligonucleotides revealed a strong correlation between the presence of “GAA” motifs and multiphase condensate formation (**Fig. 3H** and **Supplementary Fig. 27**). Oligonucleotides capable of forming multiphase condensates consistently display a higher fraction of “GAA” content, with an enrichment of guanine residues, compared to those that do not (**Supplementary Fig. 28**). Notably, these sequences also feature A-rich segments-serving as spacers-between “GAA” motifs, in contrast to the more variable, non-A-rich gaps found in non-multiphase-forming sequences (**Supplementary Fig. 28**). These sequence-dependent properties of oligonucleotides capable of forming multiphase condensates suggest that multiphase condensate formation is governed by specific sequence-mediated interactions.

In ternary systems comprising DNA, RNA, and PLL, the emergence of multiphase condensates can be attributed to a finely tuned balance between heterotypic interactions and homotypic interactions^57^. Furthermore, our results for the critical salt concentrations, as presented in **Supplementary Table 2** and **5**, indicate that while a substantial difference in critical salt concentration between RNA-PLL and DNA-PLL coacervates appears to be a necessary condition for the formation of multiphase condensates, it is not sufficient by itself. Specifically, we observed that some DNA/RNA sequence pairs, such as Seq 1 and Seq 2, exhibit large differences in critical salt concentration yet do not form multiphase droplets. These findings imply that, beyond the heterotypic interactions between RNA-PLL and DNA-PLL, additional factors, such as homotypic interactions among DNA strands or among RNA strands, are also involved in the assembly and stabilization of multiphase condensates. This interplay highlights the importance of both the sequence composition of nucleic acids and the nature of their interactions with polycations in determining the architecture of the resulting condensates. In particular, we noticed that dA_n_/rA_n_ sequence form the multiphasic droplet, whereas T_n_/U_n_ sequence form the single-phase droplet.

To probe these interactions, we employed the lattice based “sticker-and-spacer” model, and modelled poly-L-lysine, RNA, and DNA as linear chains of 20 stickers with isotropic potentials. Homotypic PLL interactions were set to 0.1 k_B_T, reflecting electrostatic repulsion. Heterotypic PLL-RNA and PLL-DNA interactions were assigned-0.28 k_B_T and-0.2 k_B_T, respectively. When RNA-RNA interactions were set to zero, core-shell droplets formed only when DNA-DNA interactions were comparable to RNA-RNA interactions. Increasing DNA-DNA interactions to-0.1 k_B_T resulted in uniform, single-phase droplets (**Fig. 3I** and **J**), suggesting that enhanced DNA-DNA interactions relative to RNA-RNA interactions facilitate single-phase assembly formation. This conclusion is supported by comparative analyses indicating that sequences forming multiphase condensates display similar strengths of DNA-DNA and RNA-RNA interactions. In contrast, sequences that give rise to single-phase condensates exhibit stronger DNA-DNA interactions relative to RNA-RNA interactions (**Fig. 3K**). Previous studies have reported that stacking free energy for dA-dA and rA-rA, as well as for dG-dG and rG-rG, are nearly identical, whereas the stacking free energy of dT-dT exceeds that of rU-rU^58^. These differences likely underpin the preferential formation of G/A-rich oligonucleotides into multiphase structures, driven by a nuanced balance of homotypic and heterotypic interactions.

Notably, while recent studies suggest that single-chain rigidity of DNA and RNA can influence phase behavior^56, 59^, experimental measurements indicate that the persistence lengths of single-stranded DNA and RNA homopolymers-such as dT₁₉, rU₁₉, dA₁₉, and rA₁₉-are all similar, approximately 1 nm^60^. Consistently, our simulations show that single-stranded DNA and RNA possess comparable chain flexibility, as reflected by their similar radius of gyration (R_g_) distributions in the absence of PLL (**Supplementary Fig. 29**). However, conformational differences may emerge upon complexation with PLL^61^, and the specific impact of sequence-dependent rigidity on complex coacervation remains unresolved. Collectively, our data indicates that the formation of multiphase versus single-phase condensates is primarily governed by the balance between homotypic and heterotypic interactions, while the contribution of single-chain rigidity to this process warrants further investigation.

### Designer SEGREGamers enable programmable multiphase droplet formation and properties

By integrating experimental observations with atomistic and coarse-grained simulations, we demonstrate that the formation of multiphase condensates by single-stranded DNA and RNA of identical sequence is fundamentally governed by structural differences at the 2’ position of their pentose sugars. Specifically, the presence of a 2’-hydroxyl group in RNA, as opposed to a 2’-hydrogen in DNA, results in stronger RNA-PLL interactions compared to DNA-PLL interactions. Furthermore, the emergence of core-shell multiphase condensates is driven by a finely tuned balance between heterotypic (RNA-PLL *v.s.* DNA-PLL) and homotypic (RNA-RNA *v.s.* DNA-DNA) interactions in a sequence-dependent manner (**Fig. 4A**). This nonlinear and multiscale complexity precludes simple heuristic rules, motivating us to establish a predictive and interpretable machine learning framework that links sequence information directly to phase behaviour.

**Fig. 4.**
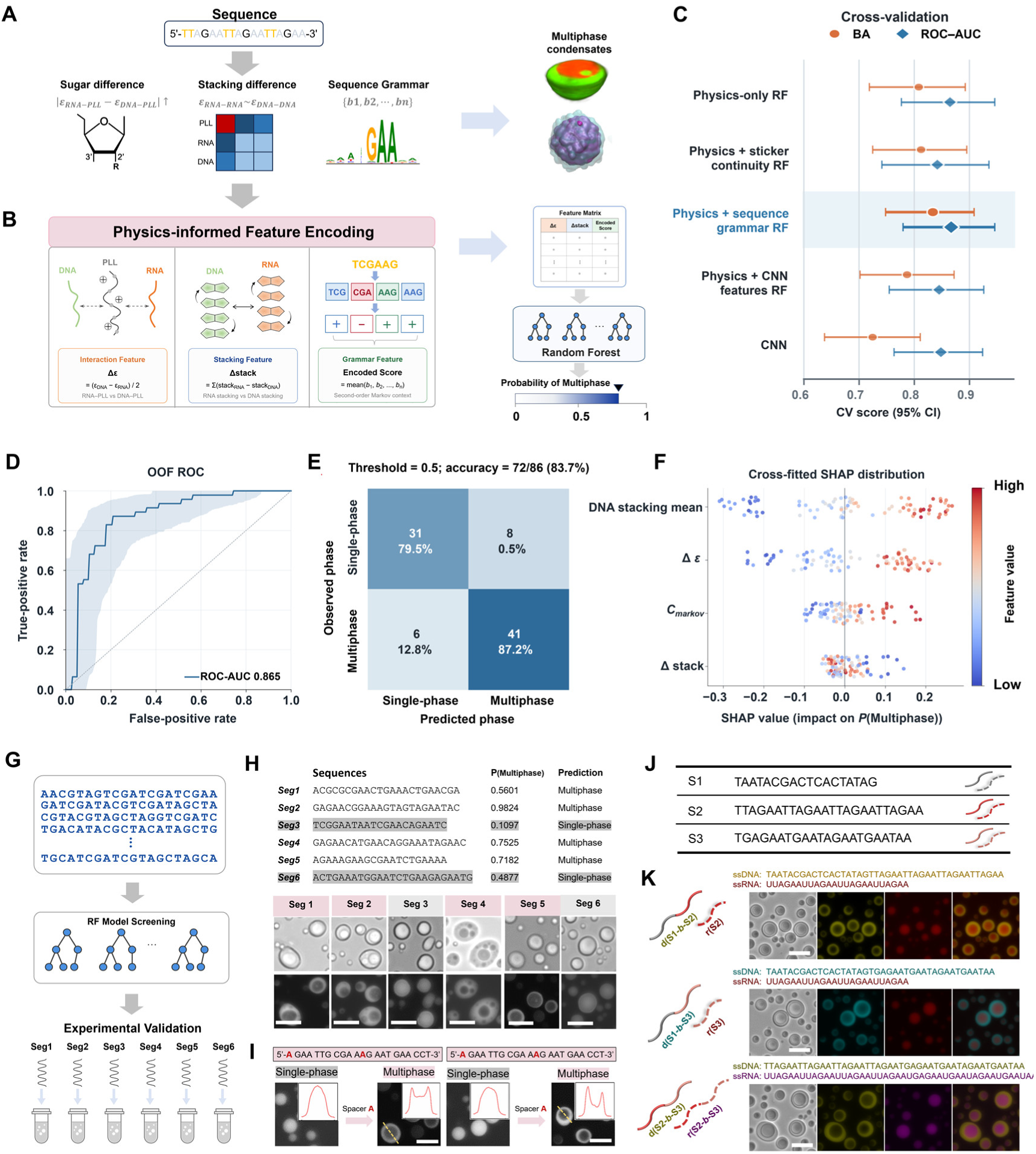
Physics-informed prediction and modular design of core-shell multiphase condensates. (**A**) Conceptual workflow from sequence encoding to prediction of multiphase condensate formation. (**B**) Physics-informed feature encoding and random-forest prediction from four sequence-derived features: Δε, Δstack, DNA stacking mean (stack_mean_) and C_markov_. (**C**) Family-grouped cross-validation of five models; the selected “Physics + sequence grammar RF” achieved a balanced accuracy of 0.834 and a ROC–AUC of 0.867 and is highlighted. Orange circles indicate balanced accuracy, blue diamonds indicate ROC–AUC, and error bars indicate 95% confidence intervals. (**D**) Out-of-fold ROC curve for the selected model (ROC–AUC = 0.867). (**E**) Confusion matrix at a classification threshold of 0.5; rows indicate the observed phase and columns indicate the predicted phase. Cell values show counts and row-wise percentages; overall accuracy was 72/86 (83.7%). (**F**) Cross-fitted SHAP distributions for the four features, ordered as DNA stacking mean, Δε, C_markov_ and Δstack; positive values favour multiphase predictions. (**G**) Frozen-model screening and experimental validation of Seg1–Seg6. (**H**) Seg1–Seg6 sequences, model probabilities, predictions and representative bright-field and fluorescence images. (**I**) Sequence redesign and corresponding phase outcomes. (**J**) Three sequence modules, S1, S2 and S3, used for modular assembly. (**K**) Multiphase condensates formed by modular diblock assemblies. Bright-field and fluorescence images are shown. Scale bars, 10 μm. Unless otherwise indicated, sequences are shown using DNA letters.

In this framework, we encoded three classes of physical determinants derived from our experimental observations into four model inputs (**Fig. 4B** and **Supplementary Text 4**): heterotypic interactions (Δε), homotypic stacking (Δstack and stack_mean_) and local sequence organization (C_markov_). The model outputs the probability of forming a core-shell multiphase condensate, P_(multiphase)_. We benchmarked five supervised learning models under a family-grouped cross-validation framework (**Supplementary Table 6**). The physics-plus-sequence-grammar random forest (RF) achieved the highest point estimates for balanced accuracy (0.834) and ROC-AUC (0.867), and was selected for its superior predictive performance and mechanistic interpretability (**Fig. 4C**). In leave-family-out validation, the selected model achieved a ROC-AUC of 0.867 (95% confidence interval, 0.754 to 0.944) and a PR-AUC of 0.847 (95% confidence interval, 0.675 to 0.962) (**Fig. 4D**). At a decision threshold of 0.5, it correctly classified 72 of 86 sequences (83.7% accuracy) (**Fig. 4E**). Cross-fitted SHAP analysis quantified the contributions of stack_mean_, Δε, C_markov_ and Δstack (in descending order), which collectively reflect three mechanistic cues: heterotypic interaction (Δε), homotypic stacking (Δstack and stack_mean_) and sequence context (C_markov_) (**Fig. 4F**).

After freezing the model and selection rules, we generated 500,000 random sequences of length 18-25 nt and screened the complete library with the frozen model (**Fig. 4G**). Candidates were filtered and paired using prespecified feature-space and novelty criteria, yielding Seg1-Seg6 for experimental validation. The predicted phase calls agreed with the experimental observations (**Fig. 4H**). Library quality control and screening records, along with complete Seg1-Seg6 sequences, experimental phases, frozen-model scores and local SHAP data, are provided in Supplementary Information (**Supplementary Text 5**).

Sequence alignment revealed that Seg5 is a variant of the single-phase sequence NRAS_M2, with two adenine residues inserted at defined positions. This insertion increased Δε from 7.77 to 9.00 and shifted Δstack from-9.58 to-12.67, with minimal change in stack_mean_ (0.983 to 0.988). The joint feature state elevated the predicted P_(multiphase)_ from 0.279 (NRAS_M2) to 0.718 (Seg5). A related redesign of hTERC_M2, yielding hTERC_M2_mut, likewise produced a clearly defined core-shell multiphase condensate (**Fig. 4I**). These results support a design principle based on the coordinated organization of heterotypic interactions, homotypic stacking and local sequence context rather than a single feature.

Notably, analysis of trinucleotide motifs among all tested sequences shows that multiphase-forming sequences exhibit a markedly higher occurrence of “GAA” motifs compared with other trinucleotide sequences (**Supplementary Figs. 31** and **32**), further corroborating the unique role of this motif. Together with the observed A-insertion effect (**Fig. 4H**), this led us to define “SEGREGamers” as DNA/RNA oligonucleotide pairs composed of multiple GAA motifs separated by adenine-rich linkers. Given the superior programmability of this design strategy, we estimated its potential diversity. Even under a conservative 10% success rate, a library of approximately 10^5^ distinct multiphase condensate types can be generated from 20-mer SEGREGamer sequences, and up to 10^9^ types from 24-mer sequences (**Supplementary Text 6** and **Supplementary Fig. 33**). This diversity is expected to increase substantially as prediction accuracy improves.

Finally, we tested the modularity and transferability of the sequence grammar by recombining three sequence modules (S1, S2 and S3) into diblock constructs (**Fig. 4J**). S1 was derived from a T7 promoter sequence, whereas S2 and S3 originated from TERRA_M2 and PU22_M2, respectively. DNA and RNA oligonucleotides of S1 alone failed to form multiphase condensates (**Supplementary Fig. 34**). By contrast, all diblock constructs combining S1 with S2 and/or S3 successfully formed multiphase condensates (**Fig. 4J** and **Supplementary Table 7**). These results confirm that the sequence information supporting core-shell formation can be decomposed, recombined and incorporated into longer nucleic-acid assemblies, demonstrating the modular nature of the identified sequence grammar. Together, these results establish a workflow from sequence feature encoding and random-forest prediction to SHAP-based interpretation, experimental validation and modular design. They demonstrate that core-shell multiphase condensates are jointly controlled by heterotypic interactions, homotypic stacking and local sequence organization, and that these behaviours can be programmed through sequence screening, site-specific insertion and modular recombination.

### Multiphase droplets as model systems to mimic nuclear condensates

For a proof-of-concept application, we elucidated the potential of multiphase droplets as synthetic nuclear compartments with spatial and functional heterogeneity. Nuclear compartments play vital roles in intracellular compartmentalization, including the uptake of small molecule therapeutics^62^, RNA biogenesis and processing, and catalytic reaction regulation (**Fig. 5A**). We first examined the partitioning of small fluorophore molecules with varying properties in multiphase droplets. As shown in **Fig. 5B** and **Supplementary Fig. 35**, all small molecules are enriched inside the multiphase condensates. Specifically, DNA-staining molecules (DAPI and SYBR Green I) partition into the DNA-rich shell. In contrast, hydrophobic molecules (ThT, rhodamine B, Nile red, and methylene blue) concentrate within the RNA-rich core phase due to the higher polymer density and reduced water content resulting from stronger RNA-PLL interactions. Additionally, negatively charged dATP preferentially localizes in the core phase where PLL concentration is highest. Collectively, these findings establish that small molecule partitioning is determined by both solute properties and phase microenvironment characteristics.

**Fig. 5.**
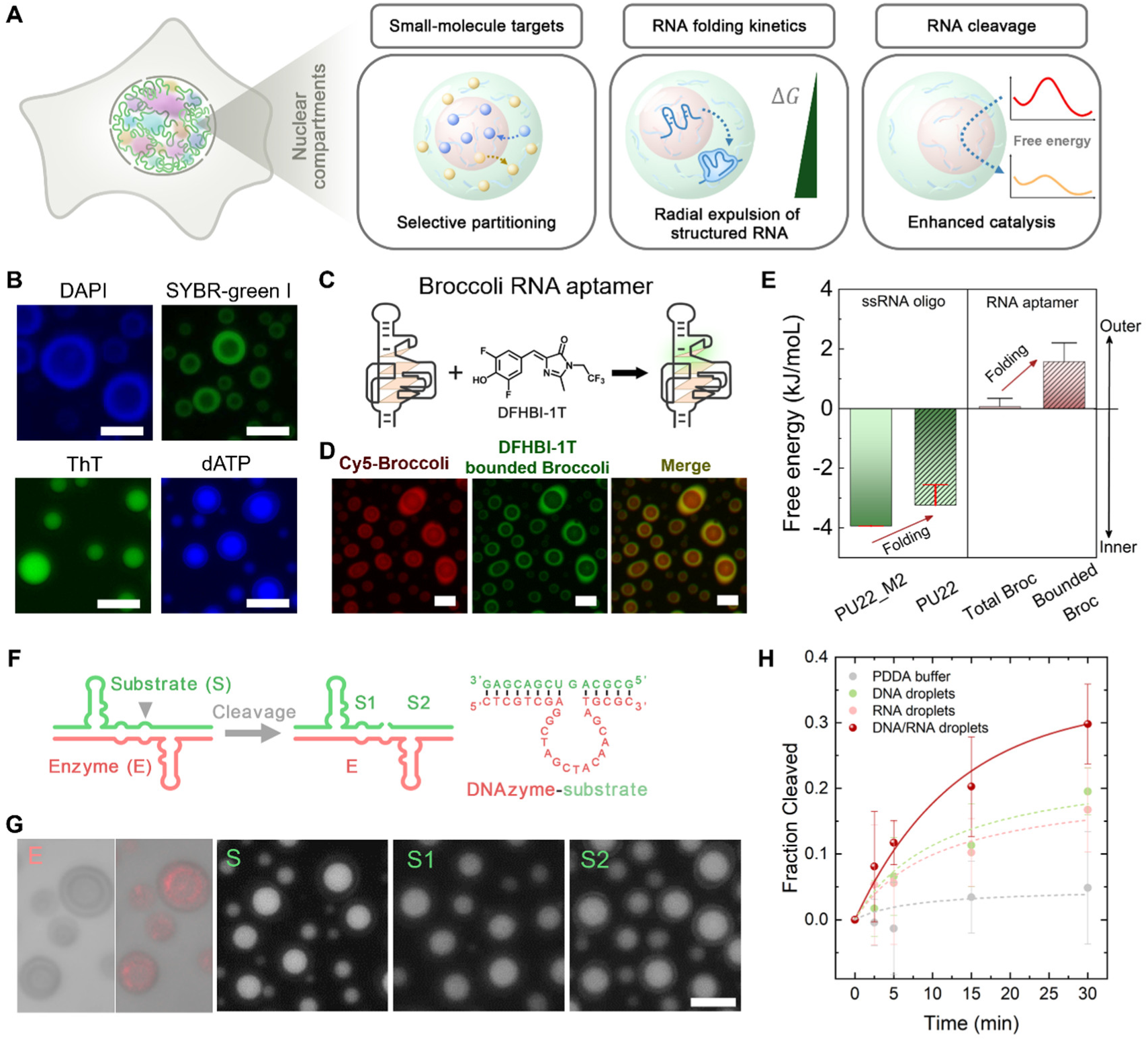
Mimicking of nuclear compartments using multiphase droplets. (**A**) Functions of nuclear compartments. (**B**) Selective partitioning of small molecules within the core and shell phase of multiphase droplets. (**C**) Broccoli RNA aptamer lights up the fluorophore molecule via folding and binding. (**D**) Distribution of the total Broccoli aptamer (red) and fluorophore-bound Broccoli aptamer (green) within multiphase droplets. (**E**) RNA structural folding altered their partitioning thermodynamics in a way from the core to shell phase. Error bars represent SEM from the analysis of thirty individual condensates. (**F**) RNA cleavage by a DNAzyme, with the cleavage site shown by the triangle. (**G**) Uptake of fluorescently labelled DNAzyme (E), RNA substrate (S), and cleaved RNA segments (S1 and S2) within multiphase DNA/RNA/PDDA droplets. Scale bar, 10 μm. (**H**) Enhanced RNA cleavage within multiphase droplets. The data points are derived from gel analysis. Error bars represent the standard deviation from four independent measurements. The curves are the best fit to the exponential equation *y* = *A* + *Be^-kt65^*. The fraction of cleaved RNA by DNAzyme at 30 minutes was significantly enhanced in DNA/RNA droplets compared to all control conditions (DNA droplets alone, RNA droplets alone, and buffer-only), as assessed by two-tailed unpaired t-tests with four biological replicates per condition (p = 0.035, p = 0.026, and p = 0.004, respectively).

We then looked at the capability of multiphase droplets as a model system for RNA biogenesis and processing using fluorophore-binding RNA aptamers. We chose the 49-nucleotide fluorescent RNA Broccoli aptamer, which activates DFHBI-1T’s green fluorescence when folded^63^ **(Fig. 5C**). To disentangle the distributions of the fluorophore-bound and unbound RNA aptamers, we used the red fluorescently labeled RNA aptamer sequence. The red fluorescence profile suggests equal distribution of total RNA aptamer molecules between shell and core regions. However, the green fluorescence demonstrated a higher intensity in the shell phase **(Fig. 5D** and **Supplementary Fig. 36A**), indicating fluorophore-bound RNA aptamers were enriched in the DNA-rich phase. The partitioning of RNA aptamer and DFHBI-1T individually within multiphase droplets suggests both entities are enriched in the core phase (**Supplementary Fig. 36B and C**). We concluded that fluorophore binding and folding of the RNA aptamer alter its partitioning behaviors within multiphase droplets, with structured and bound RNA localizing to the shell phase. This RNA structure-partitioning interplay within multiphase droplets is reminiscent of rRNA processing in the nucleolus, where fully processed and folded RNA is excluded from the core to the shell phase due to decreased valence of folded RNA^7, 64^. The preferential localization of folded Broccoli RNA aptamers within the shell region indicates that the core microenvironment disfavors the presence of structured RNA. This observation is consistent with recent findings in decapeptide-based multiphase coacervate systems, which have shown that RNA duplexes are destabilized within the core, thereby promoting the enrichment of unstructured RNA species in this region. In addition, we also tested the partitioning of rPU22, an RNA oligonucleotide known to fold into a G-quadruplex structure in the presence of KCl. Unlike its unstructured mutant control rPU22_M2, rPU22 exhibited enhanced partitioning into the shell phase (**Supplementary Fig. 37**). Compared with the partitioning difference for rPU22/rPU22_M2, the partitioning difference of folded versus unfolded Broccoli RNA aptamer (49 nt) is much more significant. These results support the view that RNA requires sufficient length or multivalency to achieve effective interaction strength differences that can significantly alter partitioning and folding dynamics in multiphase condensates, consistent with previous reports^13^. Overall, our partitioning experiments demonstrate that the distribution of guest nucleic acids within DNA/RNA/PLL multiphase condensates is governed by both their chemical identity (DNA vs. RNA) and their structural state (folded vs. unfolded). These findings suggest that such multi-compartmentalized condensates not only establish selective barriers for DNA and RNA localization but also actively influence the folding kinetics and conformational equilibria of nucleic acids between adjacent phases. Quantitatively, the Gibbs free energy of RNA partitioning (**Fig. 5E**) is consistent with the equilibrium energetic landscape of ribosome RNA exclusion within the nucleolus, highlighting the capability of multiphase droplets as synthetic nuclear compartments with functional heterogeneity.

Finally, we explored the ability of multiphase droplets to regulate RNA cleavage reactions using the 10-23 DNAzyme cleavage reaction^66^ (**Fig. 5F**). Previous studies have shown that coacervates formed by PDDA can enhance RNA catalysis by hammerhead ribozymes, owing to their lack of inhibitory interactions with nucleic acids^67^. Therefore, we selected PDDA as the polycation for assembling coacervates used in the DNAzyme-mediated RNA cleavage assays. We found that the DNAzyme (E) was enriched at the interface between the shell and the core phase, while the RNA substrate (S) and resulting cleavage products (S1 and S2, **Supplementary Table 8**) were enriched within the core phase (**Fig. 5G**). Importantly, the mobility of both the DNAzyme and RNA substrate within the multiphase condensates were preserved (**Supplementary Fig. 38**), which is critical for sustaining catalytic efficiency. Interestingly, despite the distinct partitioning behaviors of DNAzyme and RNA substrate within core-shell droplets, DNAzyme catalytic efficiency in multiphase droplets exhibited improvement compared to reactions in DNA or RNA droplets alone, particularly during the early reaction stage (**Fig. 5H** and **Supplementary Fig. 39**). The enhancement of this reaction can be attributed to two factors: 1) the interfacial accumulation of DNAzyme could generate a highly active catalytic zone, facilitating efficient contact with RNA substrates diffusing from the core and thereby enhancing catalytic activity compared to single-phase coacervates, where such spatial enrichment is absent and 2) the separation between DNAzyme and cleavage RNA products (S1 and S2), which results in a further increase in catalytic efficiency. The latter occurs as the loss of RNA substrate from the reaction is compensated for, and the newly formed RNA products are transferred to the core phase, with both processes driven by the re-equilibrated RNA partitioning. Consequently, the catalytic reaction progresses towards completion in accordance with Le Chatelier’s principle^68^. Our results demonstrate the potential of multiphase droplets as model nuclear compartments for RNA cleavage.

## Conclusions

In summary, we have demonstrated a molecular mechanism that can be deployed to form a library of genetically encoded multiphase droplets. We found the unique sugar groups in single-stranded DNA and RNA oligonucleotides, even for DNA and RNA with identical sequences, result in different interaction strengths with cationic peptides, with RNA being stronger than DNA. This difference promotes a sequence-dependent segregation between DNA and RNA oligonucleotides within complex coacervates, ultimately driving the formation of multiphase droplets featuring DNA-rich shell and RNA-rich core. Remarkably, these multiphase droplet-forming oligonucleotides, termed SEGREGamers, enable the programmed formation and biophysical properties of multiphase condensates at single-nucleotide resolution, allowing for the top-down design of condensates with desired properties for the mimicry of the heterogenous nuclear environment.

Our findings indicate that GAA-rich sequences can intrinsically modulate the balance of heterotypic and homotypic interactions among DNA, RNA and peptides. Recent discoveries showed that *in vivo*, sequence-specific homotypic RNA-RNA interactions among transcriptome mRNA, scaffolded by protein scaffolds, can drive the formation of membraneless organelles with multiphasic organizations^69^. Our study extends this conceptual framework by demonstrating that not only RNA-RNA but also DNA-DNA homotypic interactions, when scaffolded by polycations, can drive the formation of multiphase condensates. This mechanism parallels emerging evidence for protein-based demixing phenomena, such as those observed in postsynaptic density condensates^70^. Collectively, these findings suggest that the principles of sequence-encoded multiphasic organization are broadly applicable across different classes of biomolecules.

Multiphase coacervate droplets, with their structural and functional heterogeneity, have been suggested as synthetic compartments resembling intracellular membraneless organelles^71^, potentially playing crucial roles in the origin of life^72^. Our research presents a robust synthetic pathway for multiphase droplets using basic DNA and RNA building blocks, without relying on complex competing biomolecular interaction networks. Thus, our multiphase droplet formulation represents a minimal model system of transcriptional condensates around the genome that share components but remain mutually immiscible^73^. These droplets can be integrated into existing protocell systems as synthetic nuclear compartments for cellular heterogeneity^74^, and into the ‘RNA-DNA world’ hypothesis for early molecule catalysis and evolution^75, 76^. Furthermore, with potentials in achieving designer functions from designer sequences, the DNA/RNA-encoded immiscibility can be combined with DNA/RNA nanotechnology to enable precise RNA delivery^77^, multiscale superstructure assembly^78^, and DNA based hydrogels^79^.

## Methods

### Materials

DNA and RNA oligonucleotide sequences were purchased from IDT and used without further purification, as listed in **Supplementary Table 9**. Poly-L-lysine (PLL, 30-70 kDa, hydrobromide, P2636), Poly-D-lysine (30-70 kDa, hydrobromide, P7280), Poly-L-Arginine (70-150 kDa, hydrobromide, P1274), Polydiallyldimethylammonium chloride (PDDA, average M_w_<100000, 522376), Polyethyleneimine (PEI, average M_w_ ∼ 25000, 408727) and fluorescein isothiocyanate-labeled poly-L-lysine (FITC-PLL, 30-70 kDa, P3069) were purchased from Sigma-Aldrich, Inc. The averaged molecular weight of PLL is assumed to be 50 kDa, corresponding to an average of 240 lysine residues per chain. SUP-K72 was expressed and purified from Escherichia coli BLR (DE3) cells. The FUS (Fused in Sarcoma) stock solution contained 30 μM FUS protein in a buffer composed of 50 mM Tris(hydroxymethyl)aminomethane (Tris), 1 M potassium chloride (KCl), 1 mM dithiothreitol (DTT), and 5% glycerol. Detailed protocols for the preparation of full-length wild-type FUS and SUP-K72 are described in previous publications^20, 80^.

### Sample preparation

Stock solutions of DNA and RNA of 20 mM, and PLL of 100 mM total residue (or total charge) were prepared and stored under-20 ℃. In the aqueous mixture, each lysine residue contributes one positive charge, and each DNA/RNA phosphate group contributes one negative charge. Hence, the negative and positive charge concentration in the mixture is defined as [e-]= [DNA phosphate]+[RNA phosphate] = n×([DNA] + [RNA]) and [e+] = [PLL amine] =240×[PLL] respectively, where n is the number of nucleotides on a single DNA/RNA sequence, and [DNA], [RNA] and [PLL] denote the concentration of DNA, RNA and PLL in the mixture, respectively. Unless specified, [DNA phosphate] = [RNA phosphate]. In a typical experiment, an DNA/RNA solution with the target negative charge concentration containing 500 mM KCl and 40 mM Tris buffer (pH=7.5) was renatured by heating to 95 ℃ for 3 minutes using a PCR thermal cycler and then cooled down to room temperature, followed by incubation for 30 minutes. A PLL solution with the target positive charge concentration with KCl (500 mM) and 40 mM Tris buffer (pH=7.5) was separately prepared and then mixed thoroughly with the DNA/RNA solution using pipette. Salt concentration of KCl and NaCl in each experiment were adjusted according to the need. Similar protocols were employed for testing various polycations, oligopeptides, and SUP-K72. For FUS assays, 1 μL FUS stock solution was homogeneously mixed with 19 μL bulk solutions, including DI water, DNA, and RNA. The final buffer contains 2.5 mM Tris, 0.05 M KCl, 0.05 mM DTT, and 0.25% glycerol. The specific experimental conditions under which the condensates shown in each figure were assembled are provided in **Supplementary Table 10**.

### Fluorescence imaging method

We performed imaging around 10 minutes after mixing the DNA/RNA solution and the PLL solution. Samples were introduced into customed chambers made with spacers. To inhibit the coacervate droplets settling down and then wetting the wall of the glass chamber, glass slides were first cleaned by ethanol and water to remove the contaminants, then immersed in the Pluronics F-127 solution (1 wt%) for around 1 hour and carefully rinsed using MilliQ water afterwards, followed by heated and dried inside the 65 ℃ oven for 1 hour. Bright field images of the coacervates were obtained on an inverted microscope equipped with a 40X objective (Olympus, CKX53). Green fluorescence of FAM-labeled DNA and red fluorescence of Cy5-labelled RNA were activated with laser (Olympus, U-RFL-T) of the 488 nm and the 607 nm wavelength, respectively. All confocal images were acquired using a Nikon AX R microscope (AX R, Nikon Instruments Inc., Japan). For the analysis of condensates, z-stack images were captured with a 60× oil immersion objective. A resonant scanner was used for image acquisition, and each frame was averaged four times to enhance the signal-to-noise ratio. All images were representative of at least three independent tests and analyzed using the ImageJ software.

### Fluorescence recovery after photobleaching (FRAP)

FRAP experiments were performed on a Carl Zeiss LSM 880 confocal microscope with a ×40 oil immersion objective. An area with radius of 0.5 μm was bleached with a 488-nm or 561-nm laser at 100% power; subsequent recovery of the bleached area was recorded with a 488-nm or 561-nm laser. Correction for photobleaching and normalization was performed via a custom code in Matlab. The final FRAP recovery data is the average of recovery curves collected from five independent measurements.

### Flow cytometry

RNA-PLL droplets and DNA-PLL droplets were prepared with Cy5-RNA (2 μM) and FAM-DNA (2 μM), respectively. DNA-RNA-PLL multiphase droplets were stained with both Cy5-RNA and FAM-DNA. All samples were prepared at the charge concentration of [e+] = [e-] = 15 mM. Fluorescence intensity of 200 µL droplet sample were tested by the flow cytometer (BD FACSAriaIII) under the wavelength of 488/530 and 633/660. and the data wereanalyzed by FlowJo software (BD, USA).

### Evaporation assay

A 0.5 μL sample containing DNA/RNA/PLL coacervates was pipetted onto a clean microscope glass slide under room conditions (Temperature = 25 °C, Humidity = 55-65%), forming an evaporating sessile droplet. ISOLAB GmbH microscope glass slides served as solid substrates for the droplets. To remove surface contaminants, the slides were first wiped with ethanol-moistened tissue, then sonicated in ethanol for 15 minutes, and finally rinsed with isopropyl alcohol and Milli-Q water. The cleaned slides were dried with nitrogen flow and placed in an oven at 65 °C for 1 hour. Fluorescence images of the evaporating droplets were captured using a Nikon Ti2-E Widefield microscope and processed with ImageJ (NIH) software.

### CP-AFM measurements

Mechanical properties of DNA-and RNA-PLL droplets were characterized by an atomic force microscope (NanoWizard II, JPK). A spherical silica particle (*R* ≍ 4 *μ*m) was affixed to an AFM cantilever with a 0.1 N/m spring constant, following protocols described by Li and co-workers^46^. The spring constant was calibrated through a contact-based technique. AFM measurements were conducted by submerging the probe in a glass Petri dish containing 2 mL coacervate droplet suspensions. Prior to measurement, the probe and Petri dish underwent plasma treatment to eliminate contaminants and were rinsed with water to neutralize the surface. Droplet suspensions were prepared as previously mentioned, but with an extremely low concentration (0.6 mM total charge concentration, [e+] = [e-] = 0.3 mM) to improve laser signal gain. Freshly prepared droplet suspensions were always used, allowing for approximately 20 minutes of incubation to reach equilibrium. During a typical measurement, the probe made contact with the Petri dish’s bottom at a constant velocity (1 µm/s) to establish zero-separation distance and initiate capillary condensation. The probe was held at zero-separation for 10 seconds to facilitate capillary bridge formation. Subsequently, the probe was raised at varying velocities between 0.2 µm/s and 2 µm/s, covering a 5 µm distance for each sample. For every retraction rate, twenty measurements were performed, and force-distance curves were monitored for both approach and retraction steps. Data analysis involved using JPK’s data processing software to correct the vertical tip position resulting from cantilever bending. The cantilever deflection *Δx* was transformed into force using Hooke’s law, *F* = *kΔx*, where k represents the cantilever’s spring constant.

### Electrophoretic mobility shift assay (EMSA)

Binding reactions were setup of a volume 10 µL containing either 200 nM DNA TERRA_M2 or RNA TERRA_M2 and a serial dilution of PLL in 10 mM Tris pH 7.5, 100 mM KCl, 5% glycerol. The PLL concentration was varied by serial dilution to be 0, 7.8125, 15.625, 31.25, 62.5, 125, 250, 500 or 1000 nM. Binding reactions were assembled, vortexed, spun down and incubated for 30 minutes at room temperature. Binding reactions were then loaded onto a 0.5X TBE native 10% polyacrylamide gels and run at 100 V for 30 minutes. The gels were then stained using SYBR gold nucleic acid gel stain (Invitrogen) and visualized using a ChemiDoc MP (Bio-Rad). Band intensity was measured using ImageJ before data plotting and curve fitting using Origin 8.5. The Hill equation was used for curve fitting to deduce the K_D_.

### Circular dichroism (CD)

DNA and/or RNA oligos (5 µM) in Tris-HCl buffer containing 25 mM Tris-HCl and 150 mM KCl/LiCl (pH 7.5) were heated at 95 °C for 3 min and cooled down at room temperature for 10 min. CD spectra were recorded on a Jasco CD J-150 spectrometer with 1-cm path length quartz cuvette. Three scans from 220 nm to 340 nm at 1 nm intervals were accumulated and averaged.

### Droplet Microfluidics

Microfluidic devices was prepared using a typical soft lithography replica molding technology. First, a SU-8 photoresist (2025, MicroChem, USA) mold with desired channel geometry was fabricated onto a silicon wafer using maskless lithography (SF-100 Xcel, Intelligent Micro Patterning, LLC, USA). Then, the pre-polymer base and curing agent of PDMS (Sylgard 184, Dow Corning, USA) was mixed at a weight ratio of 10:1, degassed, and then poured onto the photoresist mold and solidified at 65 ℃ overnight. Subsequently, the PDMS layer was detached from the mold, and bonded to a glass substrate by oxygen plasma treatment. To fabricate the electrodes, a low-melting-point metal wire (52225, Indium Incorporation, USA) was inserted into the inlets of the designed channel near the expansion chamber and then introduced into the whole channel under negative pressure applied to the outlets at 100 ℃. DNA/RNA and PLL droplets were generated using fluorinated oil (HFE7500, 3M, USA) supplemented with 2% (w/w) of surfactant (RAN Biotechnologies, USA) as the continuous phase. The two batches of droplets were reinjected into the droplet merging platform and separated by a gapping oil. An alternating current (AC) signal with an amplitude of 750 Vp-p and a frequency of 30 kHz was applied to induce droplet coalescence.

### Encapsulation of small molecules

For partitioning experiments, 10 μL aliquots of a DNA/RNA/PLL multiphase droplet of TERRA_M2 in the buffer containing 500 mM KCl and 40 mM Tris (pH 7.5) were added to sample chambers on a cover glass slide, with the charge concentration of [e-] = [e+] = 5 mM. Small quantities of the stock solutions of the dye molecules were added to the multiphase droplets, mixed by gentle pipetting, and visualized by excitation at the indicated wavelengths. DAPI (100 μM) and dATP (50 μM) were excited at 380 nm. ThT (50 μM) and SYBI Green I (0.01 X concentrate) were excited at 488 nm. NMM (10 μM), Nile Red (50 mg/mL), Rhodamine B (100 μM) and Methylene Blue (100 μM) were excited at 561 nm.

### Encapsulation of guest RNA

A mixture of 2 µM FAM-labeled rPU22 or rPU22_M2 guest molecules and DNA/RNA oligos of TERRA_M2 was prepared in a buffer containing 500 mM KCl and 40 mM Tris (pH 7.5), with a charge concentration of [e-] = [e+] = 5 mM. The mixture was renatured at 95 ℃ for 3 minutes, cooled to room temperature, and incubated for 30 minutes. PLL was then added to the mixture and mixed thoroughly using a pipette. The mixed samples were incubated at room temperature for 10 minutes before imaging with a Leica inverted microscope. The partition coefficient (K) was calculated from average fluorescence intensities using the formula K = (I_c,in_-I_b_) / (I_c,out_ - I_b_), where I_b_, I_c,out_, and I_c,in_ represent the intensity of the dilute phase surrounding the multiphase droplets, the outer shell layer, and the inner core phase of the multiphase droplets, respectively. The Gibbs free energy change for partitioning was determined based on Δ*G* = −*RT* ln *K*, where R is molar gas constant and T is temperature. The standard deviation of Δ*G* was obtained by propagating the experimental errors from the measurements of K. The standard deviation of Δ*G* was obtained by propagating the experimental errors from the measurements of K.

### Encapsulation of Broccoli RNA aptamer

A mixture containing 5 µM Broccoli RNA aptamer and/or 200 µM DFHBI-1T was first mixed with DNA/RNA oligos in a buffer containing 150 mM KCl, 40 mM Tris (pH 7.5), and 5 mM MgCl_2_. To create multiphase droplets at 150 mM KCl, rPU22_M2 and d(S1-b-S2) were employed, maintaining a charge concentration of [e-] = [e+] = 2.5 mM. The mixture was renatured at 95 ℃ for 3 minutes, cooled to room temperature, and incubated for 30 minutes. PLL was then added to the mixture and mixed thoroughly using a pipette. The mixed samples were incubated at room temperature for 10 minutes before imaging with a Leica inverted microscope. The partition coefficient (K) was calculated from average fluorescence intensities using the formula K = (I_c,in_ - I_b_) / (I_c,out_ - I_b_), where I_b_, I_c,out_, and I_c,in_ represent the intensity of the dilute phase surrounding the multiphase droplets, the outer shell layer, and the inner core phase of the multiphase droplets, respectively. The Gibbs free energy change for partitioning was determined based on Δ*G* = −*RT* ln *K*. The standard deviation of Δ*G* was obtained by propagating the experimental errors from the measurements of K.

### Uptake of DNAzyme and substrate RNA

Guest molecules, including 5 µM Cy5-labeled DNAzyme, 2 µM FAM-labeled RNA substrate, RNA substrate 1, and RNA substrate 2, were individually incorporated into a buffer containing 40 mM Tris (pH 7.5), 10 mM MgCl2, and 150 mM KCl. The final sample volume was 10 μL. To create multiphase droplets at 150 mM KCl, rPU22_M2 and d(S1-b-S2) were utilized, maintaining a charge concentration of [e-] = [e+] = 2.5 mM. Samples were prepared by adding a PDDA solution to the renatured nucleic acid buffer.

### DNAzyme cleavage reaction

10-23 DNAzyme (1 μM) in 25 mM Tris-HCl buffer containing 5 mM KCl and 5 mM MgCl_2_ (pH=7.5) was heated at 95 °C for 3 min and cooled down at room temperature for 10 min. Then DNAzyme were mixed with FAM-labeled RNA substrate (50 nM) that was partitioned into coacervates composed of PDDA (1 mM [e+]), rPU22_M2 and/or d(S1-b-S2) (1 mM [e-] in total). After incubating at 37 ℃ for 1 h, 1 M of NaCl was added to quench reactions and disrupt coacervates. Then the results were analyzed using 15% denaturing PAGE (polyacrylamide gel) and scanned by FujiFilm FLA-9000 Gel Imager at 500 V. The data is fit to the exponential equation *y* = *A* + *Be*^ି^*^kt^*.

### All-atom molecular dynamics simulations

All simulations were performed using the GROMACS 2021 package^81–83^ on the GPU nodes of HPC2021 at The University of Hong Kong. The Amber ff14SB^84^, OL15^85^, OL3^86^ force field was used for the protein, DNA and RNA, respectively. The TIP3P water model^87^ was used. The temperature was kept at 300 K using the V-rescale thermostat^88^ with a coupling time constant of 0.1 ps. The Parrinello-Rahman method^89^ was used to maintain the pressure at 1 bar with a coupling time constant of 2.0 ps. Periodic boundary condition was sued and the Particle Mesh Ewald (PME) algorithm^90, 91^ was applied to calculate the long-rang electrostatic interactions. The cutoffs of electrostatic potential and van der Waals potential were set to 1.2 and 1.1 nm. The LINCS algorithm ^92^ was used to constrain all bonds with H-atoms. The Umbrella sampling^93^ was used to calculate the potential of mean force (PMF) between the capped Lys residue (Ace-Lys-Nme) and nucleic acids (dA_2_, rA_2_, rC_2_, dC_2_, rG_2_, dG_2_, rU_2_ or dT_2_). The initial structures for the peptide and nucleic acids were obtained from the CAMPARI molecular simulation software (https://campari.sourceforge.net/). A pair of one capped Lys residue and one nucleic acid were solvated in the cubic box with the length of ∼ 5 nm. A harmonic potential with 0.5 kcal/mol/Å^2^ was used to restrain the distance between the capped Lys residue and the nucleic acid of interest in each window. The nitrogen atom in the sidechain of Lys and phosphate atom in the nucleic acid backbone were used to apply the harmonic potential. The centers of the umbrella sampling windows were 3 Å, and 4 Å to 20 Å with an interval of 2 Å. In each window, 24 independent simulations from different starting structures were run, and each independent simulation was 40 ns. The WHAM method^94, 95^ was used to reconstruct the PMF curves from simulations.

Simulated tempering^96^ was employed to enhance sampling of the radius of gyration (Rg) for the single-stranded nucleic acids: rA_12_, dA_12_, rU_12_, dT_12_, r[GAA]_4_, d[GAA]_4_, r[UUAGAA]_2_, and d[TTAGAA]_2_. Initial structures for these nucleic acids were obtained from the CAMPARI molecular simulation software (https://campari.sourceforge.net/). Each single-chain nucleic acid was solvated to a cubic box with the length of ∼ 9 nm. The temperature for simulated tempering spans from 300 K to 600 K with an interval of 7.5 K. In order to choose the initial weights, 10 ns-long simulations at each temperature from 300 K to 600 K with an interval of 2 K were performed. Then the weights for these temperatures were calculated using the approach described by Park and Pande^97^ and the initial weights for the temperature of interest was obtained by interpolating the calculated weights. The Metropolis transition was used to sample the temperature space and Wang-Landau algorithm^98^ was used to update the weights. For each system, 4 independent simulations were performed, each independent simulation is 1000 ns long.

### Coarse-grained simulations

The coarse grained simulations were performed using LaSSI simulation engine^99^. The PLL, RNA and DNA molecules were represented by sticker chains with the length of 20. The cubic simulation box with length of 200 contains 1000 PLL chains, 500 RNA chains, and 500 DNA chains. All molecules were randomly placed in the simulation box without overlap. Subsequently, a biasing potential was applied to all the beads to drive them toward the center of the simulation box. Next, an equilibrium process was performed for 1 × 10^8^ Monte Carlo steps, during which the effective temperature was gradually reduced from 1000 to 1. Following equilibration, a production run of 2 × 10^10^ Monte Carlo steps were performed. All stickers interacted via isotropic interactions. The homotypic interactions for PLL beads were set to 0.1 kT to capture the repulsive electrostatic interactions, while the heterotypic interactions for PLL-RNA and PLL-DNA were set to-0.28 k_B_T and - 0.20 k_B_T, respectively. Homotypic interactions between RNA beads were 0. When the homotypic interactions between DNA beads were identical to those of RNA molecules, core-shell structures were observed. However, when the homotypic interactions between DNA were-0.1 k_B_T, one-phase droplets formed. The relative moving frequencies of Monte Carlo moves are listed in the following table.

| Move type | Relative frequency |
| --- | --- |
| Local | 1000 |
| Co-local | 200 |
| Multi-local | 200 |
| Snake | 400 |
| Chain translation | 200 |
| Small cluster translation | 5 |
| Cluster translation | 1 |
| Chain pivot | 200 |
| Double pivot | 100 |

## Data availability

The data that support this study are available within the paper and its Supplementary Information.

## Code availability

All-atom and coarse-grained simulation codes used in this study can be obtained from the corresponding authors upon request.

## Acknowledgements

We thank Prof. David A. Weitz and Prof. Julian A. Tanner for their continued interest in this work and for their insightful feedback. We gratefully acknowledge Prof. Andreas Herrmann and his research team for generously providing the SUP-K72 samples. This work as supported by the General Research Fund (Nos. 17306221, 17317322, and 17306820) from the Research Grant Council (RGC) of Hong Kong, as well as the National Natural Science Foundation of China (NSFC)-RGC Joint Research Scheme (N_HKU718/19). H.C.S. was funded in part by the Croucher Senior Research Fellowship from Croucher Foundation and the Health@InnoHK program of the Innovation and Technology Commission of the Hong Kong SAR Government

## Author contributions

W.G., Y.S., T.K. and H.C.S conceived and designed the project. W.G., X.L., A.B.K, H.O., Y.P., R.L., Y.W., K.K.L., T.M., F.W. and Z.Y. performed the experiments and analysed the data. F.C. and X.Z. performed molecular dynamic simulations. H.C.S supervised the study. W.G., Y.S., T.K., X.Z. and H.C.S wrote the manuscript. All authors discussed the results and commented on the manuscript.

## Competing interests

X.L. is a cofounder and director of Upgrade Biopolymers Limited, the work in this paper is not directly related to the work of the company. H.C. Shum is a scientific advisor of EN Technology Limited, MicroDiagnostics Limited, and PharmaEase Technology Limited in which he owns some equity, and a managing director of the research centre, namely Advanced Biomedical Instrumentation Centre Limited. The works in the paper is, however, not directly related to the works of these entities, as far as we know.

## Notes

### Summary of Updates

A supervised machine learning framework was employed to analyze the sequence dependency of multiphase condensate formation.

