## Supplementary Information for "Pentose Sugars Encode Sequence-Dependent DNA-RNA Segregation for Programmable Engineering of Multiphase Condensates"

### Contents:

**Supplementary Text**

Supplementary Text 1 to 6

**Supplementary Figures**

Supplementary Fig. 1 to 39

**Supplementary Tables**

Supplementary Table. 1 to 10

**Supplementary Videos**

Supplementary Videos 1 to 3

**References**

### Supplementary Text

#### 1 Interfacial tension measurement

We investigate the mechanical properties of DNA- and RNA-PLL droplets by a technique based on colloidal probe atomic force microscopy (CP-AFM), as shown in Fig. 2A. The CP-AFM method has been reported earlier by Sprakel *et al.*[1], Spruijt *et al.*[2], Li *et al.*[3], and Lim *et al.*[4] in the use of determining the low interfacial tension between biomolecular condensates and their co-existing phase. In brief, this technique measures the force-distance curve from retracting a capillary bridge between the probe particle and substrate formed by either capillary condensation in condensate dispersions or contacting a substrate with a condensate layer. The retraction causes an increase of the surface area of the bridge and a force response, from which the interfacial tension and rheological properties of condensates can be inferred.

A low polymer concentration was employed in the CP-AFM measurements to promote capillary condensation and bridge formation between the colloid probe and the substrate. Crucially, even at this low concentration, DNA-RNA segregation within the coacervates was still observed (Supplementary Fig. 10A), indicating that phase separation is robust. Furthermore, the measured interfacial tension is expected to be independent of the total polymer concentration. This is because, at equilibrium and along a single tie line in the phase diagram, the polymer concentrations in the coacervate and dilute phases are constant[5].

As introduced by Sprakel *et al.*[1], we use capillary condensation to form a bridge between two surfaces (probe and substrate). If the condensate wets both surfaces, capillary condensation could occur spontaneously when immersing the probe in condensate suspensions, and when both surfaces are close enough. We measured the force response of samples containing only buffer solutions or single polymer components of the condensates. None of these samples show obvious attraction during approaching or retraction steps. Therefore, the strong attraction from condensate samples is mainly due to the formation of the capillary bridge.

In Fig. 2B, we have observed an attractive force for DNA- and RNA-PLL droplets. We think this initial attraction is caused by the nucleation of the condensed phase. The rate of nucleation is proportional to $\exp(-\gamma A/k_B T)$, where $\gamma$ is the interfacial tension and $A$ represents the surface area of the critical nucleus[1]. The rate of nucleation should increase with decreasing interfacial tension. Therefore, for coacervates which have low interfacial tensions (usually less than 1 mN/m) with their co-existing phase, the nucleation should take place very fast. Furthermore, RNA-PLL droplets show more significant attraction compared with DNA-PLL droplets, indicating higher cohesive strength (Fig. 2B). Such observation is in good agreement with the information obtained from critical salt experiments (Fig. 2A), where RNA-PLL droplets show higher critical salt concentration, indicating the stronger internal interaction strength than that of DNA-PLL condensates.

We can determine the interfacial tension of a droplet that forms a capillary bridge following the method of Li *et al*[3]. For simplicity, we treat the peak force $F$ (Supplementary Fig. 10B) occurs at zero separation for all measurements. Therefore, a well-known estimation for the sphere-plane geometry can be used to calculate the interfacial tension from the force-distance curve:

$$F = 4\pi R\gamma\cos\theta$$

where $R$ is the radius of the probe, $\gamma$ is the interfacial tension, and $\theta$ is the contact angle between the capillary bridge and substrate, which was simplified as complete wetting ($\cos\theta \approx 1$) in our calculation. Based on the above equation, the interfacial tension of DNA-PLL and RNA-PLL droplets with their co-existing phase is 892.0 (±54.9) $\mu$N/m. and 1976.8 (±85.8) $\mu$N/m, respectively (Fig. 2C).

**2 Derivation of the $\chi$~$\gamma$ relationship**

The detailed derivation of such a relationship based on the lattice model is given in the reference 54 in the main text (Dill, K. & Bromberg, S. Molecular Driving Forces: Statistical Thermodynamics in Biology, Chemistry, Physics, and Nanoscience).

The free energy of the lattice model liquid is

$$F = U - TS$$

The translational entropy of the lattice of particles is zero, because if pairs of particles trade positions, the rearrangement can't be distinguished from the original arrangement. The lattice liquid has $S = 0$ and $F = U$.

The boundary between two condensed phases is an interface. The interfacial tension $\gamma_{AB}$ is the free energy cost of increasing the interfacial area between phases A and B. If $\gamma_{AB}$ is large, the two media will tend to minimize their interfacial contact. To determine $\gamma_{AB}$ by using the lattice model, molecules of types A and B are assumed to be identical in size.

Suppose there are $N_A$ molecules of A, $n$ of which are at the interface, and there are $N_B$ molecules of B, $n$ of which are at the interface in contact with A. (Because the particles have the same size in this model, there will be the same number $n$ of each type for a given area of interfacial contact). The total energy of the system is treated as it was for surface tension with the addition of $n$ AB contacts at the interface

$$U = (N_A - n)\left(\frac{zw_{AA}}{2}\right) + n\left(\frac{(z-1)w_{AA}}{2}\right) + nw_{AB} + (N_B - n)\left(\frac{zw_{BB}}{2}\right) + n\left(\frac{(z-1)w_{BB}}{2}\right)$$

Where the attraction between two particles of type A is represented by a "bond" energy $w_{AA}$, and so is for $w_{AB}$ and $w_{BB}$. Each particle on a lattice has $z$ nearest neighbors (z is called the coordination number of the lattice). Because the entropy of each bulk phase is zero according to the lattice model, the interfacial tension is defined by

$$\gamma_{AB} = \left(\frac{\partial F}{\partial A}\right)_{N_A,N_B,T} = \left(\frac{\partial U}{\partial A}\right)_{N_A,N_B,T} = \left(\frac{\partial U}{\partial n}\right)_{N_A,N_B,T}\left(\frac{dn}{dA}\right)$$

where A is the total area of the surface in lattice units. And

$$\frac{dn}{dA} = \frac{1}{a}$$

where $a$ is the area per molecule exposed at the surface. Based on the first equation,

$$\left(\frac{\partial U}{\partial n}\right)_{N_A,N_B,T} = w_{AB} - \frac{w_{AA} + w_{BB}}{2}$$

Combine the definition of $\chi_{AB}$

$$\chi_{AB} = \frac{z}{kT}\left(w_{AB} - \frac{w_{AA} + w_{BB}}{2}\right)$$

an expression for $\gamma_{AB}$ is assembled:

$$\gamma_{AB} = \frac{1}{a}\left(w_{AB} - \frac{w_{AA} + w_{BB}}{2}\right) = \left(\frac{kT}{za}\right)\chi_{AB}$$

Conclusion: If the entropy contribution can be neglected, we have $\gamma_{AB} \sim \chi_{AB}$

### 3 Fitting of the phase diagram

To fit the phase diagram of DNA-PLL and RNA-PLL condensates, we used an adapted Voorn-Overbeek (VO) model, which we have successfully applied in our previous work[6]. In this model, the dimensionless free energy per lattice site is given by:

$$f = \frac{\phi_n}{N_n}\ln(\phi_n) + \frac{\phi_p}{N_p}\ln(\phi_p) + \Omega\ln(\Omega) + (1-\phi_n-\phi_p-\Omega)\ln(1-\phi_n-\phi_p-\Omega) + g(\phi_n,\phi_p,\Omega)$$

where $\phi_n$ and $\phi_p$ represent the volume fractions of single-stranded nucleotides (DNA or RNA) and PLL, respectively, while $\Omega$ is the volume fraction of salt. $N_\phi$ and $N_\psi$ are the degree of polymerization for nucleotides and PLL, taken as 24 and 240, respectively.

The term $g(\phi_n,\phi_p,\Omega)$ accounts for the enthalpic contributions from electrostatic interactions between the oppositely charged species, as well as salt screening, and is expressed as:

$$g(\phi,\psi,\Omega) = -\alpha(\phi_n+\phi_p+\Omega)^{3/2}$$

Here,$\alpha$ characterizes the strength of electrostatic interactions, and crucially, is different for DNA-PLL and RNA-PLL systems. In our modeling, we used $\alpha = 1.0$ for DNA-PLL and $\alpha = 1.1$ for RNA-PLL, reflecting the stronger interaction of RNA with PLL due to its ribose chemistry.

The binodal curves in the phase diagram were derived by numerically solving for the equilibrium conditions of osmotic pressure and chemical potential between the coexisting dilute and condensed phases. This approach enables us to capture the experimentally observed trend that RNA–PLL interactions are stronger than DNA-PLL, as shown in Supplementary Fig. 14. This interaction difference plays the dominant role in the partial immiscibility and the formation of multiphase condensates with spatial segregation of DNA-rich and RNA-rich domains.

### 4. Physics-informed random-forest model

After excluding precipitate-forming sequences, we formulated the remaining sequences as a binary classification problem that distinguished core-shell multiphase condensates from single-phase droplets. The training set comprised 86 sequences, including 47 core-shell and 39 single-phase sequences.

Three mechanistic dimensions were encoded by four sequence-derived features. Heterotypic interaction asymmetry was represented by

$$\Delta\varepsilon = \frac{E_{DNA-PLL} - E_{RNA-PLL}}{2}$$

The interaction energies were calculated by summing base-specific interaction parameters along each sequence. We defined Δstack as the difference between RNA–RNA and DNA–DNA stacking energies, representing their relative homotypic stacking preference. We calculated $stack_{mean}$ as the mean absolute DNA stacking strength across all overlapping dinucleotides. This feature represents the overall stacking background. Local sequence organization was represented by the second-order Markov context feature $C_{markov}$.

For each dinucleotide context ab and following base c, we counted overlapping trinucleotides "$N_aN_bN_c$" separately in the single-phase and core-shell classes. We estimated class-conditional transition probabilities with Laplace smoothing:

$$P_y(c|ab) = \frac{N_y(ab,c) + 1}{\Sigma_{c'} N_y(ab,c') + 4}$$

For a sequence $x_1x_2\ldots x_L$, we defined $C_{markov}$ as the mean class-conditional log-likelihood ratio over all overlapping second-order transitions:

$$C_{markov} = \frac{1}{L-2}\sum_{k=3}^{L} log(\frac{P_{core-shell}(x_k|x_{(k-2)}x_{(k-1)})}{P_{single-phase}(x_k|x_{(k-2)}x_{(k-1)})})$$

$C_{markov}$ therefore captured base order and local sequence context rather than simple motif or k-mer counts. To prevent label leakage, we fitted each Markov transition table only to the sequences and labels in the corresponding training fold. We calculated $C_{markov}$ for validation sequences using the transition table derived from the training fold.

We used the Physics + sequence grammar RF (physics_markov_rf) model. The classifier contained 300 trees with a maximum depth of 5, Gini impurity, square-root feature subsampling, bootstrap sampling, and balanced class weights. We set the random state to 42. The model returned $P_{(multiphase)}$, and binary calls used a fixed threshold of 0.50. After model selection, we froze the model parameters, feature definitions, Markov estimation procedure, and classification threshold. We did not adjust them using subsequent candidate sequences or experimental results. The five-model comparison is summarized in Supplementary Table 6. GAA motif-anchored coverage and GAA-centered trimer-context enrichment are shown in Supplementary Figs. 31 and 32. Cross-fitted evaluation is reported in Fig. 4C–F in the main text.

### 5. Candidate-library generation and frozen-model screening

#### 5.1 Candidate-library generation

After the model and selection rules were frozen, we generated 500,000 candidate DNA sequences with lengths uniformly sampled from 18 to 25 nucleotides, using independent and uniform sampling from A, C, G and T with no motif constraints or model feedback. For each candidate, we computed the four physical features ($\Delta\varepsilon$, $\Delta$stack, $stack_{mean}$ and $C_{markov}$) using the frozen feature pipeline, in which $C_{markov}$ was derived from a second-order Markov model fitted to all 86 labeled training sequences. The frozen random forest returned $P_{(multiphase)}$, and binary calls were made at a threshold of 0.50. Tree-based SHAP decomposed each probability into local contributions from the four inputs. The screening record retained probability, fold-based uncertainty, physical-range and novelty fields, where uncertainty denotes the standard deviation across context-model folds rather than a predictive confidence interval. Library quality-control measures, including realized size, uniqueness, length distribution and nucleotide composition, were assessed post hoc (Table S5.1).

**Table S5.1** Candidate-library quality control

| Item | Result |
|---|---|
| Candidate sequences | 500,000 |
| Unique sequences | 500,000 |
| Length range | 18-25 nt |
| Total bases | 10,750,802 |
| Length distribution | 18 nt: 62,461; 19 nt: 62,301; 20 nt: 62,326; 21 nt: 62,701; 22 nt: 62,646; 23 nt: 62,568; 24 nt: 62,657; 25 nt: 62,340 |
| Base distribution | A: 2,688,814 (25.01%); C: 2,687,443 (25.00%); G: 2,687,059 (24.99%); T: 2,687,486 (25.00%) |

#### 5.2 Screening of Seg1-Seg6 candidates

For candidate selection, we adopted a paired design organized along three mechanistic axes: $\Delta\varepsilon$ for heterotypic interactions, $\Delta$stack for homotypic stacking and $C_{markov}$ for sequence context, with $stack_{mean}$ retained as a control feature. Candidates first satisfied sequence acceptability, training-range and maximum-homopolymer-run constraints. Each pair required a target-axis contrast of at least 0.35 training-set interquartile range (IQR) and control-feature differences below 0.80 IQR. Pair utility was defined as the normalized target contrast divided by one plus the RF-importance-weighted background mismatch. We selected three non-overlapping pairs that maximized total utility, yielding two representatives per mechanistic axis for experimental validation. Experimental labels were used only after the six sequences had been frozen (Table S5.2).

The full candidate library and detailed per-sequence scoring records are archived separately; summary statistics and the six final validation records are provided in this supplementary package. The complete sequences of Seg1–Seg6, along with their experimental phase assignments, frozen-model scores and local SHAP values, are listed in Tables S5.3–S5.5. The screening workflow is presented in Figure S5.1 below. The validation images are presented in Fig. 4G and H of the main text.

**Table S5.2** Candidate-screening data inventory

| Data component | Content | Purpose |
|---|---|---|
| Candidate library | 500,000 sequences | Random sequence generation |
| Screening record | 500,000 scored rows | Frozen features, probabilities and SHAP-based cue attribution |
| Seg1–Seg6 prediction record | Six records | Frozen-model scores and phase predictions |
| Seg1–Seg6 SHAP record | Six records | Local SHAP values and probability reconstruction |

**Table S5.3** Seg1–Seg6 sequences and experimental phases

| Sequence ID | DNA sequence | Experimental phase |
|---|---|---|
| Seg1 | ACGCGCGAACTGAAACTGAACGA | Multiphase |
| Seg2 | GAGAACGGAAAGTAGTAGAATAC | Multiphase |
| Seg3 | TCGGAATAATCGAACAGAATC | Single-phase |
| Seg4 | GAGAACATGAACAGGAAATAGAAC | Multiphase |
| Seg5 | AGAAAGAAGCGAATCTGAAAA | Multiphase |
| Seg6 | ACTGAAATGGAATCTGAAGAGAATG | Single-phase |

**Table S5.4** Seg1–Seg6 frozen-model scoring

| Sequence ID | Experimental phase | $P_{(multiphase)}$ | Prediction | Δε | Δstack | DNA stacking mean | $C_{markov}$ |
|---|---|---|---|---|---|---|---|
| Seg1 | Multiphase | 0.5601 | Multiphase | 9.577 | -11.550 | 0.828 | 0.051 |
| Seg2 | Multiphase | 0.9824 | Multiphase | 10.201 | -11.160 | 1.031 | 0.247 |
| Seg3 | Single-phase | 0.1097 | Single-phase | 7.491 | -8.260 | 0.876 | -0.244 |
| Seg4 | Multiphase | 0.7525 | Multiphase | 9.961 | -12.800 | 0.980 | -0.235 |
| Seg5 | Multiphase | 0.7182 | Multiphase | 8.998 | -12.665 | 0.988 | 0.162 |
| Seg6 | Single-phase | 0.4877 | Single-phase | 10.519 | -10.530 | 0.967 | 0.074 |

**Table S5.5** Seg1–Seg6 row-level SHAP data

| Sequence ID | $P_{(multiphase)}$ | $\Delta\varepsilon$ SHAP | $\Delta$stack SHAP | DNA stacking mean SHAP | $C_{markov}$ SHAP |
|---|---|---|---|---|---|
| Seg1 | 0.5601 | 0.134 | 0.023 | 0.035 | -0.137 |
| Seg2 | 0.9824 | 0.145 | 0.016 | 0.193 | 0.123 |
| Seg3 | 0.1097 | -0.220 | -0.026 | -0.050 | -0.098 |
| Seg4 | 0.7525 | 0.202 | -0.013 | 0.121 | -0.063 |
| Seg5 | 0.7182 | -0.027 | -0.018 | 0.251 | 0.008 |
| Seg6 | 0.4877 | 0.132 | 0.004 | 0.039 | -0.192 |

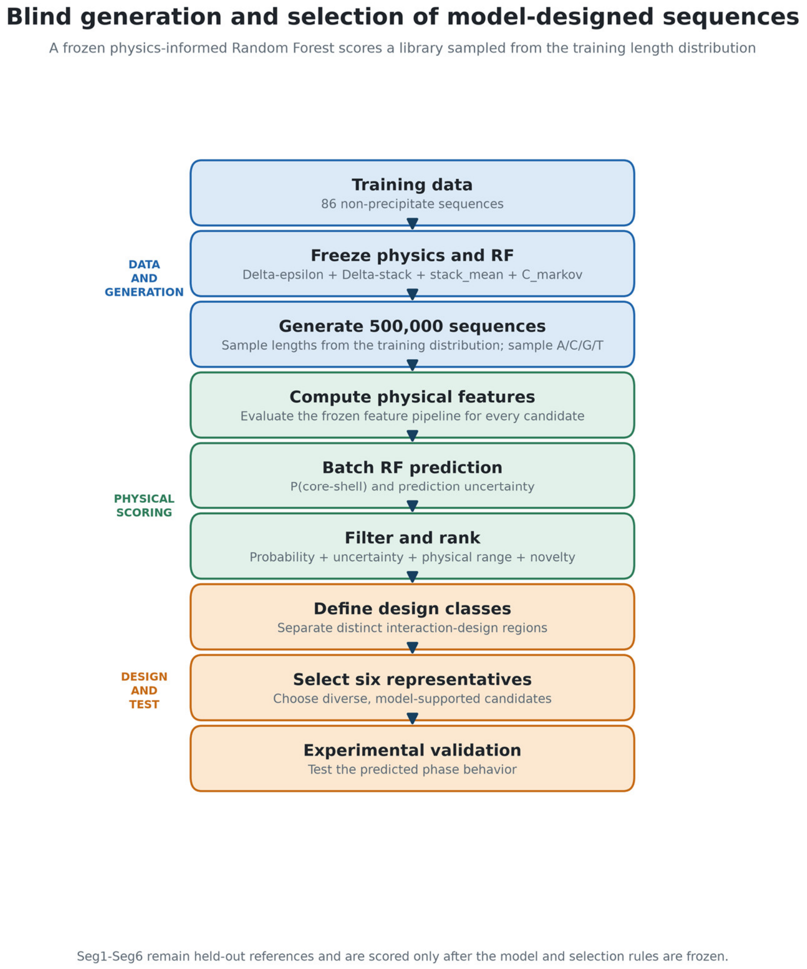


**Figure S5.1** Candidate generation and frozen-model screening workflow.

**6 Number space calculation for programmable multiphase condensate species**

To estimate the sequence space of our multiphase condensate library, we employed a combinatorial enumeration based on the motif $N_xGAAN_yGAAN_zGAAN_pGAAN_q$, where N represents any nucleotide (A, C, G, U/T) and $x \geq 1, y \geq 1, z \geq 1, p \geq 1,\ q \geq 0$ are integers.

For a total length of 20 nucleotides, the four fixed GAA blocks contribute 12 bases, leaving 8 variable positions, i.e.,

$$m = x + y + z + p + q = 8.$$

Under this constraint, exhaustive counting of all distinct length distributions (x, y, z, p, q) gave $M = 70$ combinations.

The total number of possible sequences is therefore

$$M \times 4^m = 70 \times 65{,}536 = 4{,}587{,}520.$$

Sampling 50 % and 10 % of this sequence diversity would require approximately 2,293,760 and 458,752 sequences, respectively. Applying the same method to lengths of 21, 22, 23, and 24 nt yields the library sizes summarized in the following table and in Supplementary Fig. 33.

| **Chain length** | **20** | **21** | **22** | **23** | **24** |
|---|---|---|---|---|---|
| $m$ | 8 | 9 | 10 | 11 | 12 |
| $M$ | 70 | 126 | 210 | 330 | 495 |
| $M \times 4^m$ | 4587520 | 33030144 | 220200960 | 1384120320 | 8304721920 |
| 50% Accuracy | 2293760 | 16515072 | 110100480 | 692060160 | 4152360960 |
| 10 % Accuracy | 458752 | 3303014.4 | 22020096 | 138412032 | 830472192 |

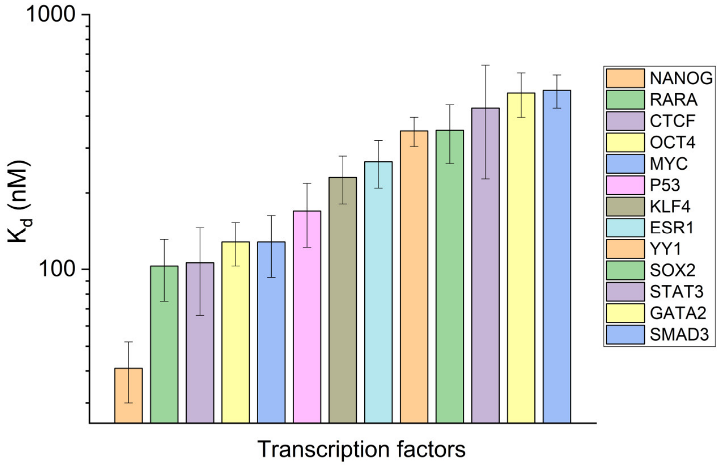


**Supplementary Fig. 1.** Binding affinities of various transcription factors with RNA. The binding affinities, ranging from 41 to 505 nM, were identified *in vitro*. All data were adapted and accessible from the referenced literature[7].

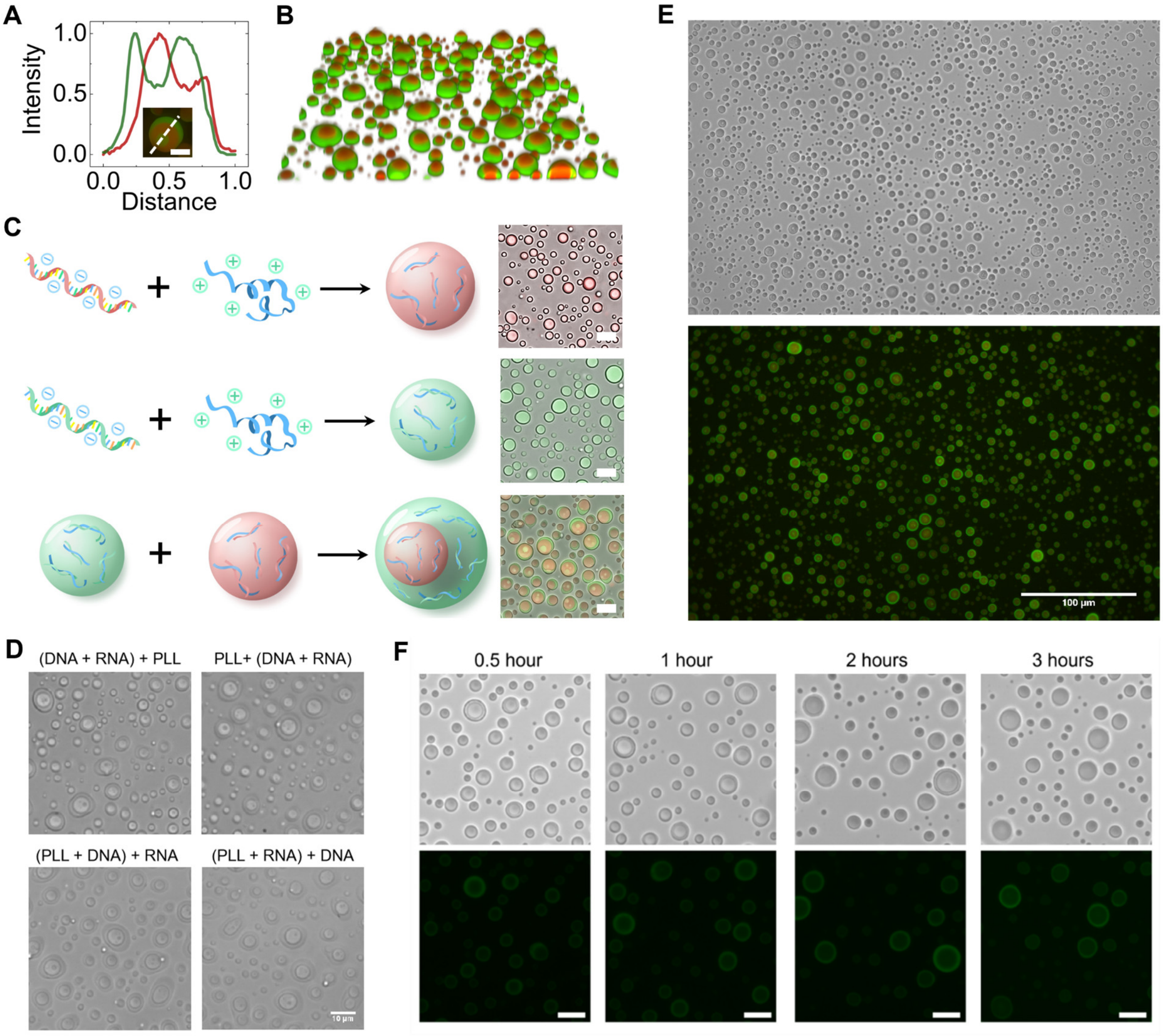


**Supplementary Fig. 2.** (A) Fluorescence intensity profile within a single multiphase droplet, illustrating the effective separation of DNA and RNA. Scale bar, 5 µm. (B) Stacked confocal microscopy image depicting multiphase droplets within a view box of 88.3 × 88.3 × 3.8 µm³ in size. (C) Mixing RNA or DNA with PLL forms single phase droplets, while mixing RNA-PLL droplets and DNA-PLL droplets forms multiple phase droplets. (D) The core-shell architecture does not depend on the order in which the components are mixed. (E) Multiphase condensates assemble within approximately 3 minutes after mixing PLL with a solution containing both DNA and RNA. These condensates feature tiny DNA-rich droplets trapped within an RNA-rich core, which is itself surrounded by a DNA-rich shell. (F) The core-shell-structured multiphase condensates remain stable for incubation times ranging from 30 minutes to 3 hours. Scale bar: 10 µm, unless otherwise specified in panel (E).

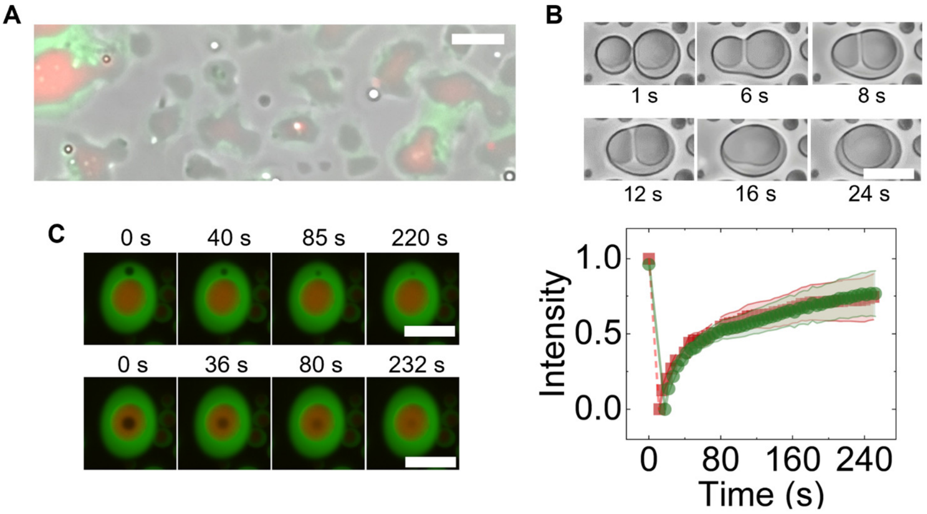


**Supplementary Fig. 3.** (A) Wetting behavior of multiphase droplets on a glass slide. (B) Coalescence process of two multiphase droplets. (C) Fluorescence Recovery After Photobleaching (FRAP) trajectory showing the liquid-like properties of the DNA-rich shell (green) and RNA-rich core (red) within the multiphase droplet. Error bars represent standard deviation from five individual repeats. Scale bar, 10 µm.

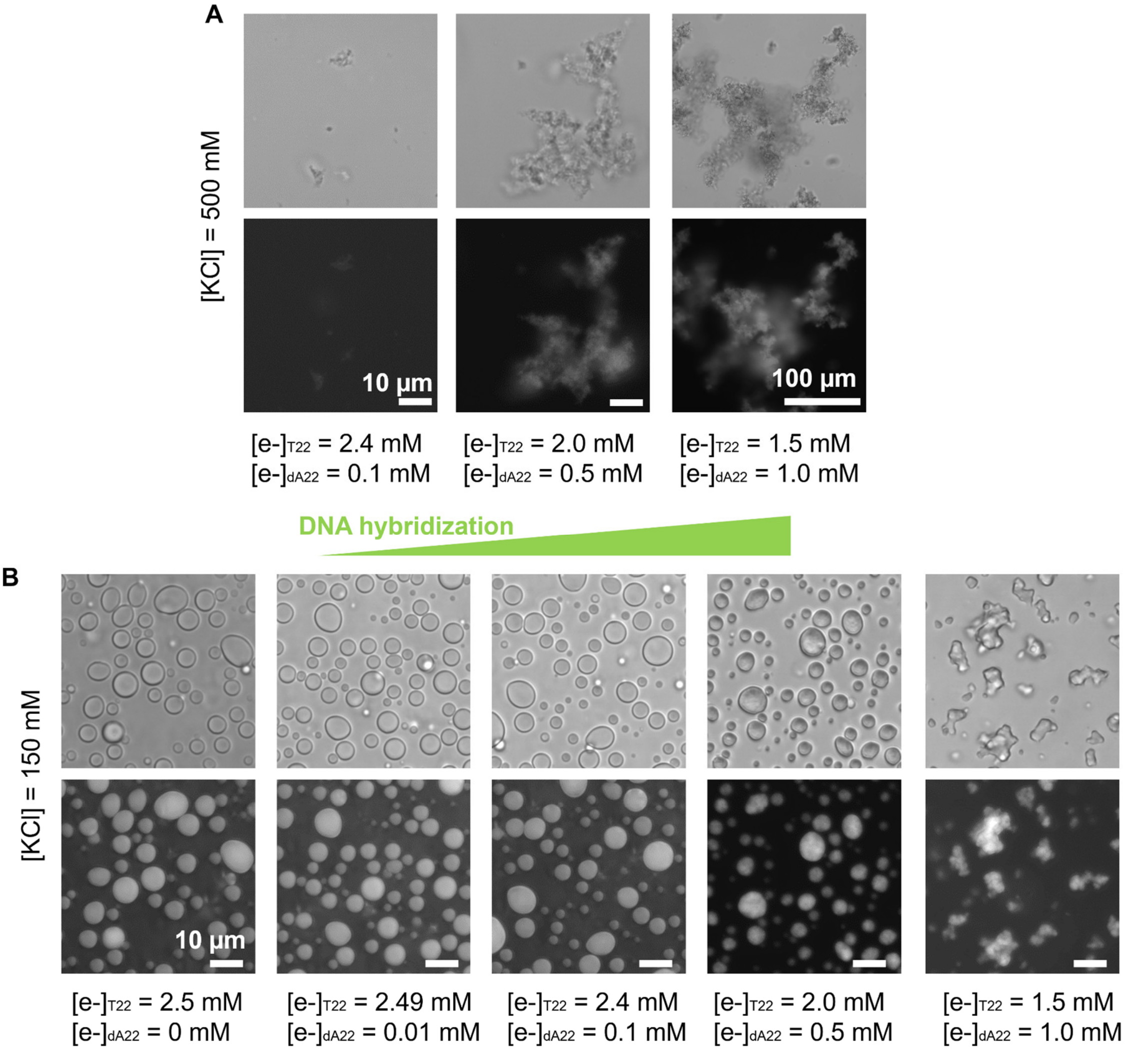


**Supplementary Fig. 4.** Increasing the DNA hybridization degree does not necessarily form multiphase condensates. (A) At high salt concentrations, solid-like precipitates formed even when only a small fraction (4% of the total charge concentration of DNA) of dA22 was added. With the addition of 40% dA22 (of the total charge concentration of DNA), large-scale precipitates were observed. (B) At low salt concentrations, an increase in the dA22 fraction resulted in a transition from liquid droplets to solid-like precipitates. The charge concentration of PLL, $U_{22}$, along with the total charge concentration of $T_{22}$ and $dA_{22}$, were maintained at 2.5 mM. FAM-labelled $T_{22}$ (2 µM) was used for fluorescence imaging and characterizing DNA distribution.

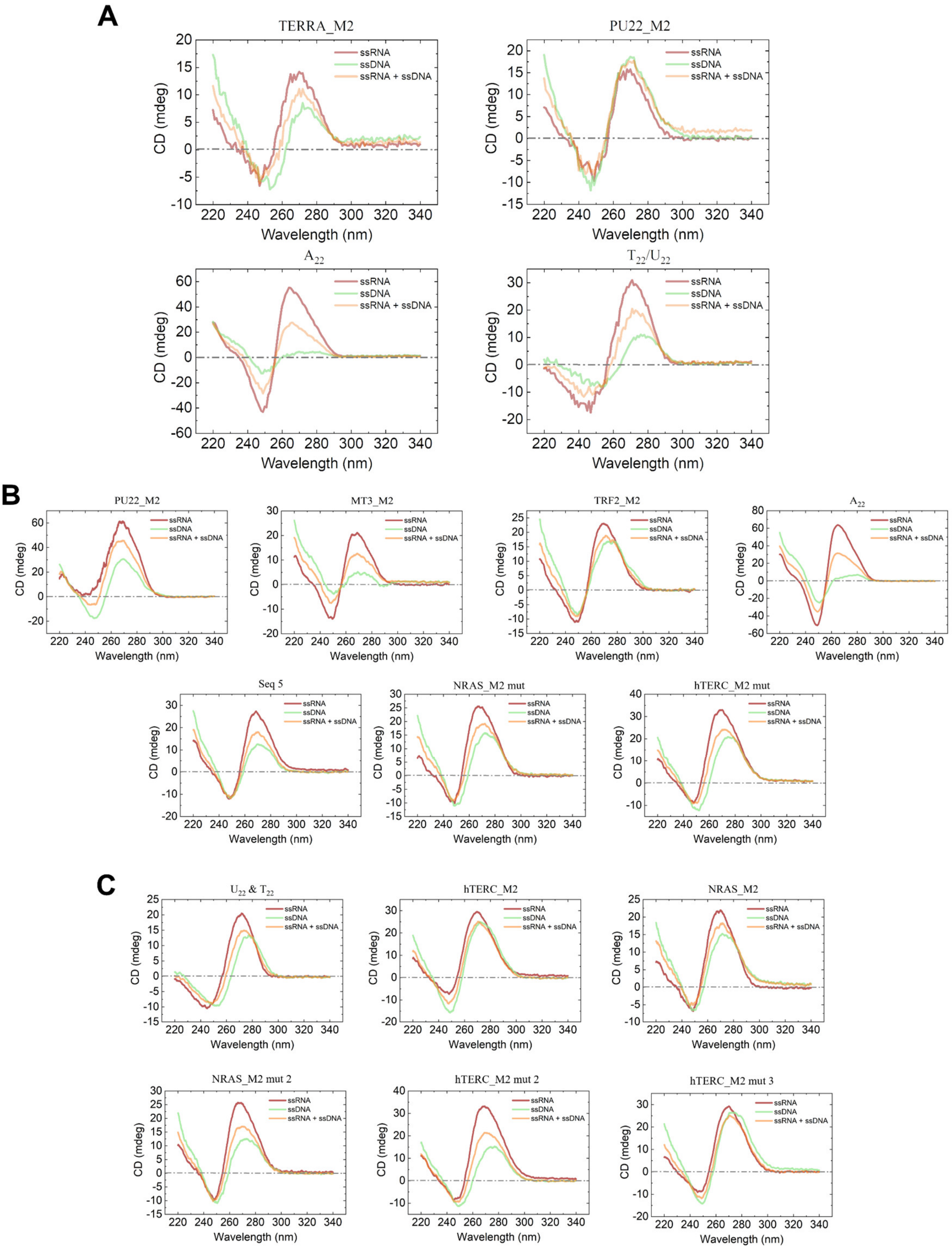


**Supplementary Fig. 5.** (A) Circular dichroism (CD) spectra of DNA and/or RNA oligonucleotides in 150 mM KCl buffer without PLL addition. (B-C) Circular dichroism (CD) spectroscopic characterization of oligonucleotides in condensate-forming buffered solutions. For each sequence, CD spectra were acquired in the presence of 500 mM KCl and 40 mM Tris. Panels (B) and (C) display spectra for oligonucleotides that successfully undergo multiphase coacervate formation and those that do not, respectively.

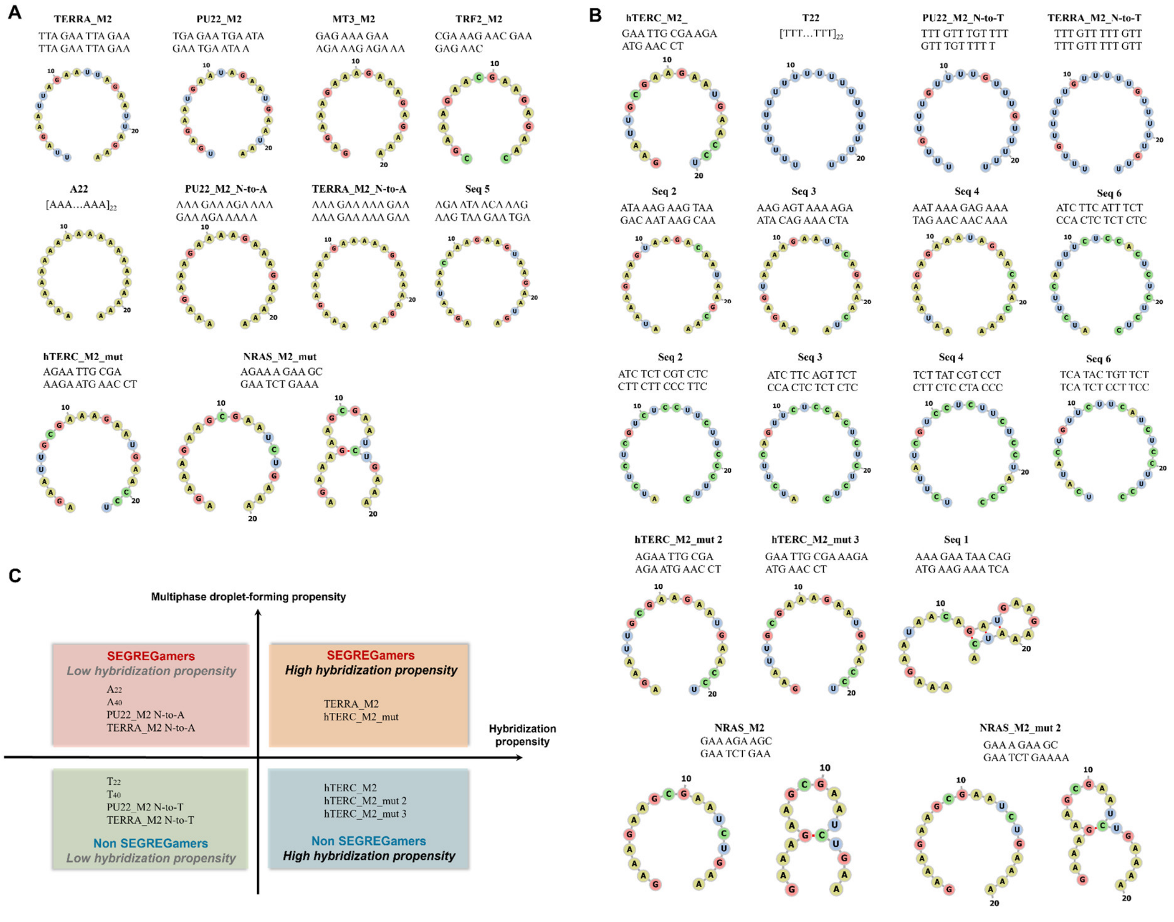


**Supplementary Fig. 6.** (A) Predicted secondary structures (from the RNAfold web server) of typical oligonucleotide sequences that can form DNA/RNA/PLL multiphase condensates. (B) Predicted secondary structures of typical oligonucleotide sequences that can form DNA/RNA/PLL multiphase condensates. (C) Weak correlation between degree of base pairing and multiphase condensate formation for representative DNA/RNA oligonucleotides.

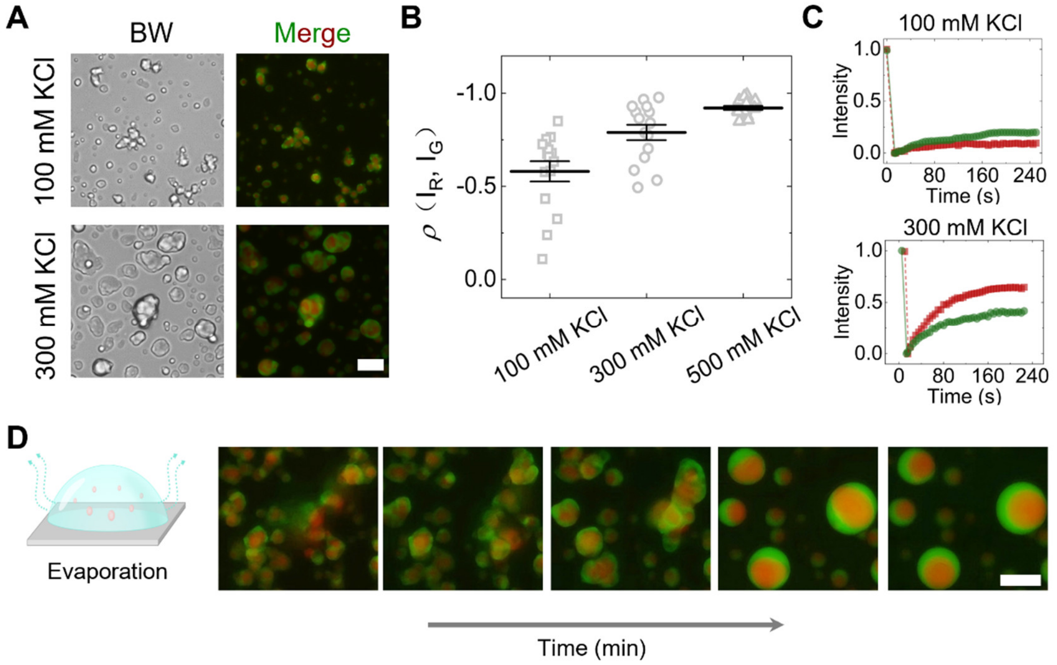


**Supplementary Fig. 7.** (A) Multiphase gel-like aggregates formed by DNA/RNA/PLL complex coacervation at 100 and 300 mM KCl, featuring an RNA-rich core surrounded by a DNA-rich shell. (B) Covariance of DNA and RNA fluorescence intensity distribution in condensates formed at varying salt concentrations. (C) FRAP trajectory of DNA- and RNA-rich domains within the aggregates. (D) Evaporation-induced solid-to-liquid transition of multiphase condensates. Scale bar, 10 µm.

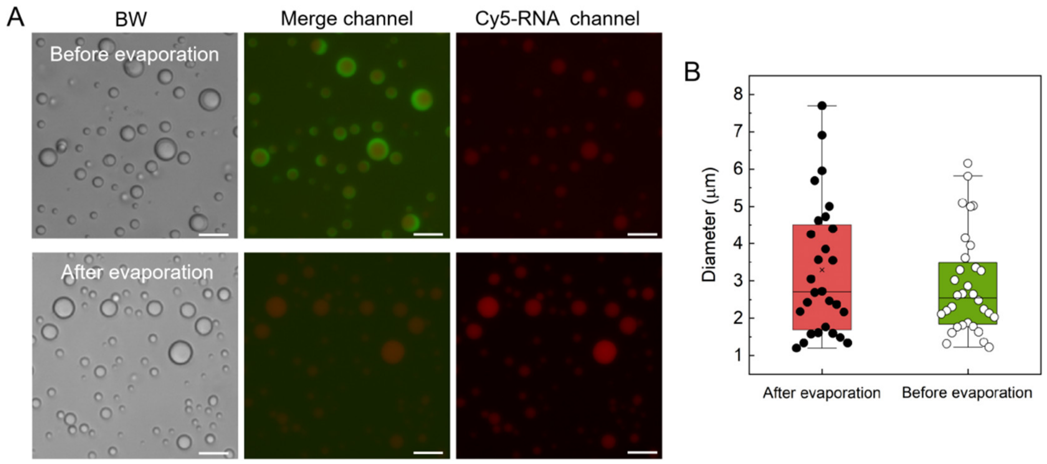


**Supplementary Fig. 8.** Evaporation-induced size change of RNA droplets. (A) Representative micrographs of Cy5-labeled RNA droplets pre- and post-evaporation. Scale bar: 10 µm. (B) Quantification of droplet diameters corresponding to the images in (A). This size increase upon evaporation is likely due to coalescence events, promoted by evaporation-induced increases in RNA and PLL concentration and a concomitant rise in salt concentration that reduces electrostatic attraction, thereby softening the droplets and facilitating fusion.

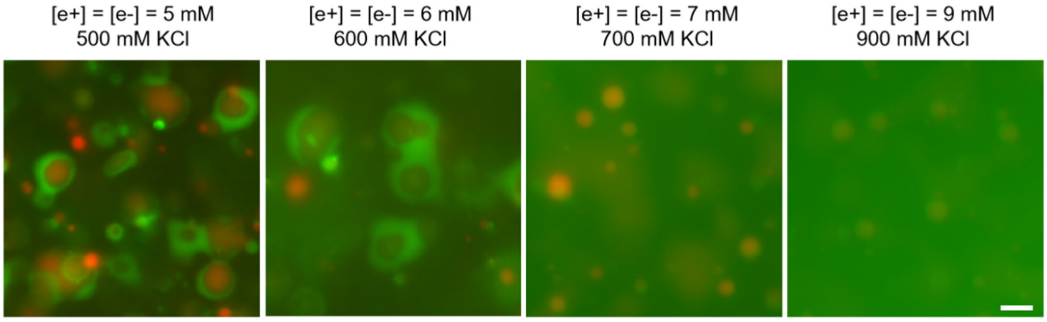


**Supplementary Fig. 9.** Dissolution of multiphase condensates. A condensate with a DNA-rich shell and an RNA-rich core gradually dissolves into an RNA-only condensate with increasing concentrations of DNA, RNA, PLL, and KCl. Scale bar: 10 µm.

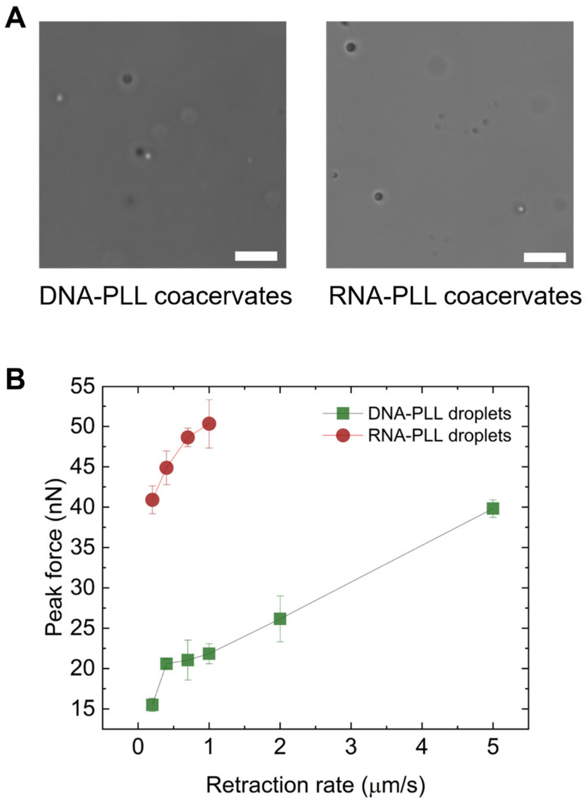


**Supplementary Fig. 10.** (A). Representative images of DNA-PLL and RNA-PLL coacervate droplets prepared for CP-AFM imaging. Droplets were formed at stoichiometric charge concentration of [e+] = [e-] = 0.3 mM for each component in the presence of 500 mM KCl. (B) Summary of peak forces of the force distance curves during retraction in CP-AFM measurements. Error bars represent the standard deviation of 20 measurements.

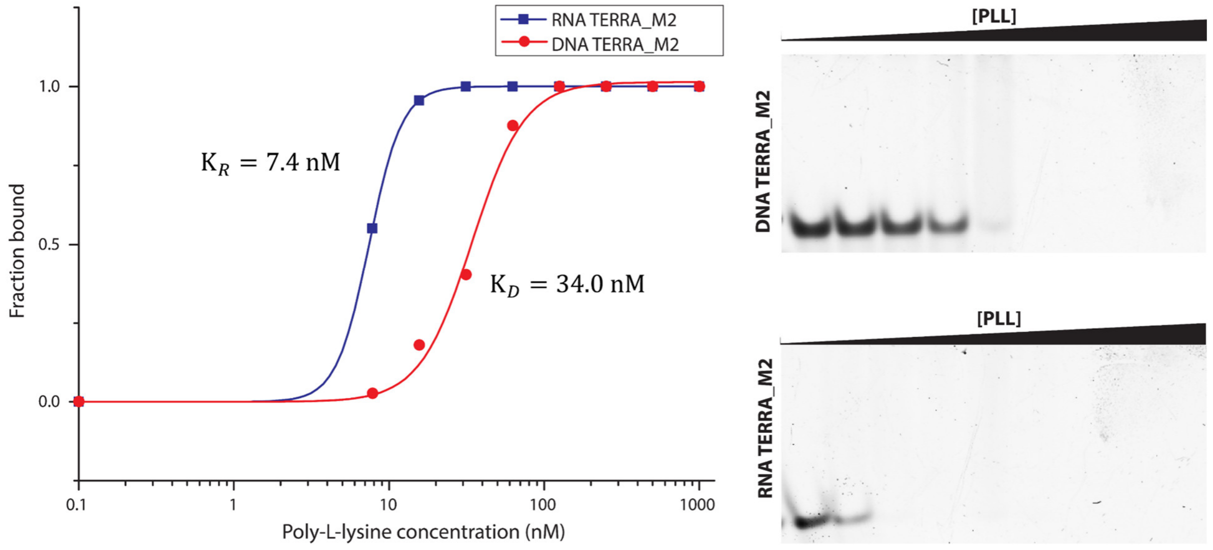


**Supplementary Fig. 11.** Binding constants between DNA/PLL and RNA/PLL. dTERRA_M2 and rTERRA_M2 bond to PLL with dissociation constants ($K_R$ and $K_D$) of 34.0 nM and 7.4 nM, respectively.

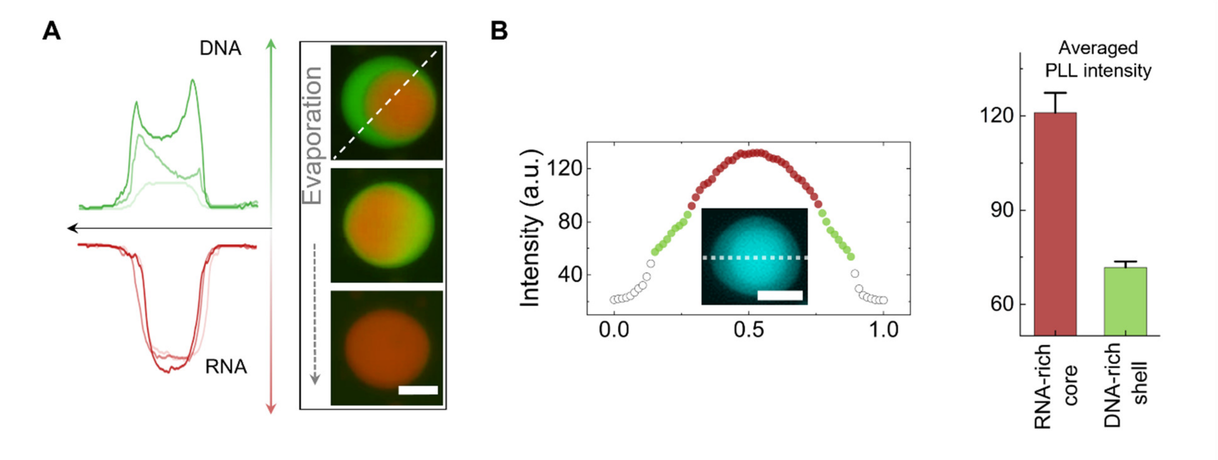


**Supplementary Fig. 12.** (A) Distribution of fluorescently labelled DNA and RNA during the evaporation of multiphase droplets. The initial salt concentration is 500 mM. (B) Higher PLL concentration in RNA-rich core. Scale bar, 5 µm.

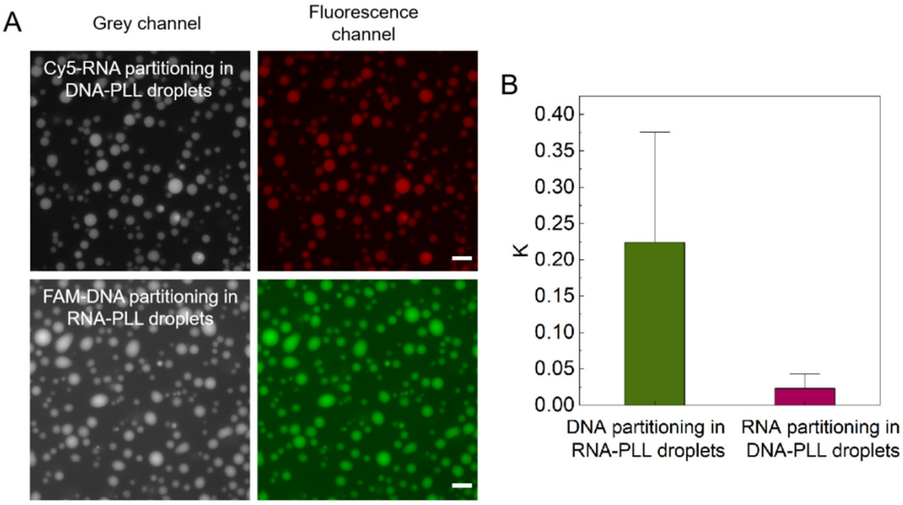


**Supplementary Fig. 13.** (A) Partitioning of fluorescently labeled RNA into DNA-PLL coacervate droplets (upper panel) and fluorescently labeled DNA into RNA-PLL coacervate droplets (lower panel). Scale bar: 10 µm. (B) Partitioning coefficients of DNA and RNA analyzed from micrographs in (A).

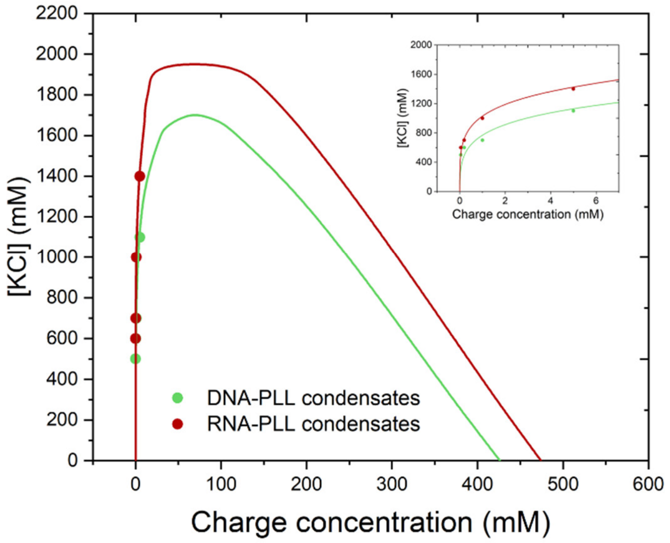


**Supplementary Fig. 14.** Experimental and theoretical Binodal curves of condensates formed by DNA-PLL and RNA-PLL condensates. The theoretical curves are solved using parameters: $N_n = 24$, $N_p = 240$, $\alpha = 1.1$ for RNA-PLL condenses, as well as $= 1.0$ for DNA-PLL condenses. Insert: the left arm of the phase diagram. A detailed derivation was provided in Supplementary Text 3.

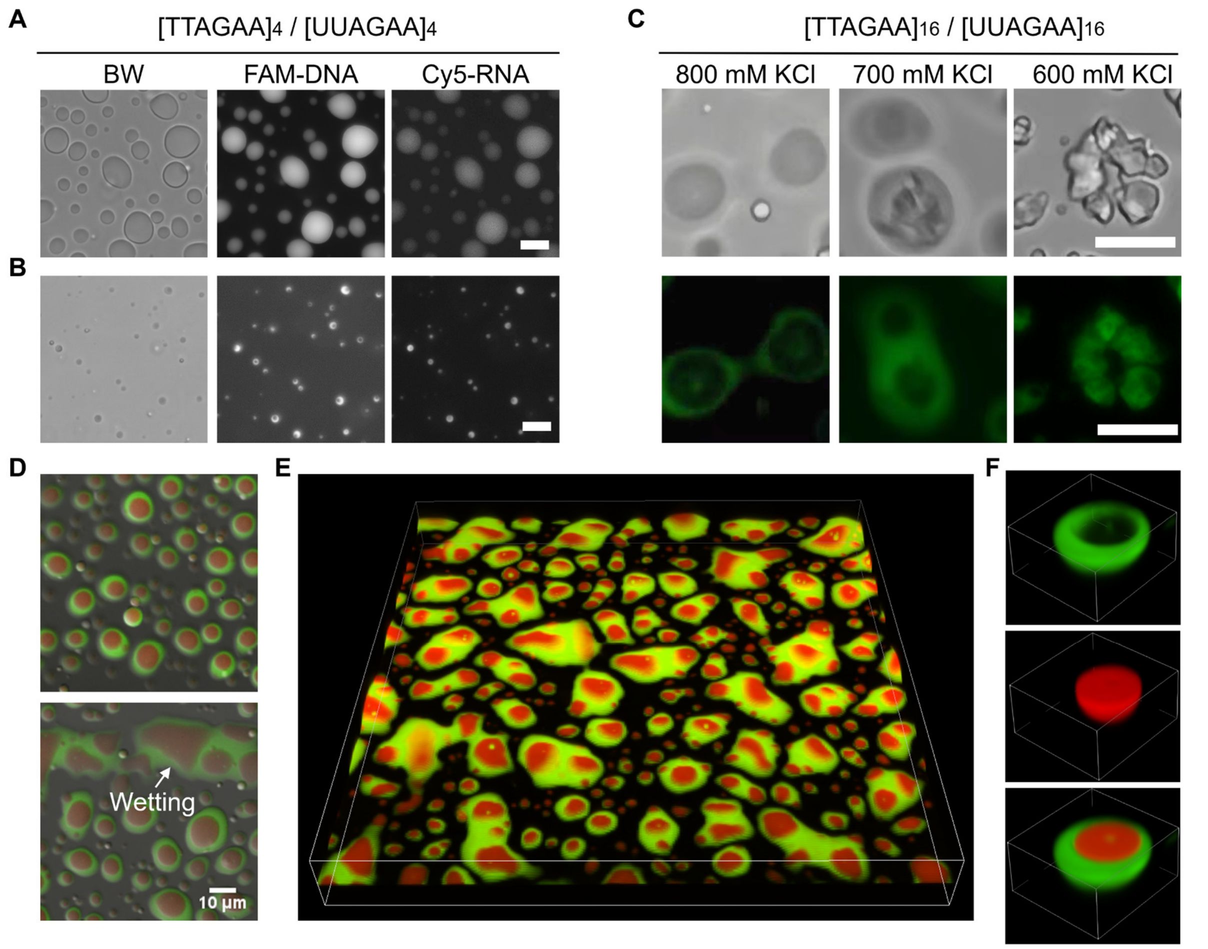


**Supplementary Fig. 15.** (A) Single-phase droplets form when RNA concentration is significantly lower than DNA, yet still containing both DNA and RNA components. Droplets were prepared at the charge concentration of [e+] = [e-] = 5 mM in 500 mM KCl buffer, with $[e\text{-}]_{DNA}$ = 4.5 mM and $[e\text{-}]_{RNA}$ = 0.5 mM. (B) In conditions of low charge concentration, multiphase droplets persist, exhibiting a DNA-rich shell and RNA-rich core. Droplets were prepared at the charge concentration of [e+] = [e-] = 1 mM in 500 mM KCl buffer, with $[e\text{-}]_{DNA}$ = 0.8 mM and $[e\text{-}]_{RNA}$ = 0.2 mM. (C) Multiphase droplets and gel-like structures formed by $[TTAGAA]_{16}/[UUAGAA]_{16}$. Droplets were prepared at the charge concentration of [e+] = [e-] = 5 mM in 800 mM KCl buffer, with $[e\text{-}]_{DNA}$ = 2.5 mM and $[e\text{-}]_{RNA}$ = 2.5 mM. (D) Fluorescence imaging of droplets formed by $[UUAGAA]_{16}$-$[TTAGAA]_{16}$-PLL phase separation. (E) Large scale 3D confocal image of multiphase DNA/RNA/PLL droplets in panel (D). An observation box of $92.6 \times 92.6 \times 4.2\ \mu m^3$ is shown. (F) 3D confocal image of multiphase DNA/RNA/PLL droplets prepared using $[UUAGAA]_{16}$-$[TTAGAA]_{16}$-PLL phase separation. FAM-$[TTAGAA]_4$ and Cy5-$[UUAGAA]_4$ were used at 2 µM. An observation box of $10.2 \times 10.2 \times 4.2\ \mu m^3$ is shown. Scale bar, 10 µm.

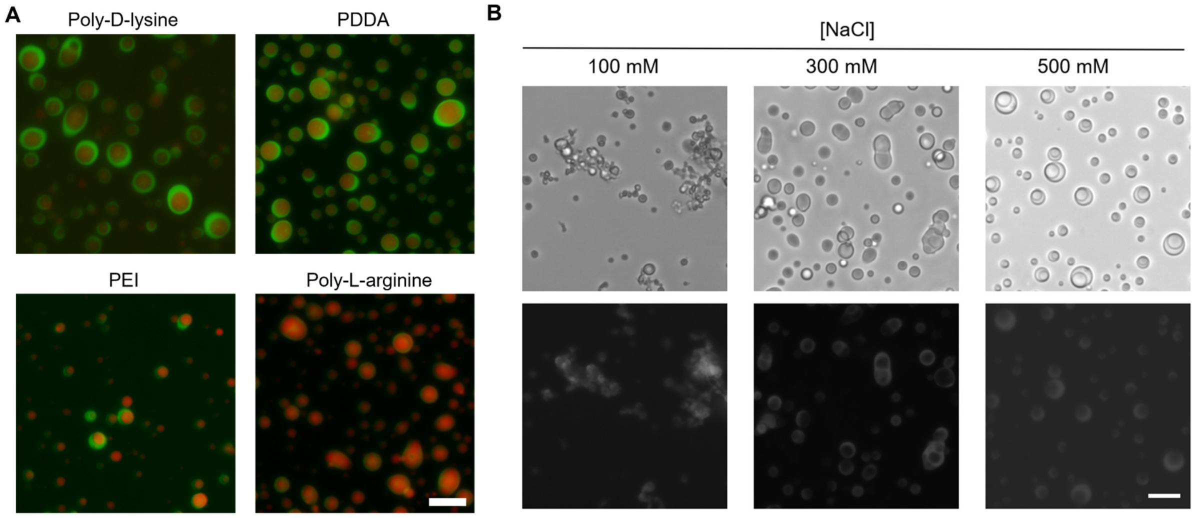


**Supplementary Fig. 16.** Formation of Multiphase Droplets with Different Cationic Polymers and Salt Conditions. (A) Formation of Multiphase Droplets with DNA/RNA Complexing with Various Cationic Polymers. Droplets were prepared at a charge concentration of [e+] = [e-] = 5 mM. For poly-D-lysine and PDDA, the [DNA]:[RNA] ratio was 1:1. For PEI and poly-L-arginine, the [DNA]:[RNA] ratio was 4:1. A buffer containing 900 mM KCl was used for the poly-L-arginine group. (B) Substituting KCl with NaCl retains the formation of multiphase droplets. Scale bar, 10 µm.

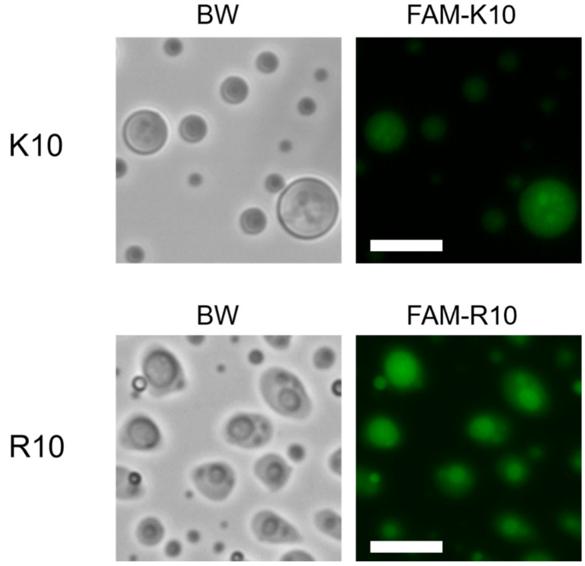


**Supplementary Fig. 17.** Formation of Multiphase Droplets with K10 and R10. Droplets were prepared at a charge concentration of [e+] = [e-] = 5 mM, [KCl] = 150 mM. Inhomogeneous distribution of K10 was shown within DNA-RNA-K10 condensates. Well-defined core-shell structures were identified within DNA-RNA-R10 condensates. Scale bar: 10 $\mu$m.

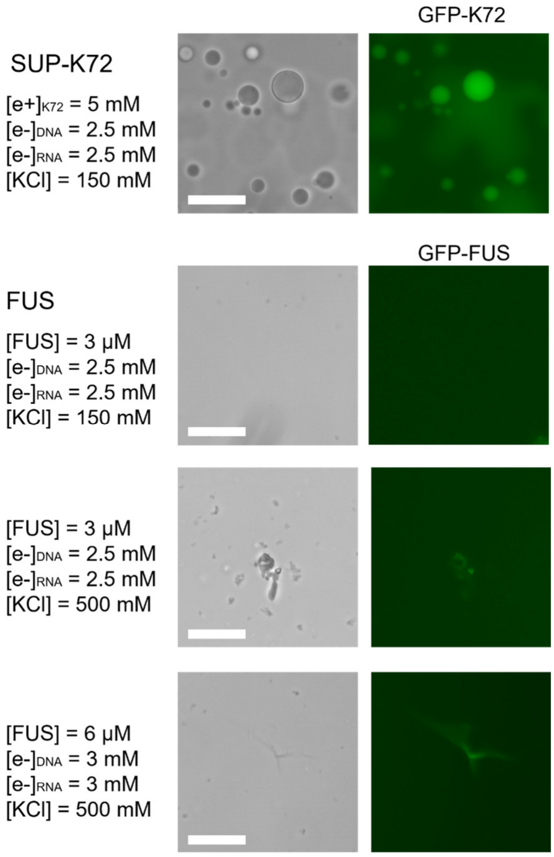


**Supplementary Fig. 18.** Replacing PLL with either SUP-K72 or FUS (Fused in Sarcoma) prevented the formation of multiphase condensates. For SUP-K72, single-phase droplets were formed. For FUS, no condensates or only solid-like condensates were formed. Scale bar: 10 $\mu$m.

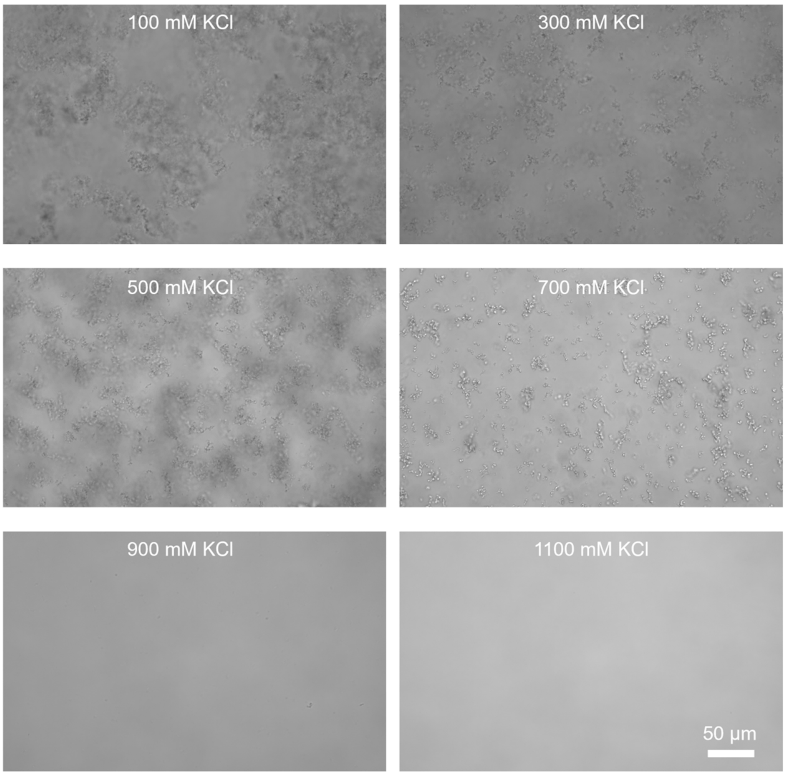


**Supplementary Fig. 19.** From 100 to 700 mM KCl conditions, methylated RNA-PLL coacervation consistently forms solid-like structures. $[UUAGAA]_4$ with 2'-O-methylation was used

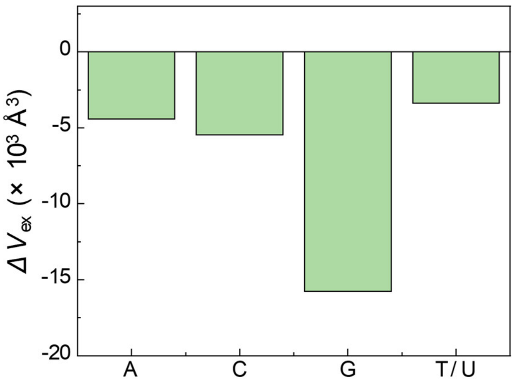


**Supplementary Fig. 20.** Excluded volume derived from PMFs between lysine-ribonucleotide and lysine-deoxyribonucleotide dimers.

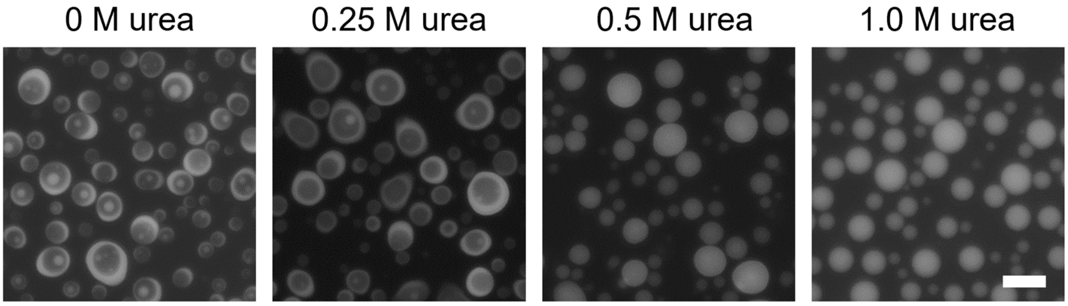


**Supplementary Fig. 21.** Addition of urea inhibits multiphase droplet formation. Scale bar, 10 μm.

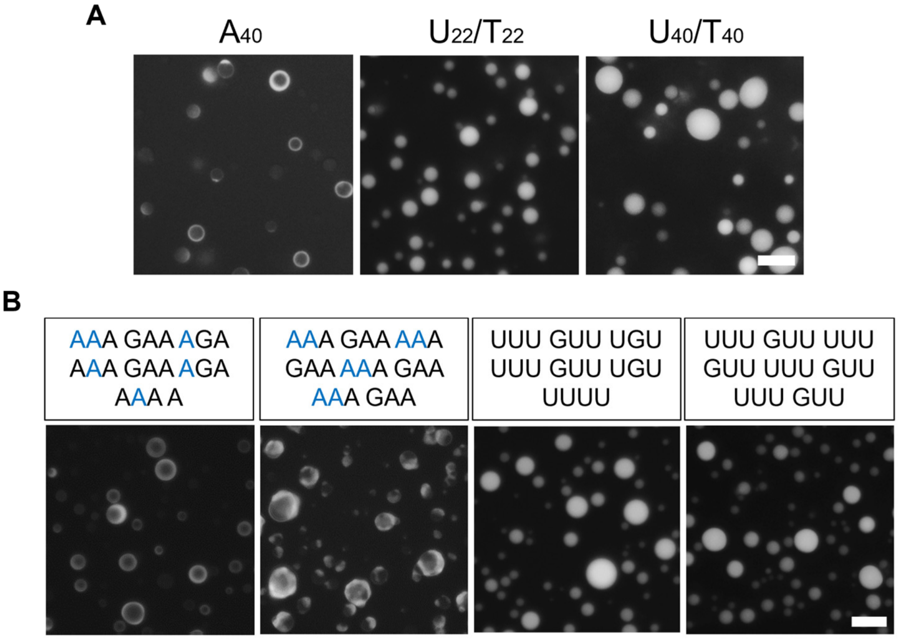


**Supplementary Fig. 22.** (A) $dA_{40}/rA_{40}$ forms multiphase droplets, while $U_{22}/T_{22}$ and $U_{40}/T_{40}$ form single-phase droplets. (B) N-to-A substitutions of TERRA_M2 and PU22_M2 at non-guanine sites result in multiphase droplets, whereas N-to-T/U substitutions do not. Scale bar, 10 μm.

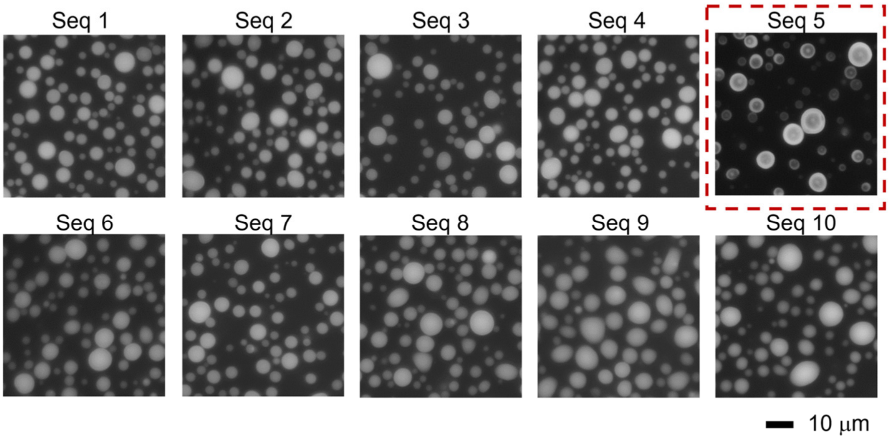


**Supplementary Fig. 23.** Droplets formed by custom-designed DNA and RNA oligonucleotides with varying adenine fractions. Scale bar, 10 µm.

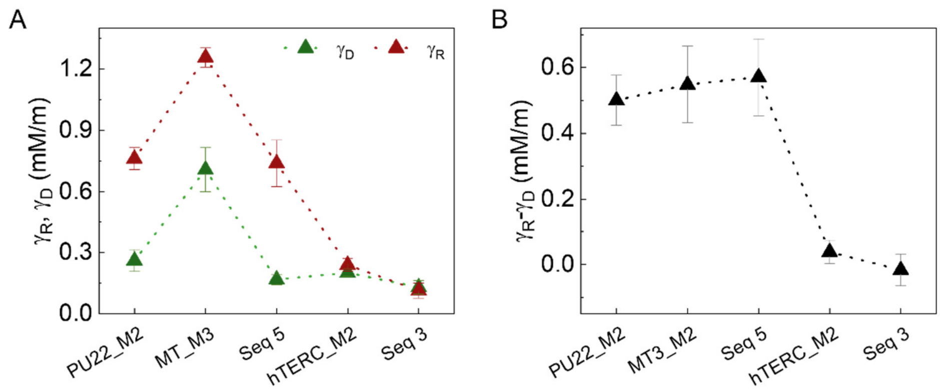


**Supplementary Fig. 24.** CP-AFM measurements were conducted to determine the interfacial tensions of DNA-PLL and RNA-PLL condensates formed with a range of sequences, including PU22_M2, MT3_M2, hTERC_M2, Seq3, and Seq5.

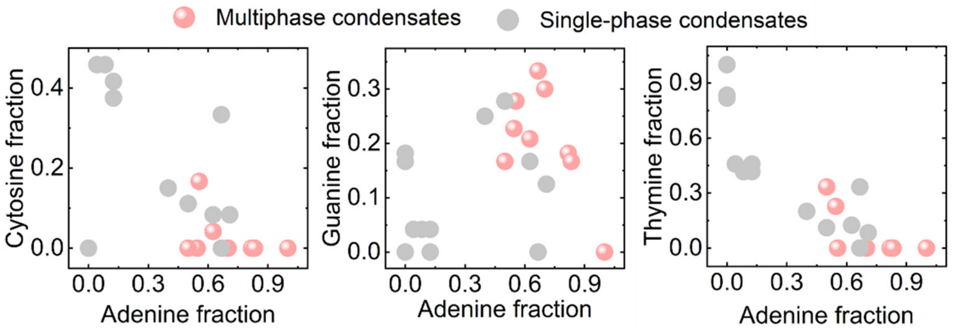


**Supplementary Fig. 25.** The fractions of cytosine (C), guanine (G), and pyrimidines (U and T) as a function of the fraction of adenine (A) for DNA/RNA sequences forming multiphase and single-phase condensates.

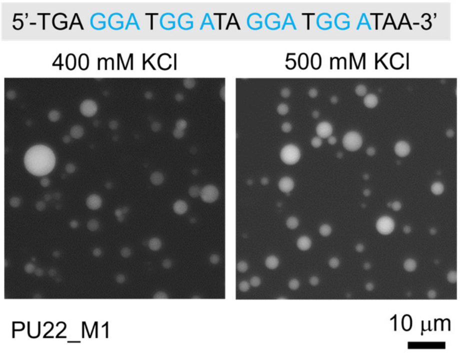


**Supplementary Fig. 26.** DNA and RNA of PU22_M1 with 'GGA' tracts do not form multiphase droplets.

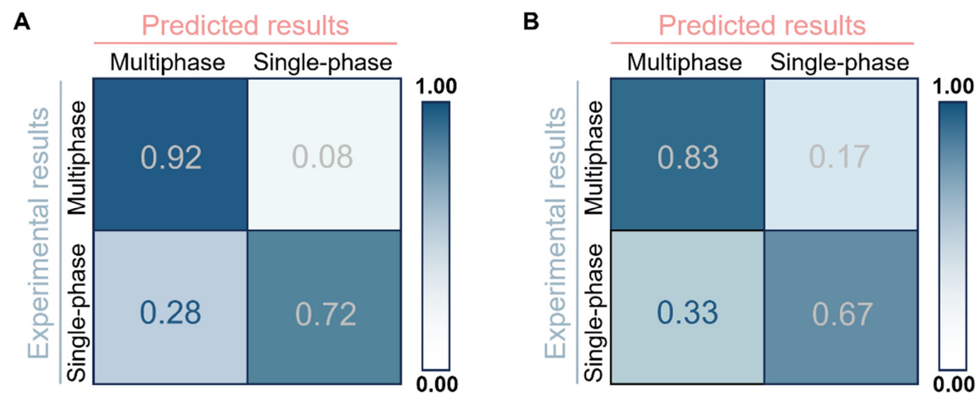


**Supplementary Fig. 27.** Predication performance of (A) the adenine fraction (> 50% or not) and (B) GAA tract numbers (> 3 or not) of oligonucleotides in forming multiphase droplets by the confusion matrix, derived from all tested DNA and RNA oligonucleotides listed in Supplementary Table S1 to S3. Specifically, in panel (A), for sequences that **can** form multiphase condensates (True Condition Positive): **92%** were correctly predicted by the model (True Positives, TP) and **8%** were incorrectly predicted (False Negatives, FN). For sequences that **cannot** form multiphase condensates (True Condition Negative): **72%** were correctly predicted by the model (True Negatives, TN) and **28%** were incorrectly predicted (False Positives, FP). Similar analysis was performed to derive the matrix in panel (B).

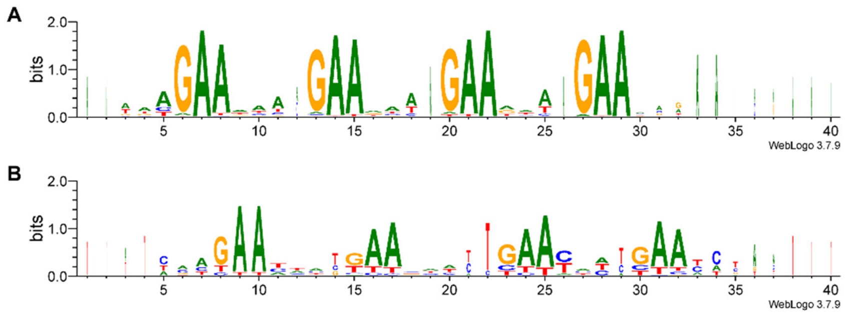


**Supplementary Fig. 28.** Consensus sequence motifs identified among all tested oligonucleotides that promote multiphase condensate formation (A) and single-phase condensate formation (N).

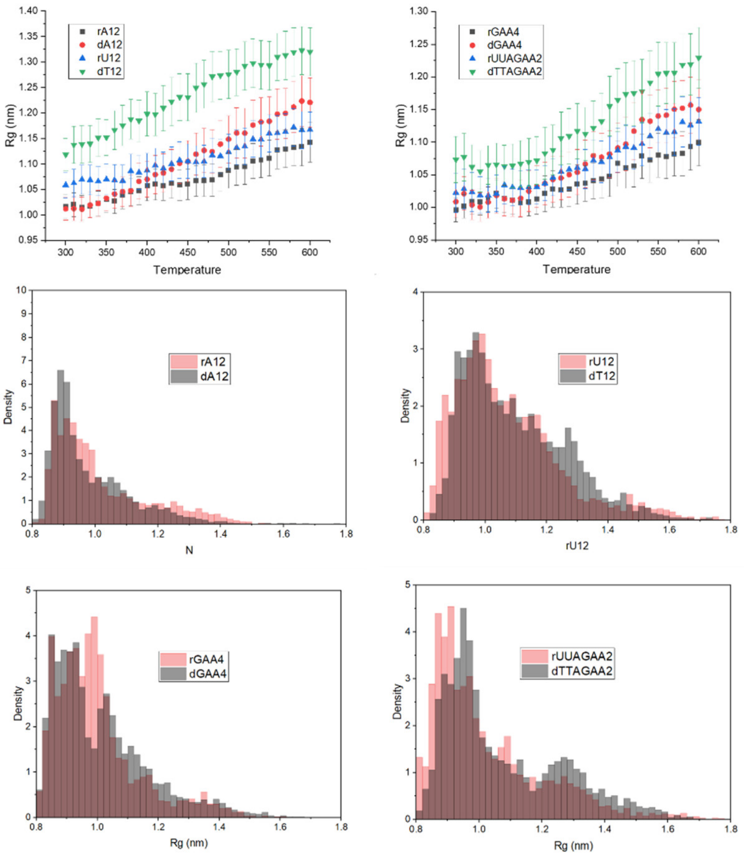


**Supplementary Fig. 29.** Radius of gyration ($R_g$) and its distribution for ssDNA and ssRNA with identical sequences. ssDNA/ssRNA pairs of $dA_{12}/rA_{12}$, $dT_{12}/rU_{12}$, $d[GAA]_4/r[GAA]_4$, as well as $d[TTAGAA]_2/r[UUAGAA]_2$ were investigated.

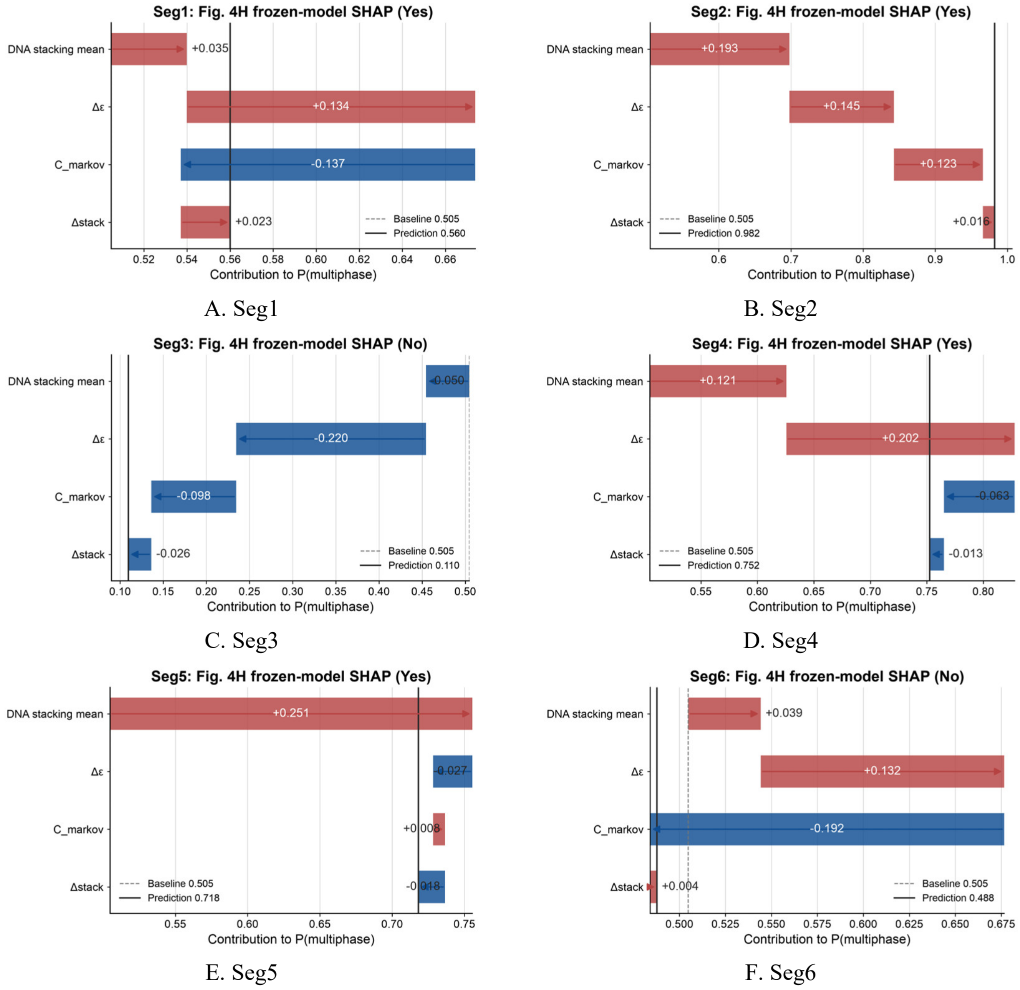


**Supplementary Fig. 30.** The panels A–F show the local SHAP decomposition for Seg1–Seg6. Each panel starts from the same baseline probability of 0.505; the feature contributions sum to $P_{(multiphase)}$. Red contributions increase the multiphase probability, whereas blue contributions decrease it. A probability of at least 0.50 maps to Multiphase, and a lower probability maps to Single-phase. The plotted feature order follows Fig. 4F in the main text: DNA stacking mean, Δε, $C_{markov}$ and Δstack.

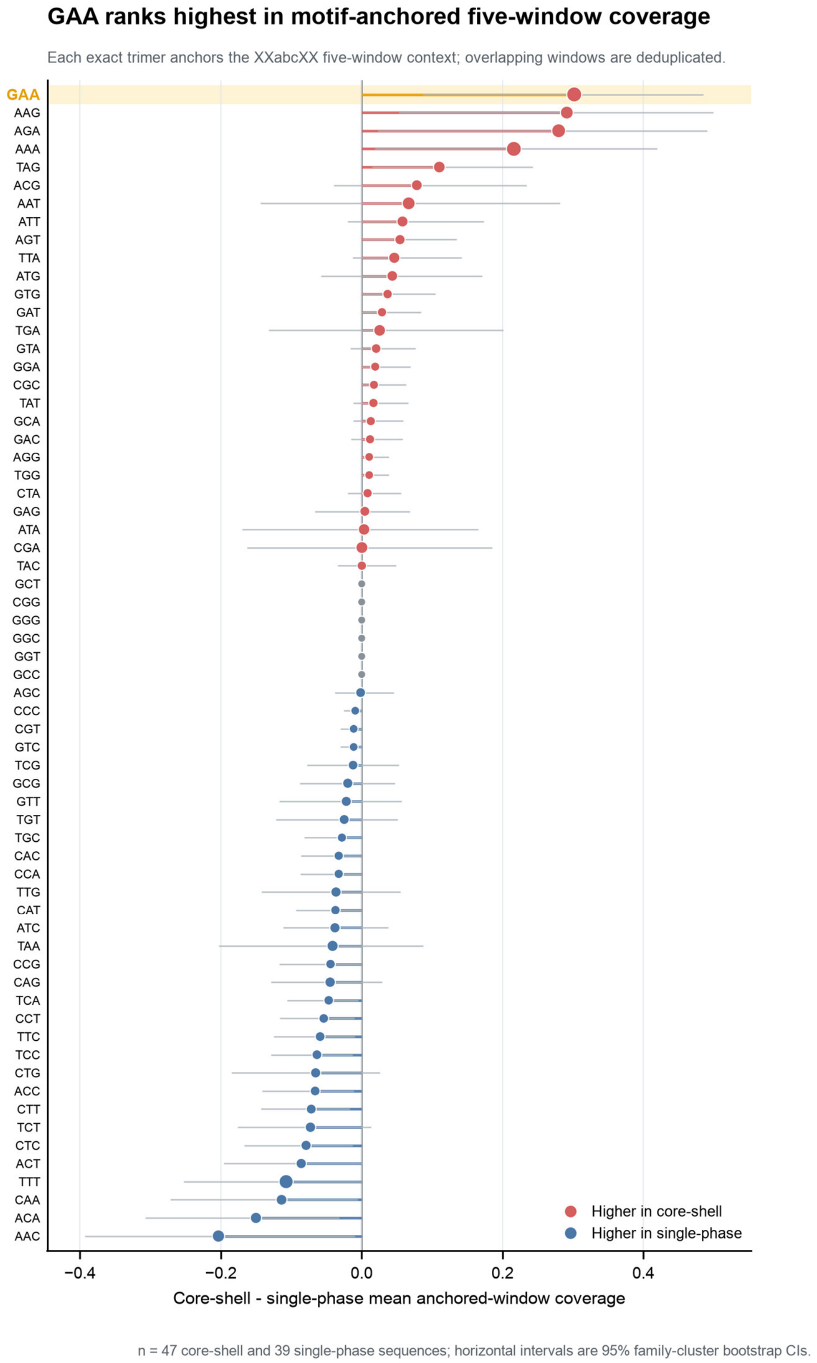


**Supplementary Fig. 31.** GAA shows the largest observed phase-class difference in motif-anchored five-window coverage. All 64 DNA trimers were evaluated using 86 non-precipitate sequence records (47 core-shell and 39 single-phase sequences). For each exact trimer occurrence, the two upstream, central and two downstream trimer windows were collected; overlapping windows were deduplicated. For GAA, mean coverage was 0.7815 in core-shell sequences and 0.4798 in single-phase sequences, giving a difference of 0.3017 (95% family-cluster bootstrap CI, 0.0863 to 0.4853). The 17 precipitate-forming sequences were excluded.

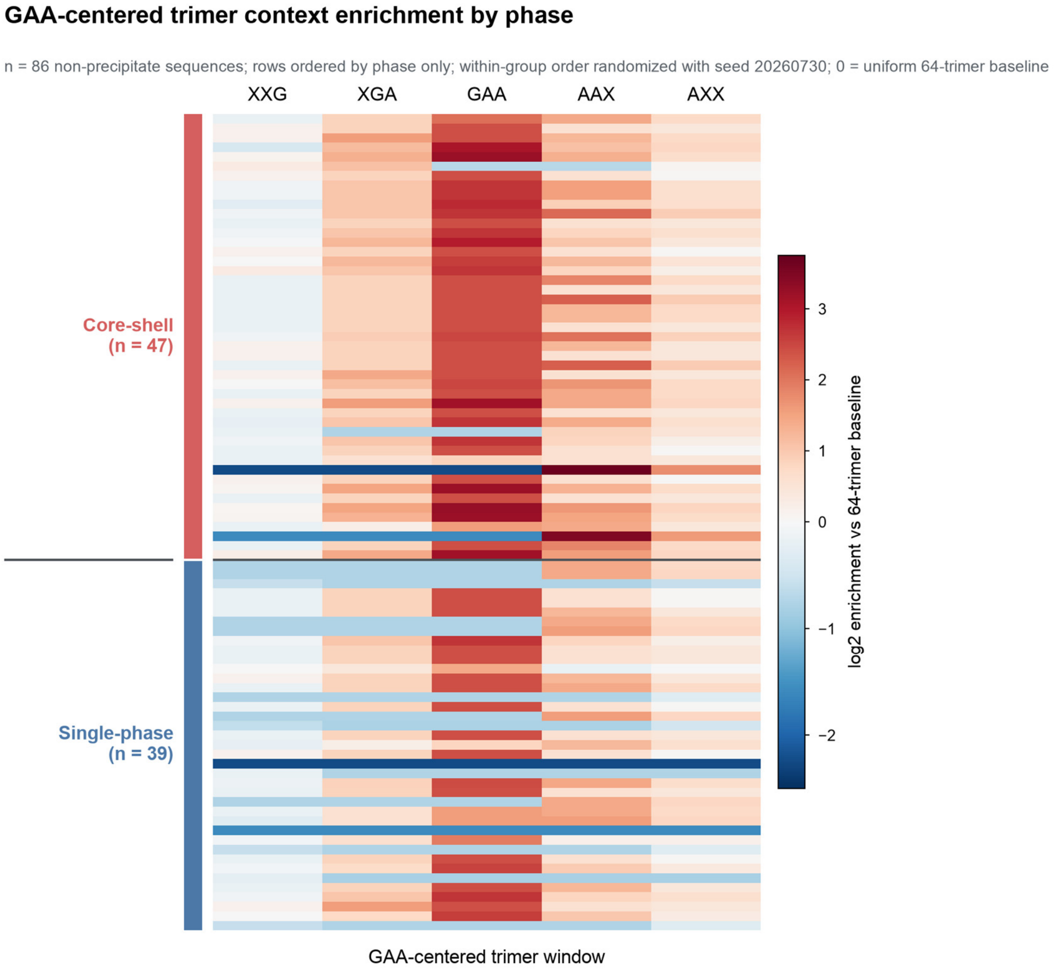


**Supplementary Fig. 32.** Global enrichment of GAA-related wildcard trimer classes by phase. The five columns show XXG, XGA, GAA, AAX and AXX classes containing 16, 4, 1, 4 and 16 concrete trimers, respectively. Here X denotes arbitrary nucleotide. Values are log2 enrichment relative to the uniform 64-trimer baseline after Jeffreys smoothing. The upper block contains 47 core-shell sequences and the lower block contains 39 single-phase sequences; within-class row order was randomized with seed. The 17 precipitate-forming sequences were excluded. This heatmap describes global sequence composition and does not represent exact GAA-site neighborhoods or an independent causal contribution to phase behavior.

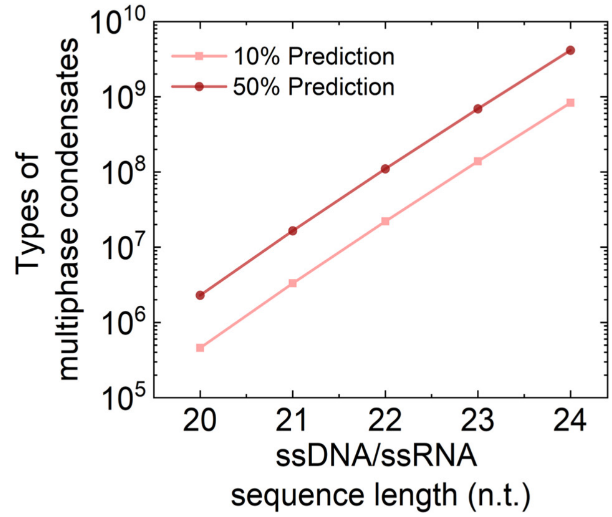


**Supplementary Fig. 33.** The diversity of multiphase condensates formed by the SEGREGamer library as a function of sequence length.

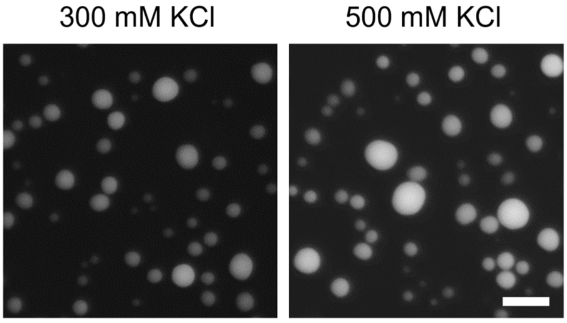


**Supplementary Fig. 34.** DNA/RNA oligonucleotide from the T7 promoter do not form multiphase droplets. Scale bar, 10 μm.

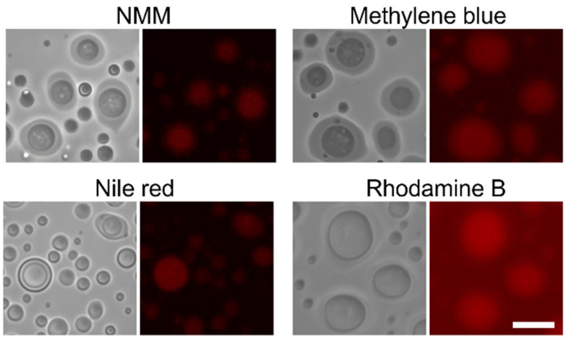


**Supplementary Fig. 35.** Partitioning of small molecule dyes within multiphase droplets. Scale bar, 10 μm.

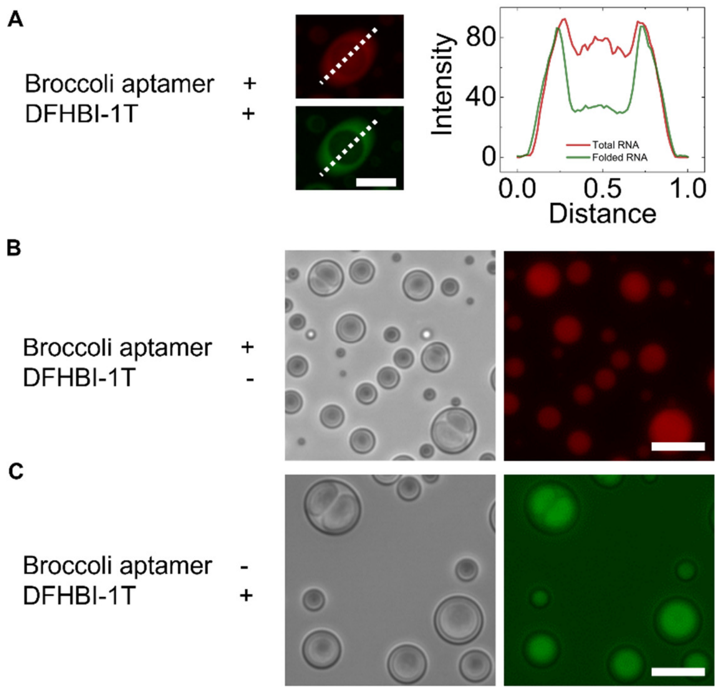


**Supplementary Fig. 36.** (A) In the presence of both DFHBI and RNA aptamers, the total RNA aptamer (red fluorescence) shows approximately equal distribution between the shell and the core phases, while the folded, DFHBI-bound form (green fluorescence) is enriched in the shell phase. (B) In the absence of DFHBI, the total RNA aptamer is enriched more in the core phase than in the shell phase. (C) In the absence of the RNA aptamer, DFHBI localizes into the core phase. Scale bar, 10 µm.

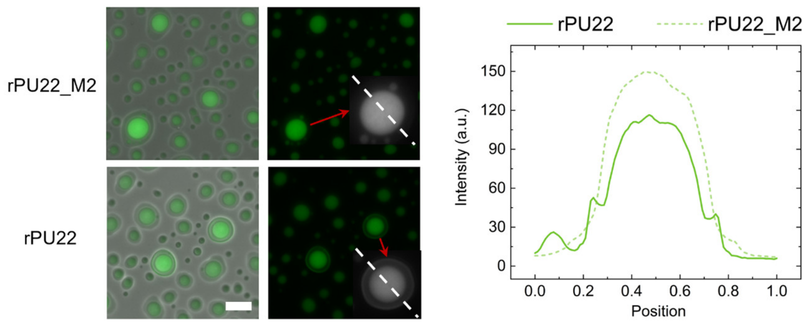


**Supplementary Fig. 37.** Partitioning of G-Quadruplex-Containing rPU22 and its Mutant rPU22_M2. The fluorescence profile within a single multiphase droplet indicates a greater exclusion of the more structured rPU22 by the core phase. Scale bar, 10 µm.

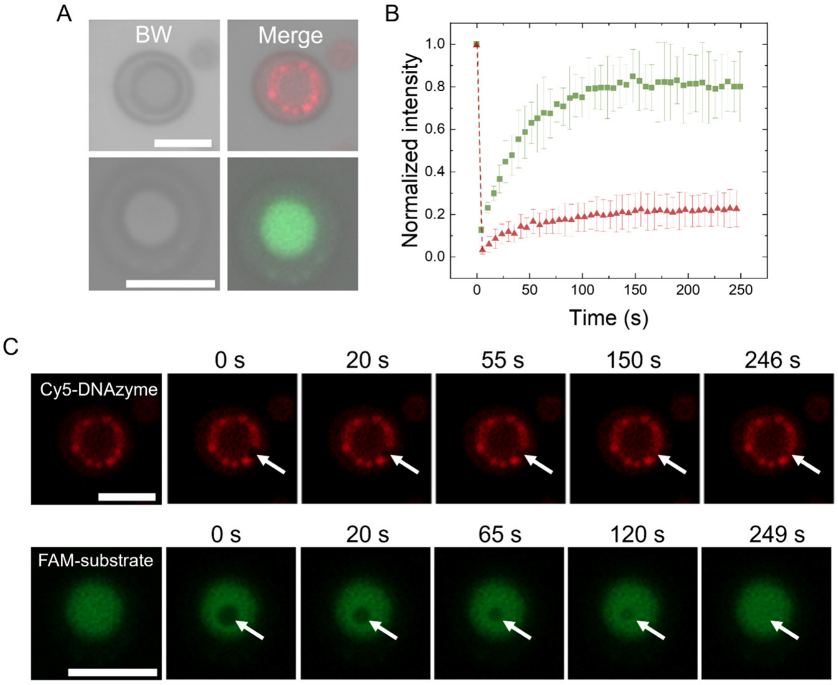


**Supplementary Fig. 38.** Intracondensate mobility of DNAzyme and RNA substrate. **(A)** Spatial distribution of Cy5-labeled DNAzyme and FAM-labeled RNA substrate within multiphase condensates. **(B)** Fluorescence recovery after photobleaching (FRAP) kinetics for the DNAzyme (red) and RNA substrate (green). **(c)** Time-lapse sequence of the fluorescence recovery process. Scale bar: 5 μm.

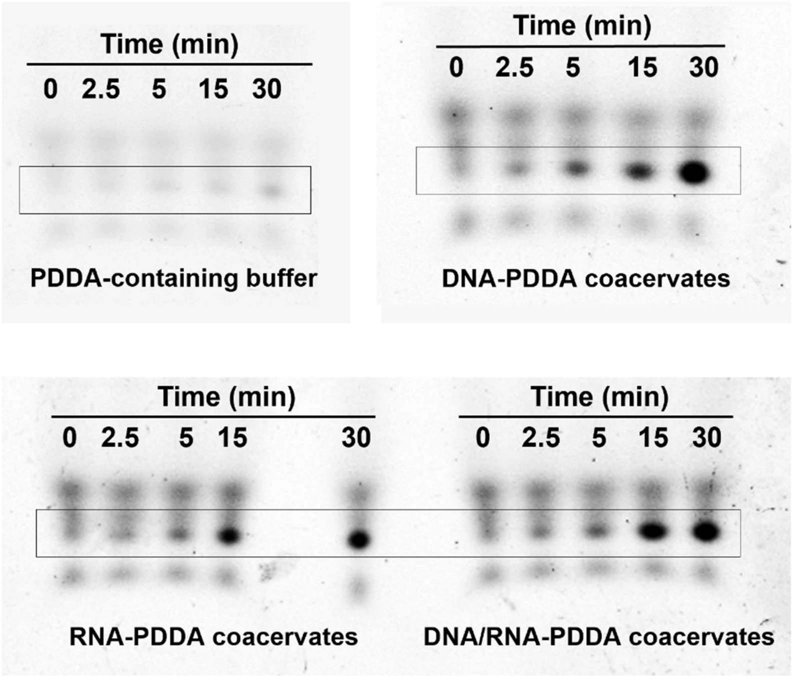


**Supplementary Fig. 39.** Gel electrophoresis of DNAzyme cleavage reactions within different coacervates. The rectangular box indicates the band for cleaved RNA.

**Supplementary Table 1.** Adenine fraction of selected oligonucleotide sequences utilized in this study. 'GAA' motifs on each sequence were blue colored (only DNA sequences are shown here).

| Name | Sequence (5'-3') | Adenine fraction (%) | Multiphase droplets? |
|---|---|---|---|
| TERRA_M2 | TTA GAA TTA GAA TTA GAA TTA GAA | 50 | Yes |
| PU22_M2 | TGA GAA TGA ATA GAA TGA ATA A | 54.5 | Yes |
| MT3_M2 | GAG AAA GAA AGA AAG AGA AA | 70 | Yes |
| TRF_M2 | CGA AAG AAC GAA GAG AAC | 55.6 | Yes |
| hTERC_M2 | GAA TTG CGA AGA ATG AAC CT | 40 | No |
| NRAS_M2 | GAA AGA AGC GAA TCT GAA | 50 | No |
| $A_{22}$ | $[AAA…AAA]_{22}$ | 100 | Yes |
| $A_{40}$ | $[AAA…AAA]_{40}$ | 100 | Yes |
| $T_{22}$ | $[TTT…TTT]_{22}$ | 0 | No |
| $T_{40}$ | $[TTT…TTT]_{40}$ | 0 | No |
| PU22_M2_N-to-A | AAA GAA AGA AAA GAA AGA AAA A | 81.8 | Yes |
| PU22_M2_N-to-U/T | TTT GTT TGT TTT GTT TGT TTT T | 0 | No |
| TERRA_M2_N-to-A | AAA GAA AAA GAA AAA GAA AAA GAA | 83.3 | Yes |
| TERRA_M2_N-to-U/T | TTT GTT TTT GTT TTT GTT TTT GTT | 0 | No |

**Supplementary Table 2.** Critical salt concentration (CSC) for dissolving DNA-PLL and RNA-PLL droplets. All test was performed under the charge concentration of [e+] = [e-] = 1mM.

| Name | Sequence (5'-3') | Droplets | CSC (mM) | Multiphase droplets? |
|---|---|---|---|---|
| TERRA_M2 | UUA GAA UUA GAA UUA GAA UUA GAA | RNA-PLL | 900 | Yes |
| | TTA GAA TTA GAA TTA GAA TTA GAA | DNA-PLL | 700 | |
| MT3_M2 | GAG AAA GAA AGA AAG AGA AA | RNA-PLL | 1100 | Yes |
| | GAG AAA GAA AGA AAG AGA AA | DNA-PLL | 700 | |
| PU22_M2 | UGA GAA UGA AUA GAA UGA AUA A | RNA-PLL | 900 | Yes |
| | TGA GAA TGA ATA GAA TGA ATA A | DNA-PLL | 600 | |
| TRF2_M2 | CGA AAG AAC GAA GAG AAC | RNA-PLL | 800 | Yes |
| | CGA AAG AAC GAA GAG AAC | DNA-PLL | 500 | |
| NRAS_M2 | GAA AGA AGC GAA UCU GAA | RNA-PLL | 900 | Yes |
| | GAA AGA AGC GAA TCT GAA | DNA-PLL | 700 | |
| hTERC_M2 | GAA UUG CGA AGA AUG AAC CU | RNA-PLL | 700 | No |
| | GAA TTG CGA AGA ATG AAC CT | DNA-PLL | 500 | |
| $A_{22}$ | $[rArArA\ldots rArArA]_{22}$ | RNA-PLL | 900 | Yes |
| | $[AAA\ldots AAA]_{22}$ | DNA-PLL | 500 | |
| $U_{22}/T_{22}$ | $[UUU\ldots UUU]_{22}$ | RNA-PLL | 600 | No |
| | $[TTT\ldots TTT]_{22}$ | DNA-PLL | 500 | |

**Supplementary Table 3.** Sequence content of Seq 1 to Seq 10. 'GAA' motifs on each sequence were blue colored (only DNA sequences are shown here).

| Name | Sequence (5'-3') | Adenine fraction (%) | Multiphase droplets? |
|---|---|---|---|
| Seq 1 | AAA GAA TAA CAG ATG AAG AAA TCA | 62.5 | No |
| Seq 2 | ATA AAG AAG TAA GAC AAT AAG CAA | 62.5 | No |
| Seq 3 | AAG AGT AAA AGA ATA CAG AAA CTA | 62.5 | No |
| Seq 4 | AAT AAA GAG AAA TAG AAC AAC AAA | 70.8 | No |
| Seq 5 | AGA ATA ACA AAG AAG TAA GAA TGA | 62.5 | Yes |
| Seq 6 | ATC TTC ATT TCT CCA CTC TCT CTC | 12.5 | No |
| Seq 7 | ATC TCT CGT CTC CTT CTT CCC TTC | 4.17 | No |
| Seq 8 | ATC TTC AGT TCT CCA CTC TCT CTC | 12.5 | No |
| Seq 9 | TCT TAT CGT CCT CTT CTC CTA CCC | 8.33 | No |
| Seq 10 | TCA TAC TGT TCT TCA TCT CCT TCC | 12.5 | No |

**Supplementary Table 4.** Sequences containing only G and A residues form multiphase condensates but have no base pairing propensity.

| Name | Sequence (5'-3') | U/T fraction (%) | Multiphase droplets? |
|---|---|---|---|
| MT3_M2 | GAG AAA GAA AGA AAG AGA AA | 0 | Yes |
| TRF_M2 | CGA AAG AAC GAA GAG AAC | 0 | Yes |
| $A_{22}$ | $[AAA…AAA]_{22}$ | 0 | Yes |
| $A_{40}$ | $[AAA…AAA]_{40}$ | 0 | Yes |
| PU22_M2_N-to-A | AAA GAA AGA AAA GAA AGA AAA A | 0 | Yes |
| TERRA_M2_N-to-A | AAA GAA AAA GAA AAA GAA AAA GAA | 0 | Yes |
| $[AAG]_8$ | AAG AAG AAG AAG AAG AAG AAG AAG | 0 | Yes |
| $[AGA]_8$ | AGA AGA AGA AGA AGA AGA AGA AGA | 0 | Yes |
| $[GAA]_8$ | GAA GAA GAA GAA GAA GAA GAA GAA | 0 | Yes |
| $[GAA]_7$ | GAA GAA GAA GAA GAA GAA GAA | 0 | Yes |
| $[GAA]_7$ mut1 | AGAA GAA AGAA GAA AGAA GAA GAA | 0 | Yes |
| $[GAA]_7$ mut2 | GAA GAA GAA GAA A GAA GAA GAA | 0 | Yes |
| $[GAA]_7$ mut3 | AGAA GAA GAA GAA GAA GAA GAA | 0 | Yes |
| $[CGCGAA]_4$ | CGCGAA CGCGAA CGCGAA CGCGAA | 0 | Yes |
| $[AACGAA]_4$ | AACGAA AACGAA AACGAA AACGAA | 0 | Yes |
| $[GACGAA]_4$ | GACGAA GACGAA GACGAA GACGAA | 0 | Yes |
| $[AAAGAA]_4$ | AAAGAA AAAGAA AAAGAA AAAGAA | 0 | Yes |
| $[AGAGAA]_4$ | AGAGAA AGAGAA AGAGAA AGAGAA | 0 | Yes |
| $[ACAGAA]_4$ | ACAGAA ACAGAA ACAGAA ACAGAA | 0 | Yes |
| $[GCAGAA]_4$ | GCAGAA GCAGAA GCAGAA GCAGAA | 0 | Yes |

**Supplementary Table 5.** Critical salt concentration (CSC) for dissolving DNA-PLL and RNA-PLL droplets formed by PU22_M2 N-to-U, TERRA_M2_N-to-U, Seq 1 and Seq 2, all of which cannot form multiphase condensates. All tests were performed under the charge concentration of [e+] = [e-] = 1mM. These results indicate that while a substantial difference in critical salt concentration between RNA-PLL and DNA-PLL coacervates appears to be a necessary condition for the formation of multiphase condensates, it is not sufficient by itself. Specifically, we observed that some DNA/RNA sequence pairs, such as Seq 1 and Seq 2, exhibit large differences in critical salt concentration yet do not form multiphase droplets.

| Name | Sequence (5'-3') | Droplets | CSC (mM) | Multiphase droplets? |
|---|---|---|---|---|
| PU22_M2_N-to-U | UUU GUU UGU UUU GUU UGU UUU U<br>TTT GTT TGT TTT GTT TGT TTT T | RNA-PLL<br>DNA-PLL | 700<br>500 | No |
| TERRA_M2_N-to-U | UUU GUU UUU GUU UUU GUU UUU GUU<br>TTT GTT TTT GTT TTT GTT TTT GTT | RNA-PLL<br>DNA-PLL | 700<br>500 | No |
| Seq 1 | AAA GAA UAA CAG AUG AAG AAA UCA<br>AAA GAA TAA CAG ATG AAG AAA TCA | RNA-PLL<br>DNA-PLL | 900<br>600 | No |
| Seq 2 | AUA AAG AAG UAA GAC AAU AAG CAA<br>ATA AAG AAG TAA GAC AAT AAG CAA | RNA-PLL<br>DNA-PLL | 1000<br>600 | No |

**Supplementary Table 6.** Five candidate models were compared with the same family-grouped cross-validation procedure. Point estimates were transcribed from the final model comparison. The Physics + sequence grammar RF was selected for candidate-sequence screening.

| Model | Balanced accuracy | ROC-AUC |
|---|---|---|
| Physics-only RF | 0.808 | 0.865 |
| Physics + sticker continuity RF | 0.812 | 0.842 |
| Physics + sequence grammar RF | 0.834 | 0.867 |
| Physics + CNN features RF | 0.787 | 0.845 |
| CNN | 0.724 | 0.848 |

**Supplementary Table 7.** Diblock DNA and RNA oligonucleotides from S1, S2, and S3 motifs.

| Name | Sequence (5'-3') |
|---|---|
| d(S1-b-S2) | TAA TAC GAC TCA CTA TAG TGA GAA TGA ATA GAA TGA ATA A |
| d(S1-b-S3) | TAA TAC GAC TCA CTA TAG TTA GAA TTA GAA TTA GAA TTA GAA |
| d(S2-b-S3) | TGA GAA TGA ATA GAA TGA ATA A TTA GAA TTA GAA TTA GAA TTA GAA |
| r(S2-b-S3) | UGA GAA UGA AUA GAA UGA AUA A UUA GAA UUA GAA UUA GAA UUA GAA |

**Supplementary Table 8.** Sequences used for RNA partitioning and cleavage assays.

| Name | Sequence (5'-3') |
|---|---|
| DNAzyme | TCA TGA GGC TAG CTA CAA CGA GGT TAG |
| RNA substrate | CUA ACC GUC AUG A |
| RNA substrate 1 | CUA ACCG |
| RNA substrate 2 | UCA UGA |
| Broccoli | UG AGA CGG UCG GGU CCA GAU AUU CGU AUC UGU CGA GUA GAG UGU GGG CUC A |

**Supplementary Table 9.** Oligonucleotide sequence utilized in this study (only DNA sequences are shown here).

| Number | Name | Sequence (5'-3') | Multiphase droplets? |
|---|---|---|---|
| **1** | TERRA_M2 | TTA GAA TTA GAA TTA GAA TTA GAA | Yes |
| **2** | [TTAGAA]1 | TTA GAA (as the Name implies) | No |
| **3** | [TTAGAA]2 | TTA GAA TTA GAA (as the Name implies) | No |
| **4** | [TTAGAA]8 | as the Name implies | Yes |
| **5** | [TTAGAA]16 | as the Name implies | Yes |
| **6** | TERRA_M2_mut1 | AAT AAT AAT AAT TAG TAG TAG TAG | Yes |
| **7** | TERRA_M2_mut3 | AAA TTG AAA GTT GTT AAA TTG AAA | Yes |
| **8** | TERRA_M2_mut4 | AAG ATT AAG ATT ATT AAG TTA GAA | Yes |
| **9** | PU22_M2 | TGA GAA TGA ATA GAA TGA ATA A | Yes |
| **10** | MT3_M2 | GAG AAA GAA AGA AAG AGA AA | Yes |
| **11** | TRF_M2 | CGA AAG AAC GAA GAG AAC | Yes |
| **12** | hTERC_M2 | GAA TTG CGA AGA ATG AAC CT | No |
| **13** | NARS_M2 | GAA AGA AGC GAA TCT GAA | No |
| **14** | $A_{22}$ | $[AAA\ldots AAA]_{22}$ | Yes |
| **15** | $A_{40}$ | $[AAA\ldots AAA]_{40}$ | Yes |
| **16** | $T_{22}$ | $[TTT\ldots TTT]_{22}$ | No |
| **17** | $T_{40}$ | $[TTT\ldots TTT]_{40}$ | No |
| **18** | PU22_M2_N-to-A | AAA GAA AGA AAA GAA AGA AAA A | Yes |
| **19** | PU22_M2_N-to-T/U | TTT GTT TGT TTT GTT TGT TTT T | No |
| **20** | TERRA_M2_N-to-A | AAA GAA AAA GAA AAA GAA AAA GAA | Yes |
| **21** | TERRA_M2_N-to-T/U | TTT GTT TTT GTT TTT GTT TTT GTT | No |
| **22** | Seq 1 | AAA GAA TAA CAG ATG AAG AAA TCA | No |
| **23** | Seq 2 | ATA AAG AAG TAA GAC AAT AAG CAA | No |
| **24** | Seq 3 | AAG AGT AAA AGA ATA CAG AAA CTA | No |
| **25** | Seq 4 | AAT AAA GAG AAA TAG AAC AAC AAA | No |
| **26** | Seq 5 | AGA ATA ACA AAG AAG TAA GAA TGA | Yes |
| **27** | Seq 6 | ATC TTC ATT TCT CCA CTC TCT CTC | No |
| **28** | Seq 7 | ATC TCT CGT CTC CTT CTT CCC TTC | No |
| **29** | Seq 8 | ATC TTC AGT TCT CCA CTC TCT CTC | No |
| **30** | Seq 9 | TCT TAT CGT CCT CTT CTC CTA CCC | No |
| **31** | Seq 10 | TCA TAC TGT TCT TCA TCT CCT TCC | No |
| **32** | [AAG]8 | as the Name implies | Yes |
| **33** | [AGA]8 | as the Name implies | Yes |
| **34** | [GAA]8 | as the Name implies | Precipitates |
| **35** | [AAT]8 | as the Name implies | No |
| **36** | [ATA]8 | as the Name implies | No |
| **37** | [TAA]8 | as the Name implies | No |
| **38** | [AAC]8 | as the Name implies | No |
| **39** | [ACA]8 | as the Name implies | No |
| **40** | [CAA]8 | as the Name implies | No |
| **41** | PU22_M1 | TGA GGA TGG ATA GGA TGG ATA A | Yes |
| **42** | [CGAA]6 | as the Name implies | Yes |
| **43** | [AGAA]6 | as the Name implies | Precipitates |
| **44** | [TGAA]6 | as the Name implies | Precipitates |

| | | | |
|---|---|---|---|
| **45** | [TCGAA]5 | as the Name implies | Yes |
| **46** | [TAGAA]5 | as the Name implies | Yes |
| **47** | [ATGAA]5 | as the Name implies | Yes |
| **48** | [ACGAA]5 | as the Name implies | Yes |
| **49** | [AAGAA]5 | as the Name implies | Yes |
| **50** | [GCGAA]5 | as the Name implies | Yes |
| **51** | [TTGAA]5 | as the Name implies | Precipitates |
| **52** | [CCGAA]5 | as the Name implies | Precipitates |
| **53** | [GTGAA]5 | as the Name implies | Precipitates |
| **54** | [GAGAA]5 | as the Name implies | Precipitates |
| **55** | [CTGAA]5 | as the Name implies | No |
| **56** | [CAGAA]5 | as the Name implies | No |
| **57** | [TCTGAA]4 | as the Name implies | Yes |
| **58** | [TATGAA]4 | as the Name implies | Yes |
| **59** | [TGTGAA]4 | as the Name implies | Yes |
| **60** | [TACGAA]4 | as the Name implies | Yes |
| **61** | [TTAGAA]4 | as the Name implies | Yes |
| **62** | [TAAGAA]4 | as the Name implies | Yes |
| **63** | [CGCGAA]4 | as the Name implies | Yes |
| **64** | [CTAGAA]4 | as the Name implies | Yes |
| **65** | [AATGAA]4 | as the Name implies | Yes |
| **66** | [AGTGAA]4 | as the Name implies | Yes |
| **67** | [AACGAA]4 | as the Name implies | Yes |
| **68** | [ATAGAA]4 | as the Name implies | Yes |
| **69** | [ACAGAA]4 | as the Name implies | Yes |
| **70** | [AAAGAA]4 | as the Name implies | Yes |
| **71** | [AGAGAA]4 | as the Name implies | Yes |
| **72** | [GTTGAA]4 | as the Name implies | Yes |
| **73** | [GACGAA]4 | as the Name implies | Yes |
| **74** | [GTAGAA]4 | as the Name implies | Yes |
| **75** | [GCAGAA]4 | as the Name implies | Yes |
| **76** | [TTTGAA]4 | as the Name implies | Precipitates |
| **77** | [TGAGAA]4 | as the Name implies | Precipitates |
| **78** | [CGTGAA]4 | as the Name implies | Precipitates |
| **79** | [ATTGAA]4 | as the Name implies | Precipitates |
| **80** | [ATCGAA]4 | as the Name implies | Precipitates |
| **81** | [GCTGAA]4 | as the Name implies | Precipitates |
| **82** | [GATGAA]4 | as the Name implies | Precipitates |
| **83** | [GTCGAA]4 | as the Name implies | Precipitates |
| **84** | [GCCGAA]4 | as the Name implies | Precipitates |
| **85** | [GAAGAA]4 | as the Name implies | Precipitates |
| **86** | [TCCGAA]4 | as the Name implies | No |
| **87** | [TGCGAA]4 | as the Name implies | No |
| **88** | [TCAGAA]4 | as the Name implies | No |
| **89** | [CTTGAA]4 | as the Name implies | No |
| **90** | [CCTGAA]4 | as the Name implies | No |
| **91** | [CATGAA]4 | as the Name implies | No |
| **92** | [CTCGAA]4 | as the Name implies | No |
| **93** | [CACGAA]4 | as the Name implies | No |

| | | | |
|---|---|---|---|
| **94** | [CCAGAA]4 | as the Name implies | No |
| **95** | [CAAGAA]4 | as the Name implies | No |
| **96** | [CGAGAA]4 | as the Name implies | No |
| **97** | [ACTGAA]4 | as the Name implies | No |
| **98** | [ACCGAA]4 | as the Name implies | No |
| **99** | [AGCGAA]4 | as the Name implies | No |
| **100** | [GAA]7 | as the Name implies | Yes |
| **101** | [GAA]7 mut 1 | AGAA GAA GAA GAA GAA GAA GAA | Yes |
| **102** | [GAA]7 mut 2 | GAA GAA GAA GAA A GAA GAA GAA | Yes |
| **103** | [GAA]7 mut 3 | AGAA GAA A GAA GAA A GAA GAA GAA | Yes |
| **104** | Seg 1 | ACGCGCGAACTGAAACTGAACGA | Yes |
| **105** | Seg 2 | GAGAACGGAAAGTAGTAGAATAC | Yes |
| **106** | Seg 3 | TGCGAATAATCGAACAGAATC | No |
| **107** | Seg 4 | GAGAACATGAACAGGAAATAGAAC | Yes |
| **108** | Seg 5 | TAGGAATGAAAGTGAAAGAACAT | Yes |
| **109** | Seg 6 | ACTGAAATGGAATCTGAAGAGAATG | No |
| **110** | hTERC_M2_ mut | AGAA TTG CGA AAGA ATG AAC CT | Yes |
| **111** | NARS_M2_ mut | AGAA A GAA GC GAA TCT GAAA | Yes |
| **112** | hTERC_M2_ mut 2 | AGAA TTG CGA AGA ATG AAC CT | No |
| **113** | hTERC_M2_ mut 3 | GAA TTG CGA AAGA ATG AAC CT | No |
| **114** | NARS_M2_ mut 2 | GAA A GAA GC GAA TCT GAAAA | No |
| **115** | Promotor (S1) | TAA TAC GAC TCA CTA TAG | No |

**Supplementary Table 10.** The specific experimental conditions under which the condensates shown in each figure were assembled.

| Data in Figures | [DNA] (by charge) | [RNA] (by charge) | [PLL] (by charge) | [KCl] |
|---|---|---|---|---|
| Figure 1(D, F, H) | 2.5 mM | 2.5 mM | 5 mM | 500 mM |
| Figure 2A | 2.5 mM | 2.5 mM | 5 mM | 100 to 900 mM as indicated in the figure |
| Figure 2F | 4 mM, 2.5 mM and 1.5 mM as indicated in the figure | 1 mM, 2.5 mM and 3.5 mM as indicated in the figure | 5 mM | 500 mM |
| Figure 3A to 3E | 2.5 mM | 2.5 mM | 5 mM | 500 mM |
| Figure 4B, 4C, 4D | 2.5 mM | 2.5 mM | 5 mM | 500 mM |
| Figure 4F | 2.5 mM | 2.5 mM | 5 mM | 150 mM and 500 mM as indicated in the figure caption |
| Figure 5B | 2.5 mM | 2.5 mM | 5 mM | 500 mM |
| Figure 5D, G | 2.5 mM | 2.5 mM | 5 mM | 150 mM |

**Supplementary Video 1**

Assembly of multiphase droplets upon addition of poly-L-lysine (PLL) to a solution containing DNA and RNA.

**Supplementary Video 2**

Evaporation-induced solid-to-liquid transition of DNA/RNA multiphase droplets.

**Supplementary Video 3**

Reversal of evaporation-induced transition: addition of water to the evaporated sample restores multiphase droplet formation.
